# Safe Redosable Low-Immunogenic In Vivo CAR-T Therapy for B Cell Malignancies and Solid Tumors

**DOI:** 10.64898/2026.06.30.735484

**Authors:** Ruquaiya Alam, Shantanu Kumar, Rohit Shukla, Nisha Chaudhary, Juli Gupta, Akshita Sinha, Rituparna Chaudhary, Mohan Ranganathan, Kashif Husain, Nisar Rahim Shaikh, Deeksha Joshi, Jahnvi Hora, Syed Ahmad Ali, Subhankar Bose, Prasad Iyer, Irfan Ahmad Mir, Mohammad Husain, Vishnu Hari, Amit Kumar Srivastava, Ulaganathan Mabalirajan, Gaurav Kharya, Sivaprakash Ramalingam, Asimul Islam, Tanveer Ahmad

**Author notes:** **Correspondence:** Dr. Tanveer Ahmad; Dr. Asimul Islam. ^#^ Authors have equally contributed in the study.

## Abstract

In vivo CAR-T cell therapy eliminates manufacturing complexities associated with ex vivo autologous approaches, but safety concerns have limited adoption. We developed viroVbot, a next-generation in vivo CAR-T platform, by combining computational immunogenicity prediction (CIMMEX^TM^) with envelope engineering. Screening 22,562 glycoprotein sequences, we identified 641 vesiculovirus homologs, from which we selected Piry virus glycoprotein (PIRYV) as the optimal candidate. PIRYV exhibited lower MHC-epitope density, reduced human seroprevalence, with decreased T cell activation compared to VSV-G. To enhance targeting specificity, we engineered receptor-binding-deficient PIRYV (ePIRYV^RBD^) displaying CD3/CD7 nanobodies for T cell-selective transduction. To maximize safety, we engineered CAR-TRAP producer cells to eliminate unwanted B cell transduction and incorporated machine learning-optimized T cell-specific promoters that restrict CAR activation exclusively to lymphocytes. Additional modifications suppressed hepatocyte expression and prevented phagocytic uptake. In humanized xenograft models, viroVbot3 generated potent BCMA/CD19 specific CAR-T responses against multiple myeloma and Claudin18.2-targeting gastric cancer, demonstrating sequential redosing with alternative envelopes. Critically, viroVbot3 exhibited minimal off-target organ biodistribution with CAR expression restricted to T lymphocytes. These findings establish viroVbot as a low-immunogenic platform for scalable in vivo CAR-T manufacturing with capability for sequential redosing across hematologic and solid tumors.

## Introduction

CAR-T cell therapy has transformed the treatment of B cell malignancies and, more recently, autoimmune diseases, with several products approved for relapsed and refractory leukemias, lymphomas, and multiple myeloma.^1–4^ However, only a small fraction of eligible patients have received this therapy, with the cumulative number of patients treated globally reaching only an estimated < 70,000 by 2025 since the first two products were approved in 2017.^5,6^ This limited access reveals the complexity of the autologous manufacturing process, which requires apheresis, shipment of cells to specialized facilities, viral transduction and expansion under GMP conditions, quality release testing, and re-infusion after lymphodepleting chemotherapy.^7^ The resulting long vein-to-vein times, high costs, conditioning-related toxicities, and dependence on specialized infrastructure limit patient access, particularly in regions without established cell therapy centers.^8,9^

To address these limitations, in vivo CAR-T generation has emerged as a promising alternative in which CAR transgenes are delivered directly to endogenous T cells, removing the need for apheresis, ex vivo manipulation, and lymphodepleting chemotherapy.^10–12^ Lentiviral vectors (LVVs) are the gold standard for this approach because they support stable, integration-dependent CAR expression, accommodate large payloads, and have an established clinical safety profile.^13^ Several first-generation in vivo CAR-T platforms have entered clinical trials, including the ESO-T01 anti-BCMA vector, which produced stringent complete responses in relapsed or refractory multiple myeloma,^14^ and the KLN-1010 platform, which achieved MRD-negative responses in early-phase trials ^15^. In autoimmune indications, lipid nanoparticle (LNP)-based delivery of CD19 CAR mRNA has driven transient B cell depletion in refractory systemic lupus erythematosus.^16,17^ Nonetheless, in vivo CAR-T therapy still faces several technical and immunological challenges.

A primary concern is lentiviral envelope immunogenicity, particularly with vesicular stomatitis virus glycoprotein G (VSV-G), the standard envelope for ex vivo applications. VSV-G is rapidly inactivated by human serum complement and pre-existing or treatment-induced anti-envelope antibodies, compromising systemic delivery and preventing re-dosing.^18,19^ While directed evolution has generated serum-resistant VSV-G variants (S162T, T230N, T368A),^20^ heterologous vesiculovirus glycoproteins from Piry, Cocal, and Maraba viruses bypass anti-VSV-G immunity through low human seroprevalence.^19^ Cocal glycoprotein, now adopted in clinical-stage in vivo CAR-T platforms, efficiently transduces hematopoietic cells, but off-target remains a concern.^21,22^ Paramyxoviral glycoproteins from Nipah virus offer low seroprevalence and fusogenic flexibility for scFv-mediated retargeting; however, limited clinical experience, potential off-target hepatic sequestration, and unknown long-term immunogenicity profiles present developmental challenges.^23–25^ In contrast, measles virus glycoproteins, though offering attractive targeting properties, are constrained by nearly universal vaccine-induced anti-measles antibodies that substantially reduce transduction efficiency in the clinical setting.^26^ Recently explored alternatives such as dolphin morbillivirus (DMV) glycoproteins similarly face challenges with immunogenicity and off-target biodistribution,^27^ suggesting the need for an integrated envelope engineering strategy that addresses both viral immunogenicity and tissue-specific targeting simultaneously.

A second concern is inadvertent CAR display on vector particles. Anti-CD19 and anti-CD20 CARs retain antigen-binding activity, risking transduction of B cells including malignant clones.^28^ This risk is exemplified by a B-ALL relapse with CD19-negative leukemia expressing anti-CD19 CAR, caused by contaminating leukemic B cell transduction during manufacturing.^29^ The TRiP system suppresses CAR translation during production,^30^ but combined with fourth-generation modifications, still leaves residual surface CAR display and off-target transduction risk.^25^ A third challenge is off-target liver biodistribution. The low-density lipoprotein receptor (LDL-R), highly expressed on hepatocytes, drives accumulation of VSV-G- and Cocal-pseudotyped vectors and ectopic transgene expression.^31^ Kupffer/macrophage cell uptake further triggers innate immune activation and constrains CAR-T persistence.^32^ Current mitigation strategies include CD47 incorporation^11,32^, miR-122 hepatocyte silencing,^33,34^ and T cell-selective promoters that confine CAR expression to target cells while reducing tonic signaling and exhaustion.^35^ Despite these recent advances, no current platform integrates envelope safety, off-target CAR suppression, hepatic sequestration prevention, and transcriptional specificity within a single vector architecture.

In this study, we describe viroVbot, a next-generation in vivo CAR-T platform that integrates AI-driven design solutions for envelope immunogenicity, off-target CAR display, hepatic sequestration, and transcriptional specificity.

## Results

### CIMMEXA Identifies PIRYV as a Low-Immunogenic Envelope Candidate

We developed CIMMEXA, an AI-driven computational framework that combines large-scale sequence analysis with ensemble MHC class I epitope prediction to identify viral envelope proteins with intrinsically low immunogenicity (**Fig. 1a, Supplementary Fig. 1**). From 22,562 pH-dependent glycoprotein sequences in the family Rhabdoviridae, homology searches using VSV-G as the reference sequence and BLASTP, PSI-BLAST, and DELTA-BLAST identified 641 non-redundant VSV-G homologs after duplicate removal. These homologs encompassed substantial evolutionary diversity while preserving key functional features of VSV-G (**Table 1**). Immunogenicity profiling with CIMMEXA enabled systematic ranking of envelope proteins, identifying the PIRYV glycoprotein as a candidate with markedly reduced immunogenic potential (**Table 2**). Subsequently, we generated a full library of overlapping MHC class I and II peptides, which were stratified into strong binders, weak binders, and non-binders based on standardized binding thresholds using CIMMEXA. A weighted scoring algorithm was developed based on the distribution of strong and weak binders across both MHC classes (*see Methods*), enabling calculation of an aggregate immunogenicity burden/score for each viral envelope candidate (**Table 2**). Ranking against the VSV-G reference revealed conserved low-immunogenic regions within the 12 top-ranked envelope proteins. Notably, Vesicular stomatitis Alagoas virus envelope protein (VSAV-G) and Piry virus envelope protein (PIRYV) demonstrated substantially lower immunogenicity than the VSV-G (**Fig. 1b, c, Supplementary Fig. 2 & Table 3, 4**). This robust analysis established a set of naturally occurring viral envelope candidates with reduced MHC binding density, providing a strong foundation for subsequent mechanistic validation and translational development.

**Figure 1:**
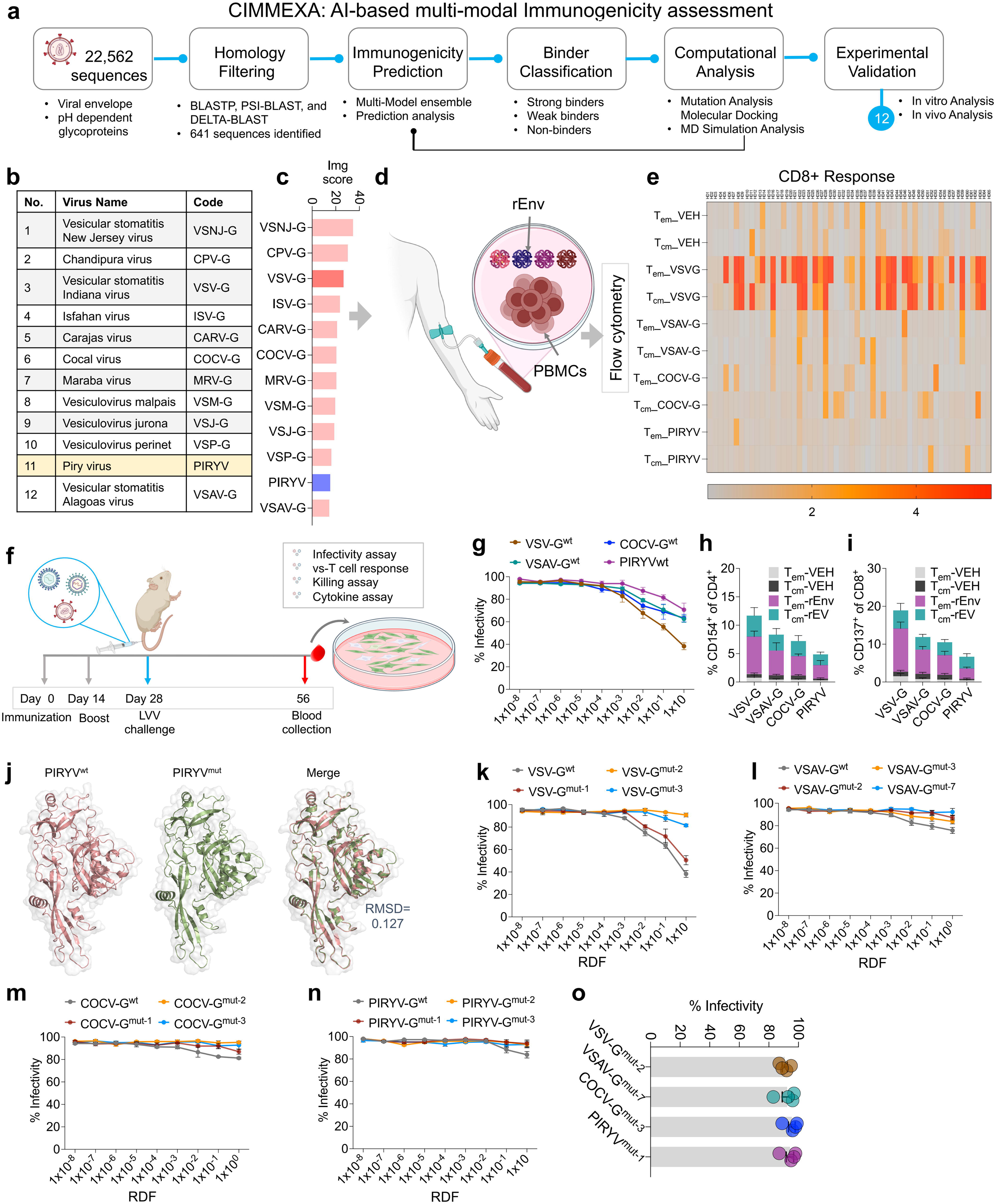
CIMMEXA identifies low-immunogenic vesiculovirus envelopes for in vivo CAR-T cell therapy. **a,** Schematic of CIMMEXA (Computational Immunogenicity Prediction and De-immunization), an AI-based multi-modal immunogenicity assessment platform integrating homology-based filtering, machine learning-based immunogenicity scoring, and comparative profiling. **b, c** Twelve representative vesiculovirus glycoproteins assessed by CIMMEXA, ranked by computational immunogenicity (Img) score. **d**, Schematic illustration showing blood collection, PBMCs isolation and incubation with respective recombinant envelope (rEnv) proteins. **e,** Heat map showing immune response to rEnv, showing CD8^+^ T cell responses quantified using costimulatory molecule expression (CD154/CD137) for T_em_ and T_cm_ subsets. VEH represents vehicle control treated samples. **f**, Schematics of immunization timeline in mice received subcutaneous immunization and boost injections of wild-type rEnv of VSV-G, VSAV-G, COCV-G, and PIRYV at day 0 and day 14, followed by lentiviral vector challenge at week 28, with blood collection and analysis at day21. **g**, Line graph showing neutralizing antibody response (as % reduction in infectivity) in HEK293T cells after incubation with various concentrations of sera from mice (n=12 mice per group) injected with various rEnv proteins. RDF = Reciprocal Dilution Factor. **h, i**, Bar graph of % CD154^+^ of CD4^+^ and % CD137^+^ of CD8^+^ T cell subsets (n=5; pooled from 12 animals). **j**, Structural comparison of wild-type (PIRYVwt) and mutant (PIRYVmut) envelopes showing overlay with root-mean-square deviation (RMSD) analysis (RMSD = 0.127 Å), indicating high structural similarity. **k-n**, Line graphs showing neutralizing antibody response (as % reduction in infectivity) in HEK293T cells after incubation with various concentrations of sera from mice injected with wild-type and computationally optimized mutant versions of rEnv proteins. RDF = Reciprocal Dilution Factor. **o**, % Infectivity of the final optimized mutant versions of VSV-G, VSAV-G, and COCV-G, selected from CIMMEXA analysis as candidates with the lowest immunogenicity while maintaining functional infectivity (n=12 mice per group).

To validate our computational predictions experimentally, we assessed pre-existing immune responses against these envelope glycoproteins in human subjects. Recombinant envelope proteins (rEnv) were generated for PIRYV, COCV-G, VSAV-G, and VSV-G and evaluated in 65 healthy volunteers to detect any pre-existing immune responses. Blood samples were analyzed using an ex vivo testing method with recombinant proteins for each envelope as inducers. Antigen-specific T cell responses were measured in both CD4^+^ and CD8^+^ memory subsets using costimulatory molecules CD154 and CD137 as activation readouts, following previously established protocols ^36^ (**Fig. 1d**). Validation of computational predictions revealed measurable antigen-specific CD4^+^ and CD8^+^ T cell responses following induction with VSV-G, while other envelope proteins elicited comparatively weaker immune responses, with PIRYV demonstrating the least immunogenicity among all tested envelope glycoproteins (**Fig. 1e and Supplementary Fig. 3**). Screening identified PIRYV as the envelope with the lowest pre-existing immune response, in line with the CIMMEXA prediction.

We subsequently evaluated all these envelope proteins in vivo for acute immune responses in BALB/c mice using a previously developed neutralization assay ^19^ (**Fig. 1f**). To ensure that host cell proteins, particularly MHC class I molecules, did not interfere with immunogenicity, we generated CRISPR-Cas9-mediated HEK293T producer cells with specific MHC-I knockout and thoroughly characterized these cells (**Supplementary Fig. 4a-f**). GFP-expressing lentiviral vectors with respective pseudotyped envelopes were used for transduction studies to determine the neutralizing antibody responses developed in mice. Among all tested envelopes, PIRYV consistently demonstrated the lowest neutralization compared to other envelope proteins, followed by COCV-G (**Fig. 1g**). In parallel, ex vivo virus-specific T cell assays revealed that PIRYV induced the least activation of envelope-specific CD4^+^ and CD8^+^ T cell populations (**Fig. 1h, i**). We also evaluated virus-specific CD8^+^ T cells for their cytotoxic capacity against cell lines expressing all these envelope proteins on their surface, respectively. Virus-specific CD8^+^ T cells from VSV-G-immunized animals showed the most robust cytotoxicity against VSV-G-expressing HEK293T cells compared to other targets (**Supplementary Fig. 4g**). Correspondingly, inflammatory cytokine levels including IFN-γ and TNF-α were highest in the VSV-G group and lowest in the PIRYV group (**Supplementary Fig. 4h, i**). These results establish PIRYV as a promising, low-immunogenicity envelope protein for further development.

### Computational optimization of envelope immunogenicity through targeted mutagenesis

To further optimize immunogenic profiles, we used CIMMEXA to pinpoint immunogenic residues within high-binding peptides and rationally designed amino acid substitutions to reduce predicted immunogenicity. Structural modeling confirmed that these mutations-maintained envelope stability and proper protein folding without compromising overall functionality (**Fig. 1j, Supplementary Fig. 5, & Table 5**).

Based on this strategy, the top 10 optimized candidates were selected alongside their respective natural counterparts and synthesized for in vitro functional evaluation. All engineered envelopes were assessed for their potential to mediate lentiviral transduction in HEK293T and Jurkat T cells using GFP as a reporter (**Supplementary Fig. 6**). High-efficiency variants were selected. The three top-performing envelope candidates were subsequently evaluated in vivo for acute immune responses through ex vivo infectivity assays. All mutated envelope proteins exhibited significantly reduced immunogenic response, including the optimized VSV-G variant (**Fig. 1k-o**). Computational mutagenesis effectively generated low-immunogenicity envelopes that preserve transduction capacity while minimizing immune activation.

### Structure-guided identification of PIRYV receptor interaction residues for receptor detargeting

We conducted structure-guided analysis of PIRYV to identify key residues involved in receptor binding, as this envelope demonstrated the lowest immunogenicity profile. First, we sought to determine whether PIRYV utilizes the conventional LDL-R or alternative receptors (LDL family receptors) for cellular entry, we generated HEK293T knockout cell lines targeting each of these receptors (**Supplementary Fig. 7a**). The results indicate that, similar to VSV-G and COCV-G, PIRYV also preferentially utilizes LDL-R compared to other potential receptors such as VLDL-R (very-low-density lipoprotein receptor) or LRP1 (LDL receptor-related protein 1). However, we found that PIRYV additionally utilizes LRP1 beyond LDL-R. In triple knockout HEK293T cells, PIRYV entry was completely abrogated, consistent with observations for VSV-G and COCV-G (**Supplementary Fig. 7b-d**).

Having established that PIRYV utilizes similar receptor proteins as other vesiculovirus family members, we proceeded to determine the specific residues in PIRYV that mediate interactions with LDLR. Structural analysis and comparative modeling with VSV-G and COCV-G identified four major residues in PIRVY: K182, Y207, Y352, and K354 (**Fig. 2a, b & Supplementary Fig. 8-10**). In silico analysis revealed that among these four residues of PIRYV, K182 and Y352, exhibited the most significant effects on receptor binding (**Supplementary Fig. 9**).

**Figure 2:**
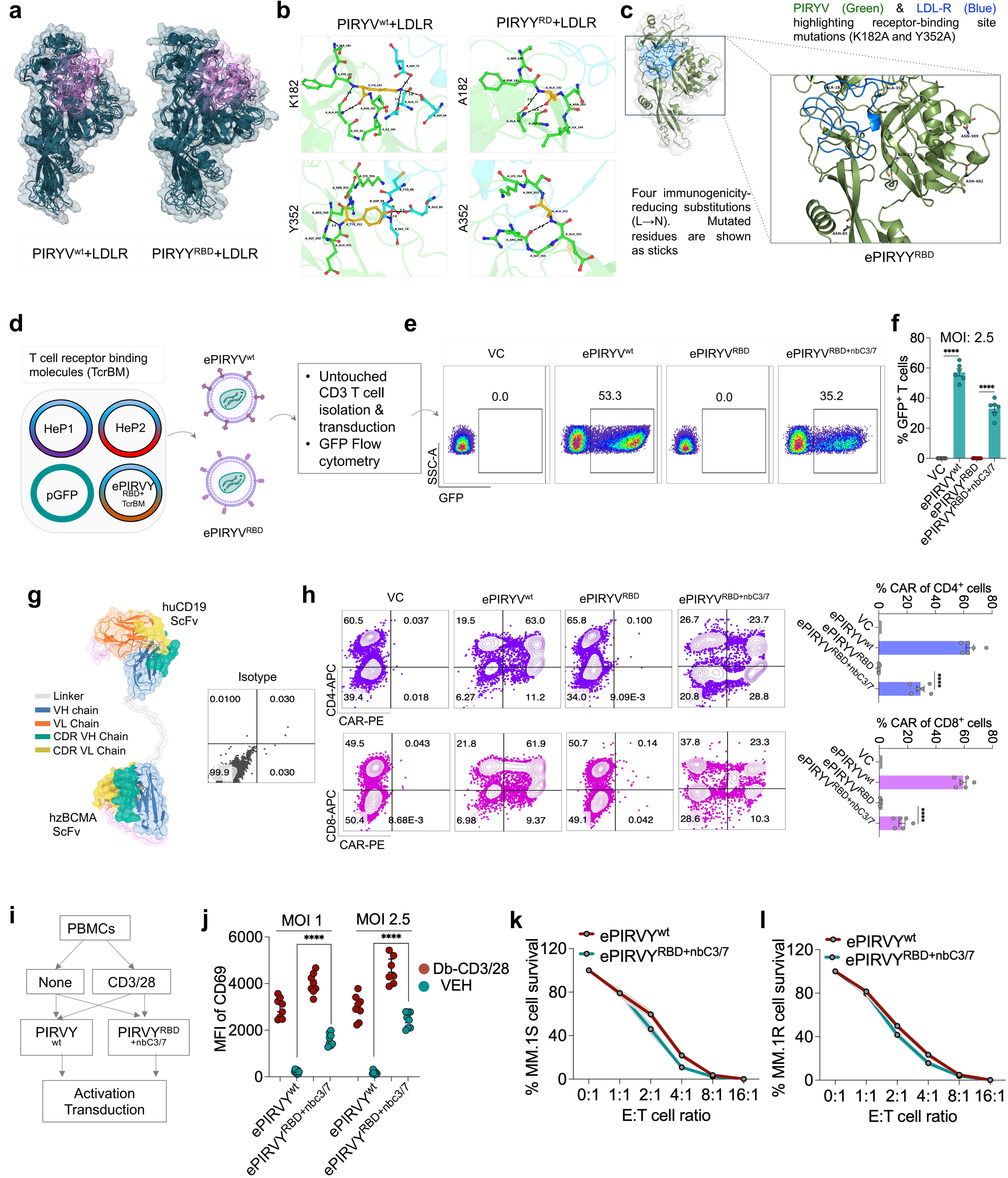
Engineering low-immunogenic envelope with T cell-targeted nanobodies and bispecific CAR construct. **a,** Structural models of receptor-binding-deficient PIRYV (ePIRYV^RBD^) engineered with K182A and Y352A mutations to eliminate low-density lipoprotein receptor (LDL-R) binding while preserving fusogenic capacity. Receptor-interacting region is highlighted in magenta **b,** Close up view of the two LDL-R contact residues, glycoprotein (green sticks) form hydrogen bond (dashed lines) with LDL-R (cyan) whereas in the mutant complex, substitutions abolish these bonds. **c,** Complete structural model of the engineered PIRYV^RBD^ glycoprotein in complex with LDL-R depicting ablation of LDL-R recognition, alongside the four immunogenicity-reducing residues. **d**, Schematic of lentiviral production and T cell isolation. **e,** Representative flow cytometry plots of transduction efficiency (%GFP^+^) in CD3^+^ T cells at MOI 2.5 with vector control (VC). ePIRYV^wt^. ePIRYV^RBD^ or ePIRYV^RBD+nbC3/7^. **f**, Bar graph showing the mean fluorescence intensity (MFI) of GFP^+^ cells. Representative of three independent experiments (n=6 biologically independent samples). **g,** Humanized scFv design for bispecific BCMA/CD19 CAR construct showing complementarity-determining regions (CDRs) in heavy and light chains. **h,** Representative contour plots showing comparative CAR expression (PE) in transduced CD4^+^ (APC, top) and CD8^+^ (APC, bottom) T cell subsets in activated primary CD3^+^ T cells from healthy donors (n = 6). Bar graph quantifies the percentage of CAR^+^ cells (using an anti-G4S linker antibody) among CD4^+^ and CD8^+^ T cells. **i,** PBMCs were either left unstimulated (None) or activated with anti-CD3/CD28 (CD3/CD28) Dynabeads before transduction with PIRVYwt or ePIRYRBD^+nbC3/7^, followed by analysis of activation. **j,** T cell activation markers (CD69) measured as MFI following transduction with PIRYV^wt^ or ePIRYV^RBD+nbC3/7^ at MOI 1 and MOI2.5 after 24 h post-transduction. Each point represents one donor (n = 8). **k, l,** In vitro cytotoxicity of CAR-T cells generated with PIRY^wt^ or ePIRY^RBD+nbC3/7^ against BCMA^+^ multiple myeloma targets MM.1S (k) and MM.1R (l), expressed as percentage target-cell survival across the indicated effector-to-target (E:T) ratios after 24 h coculture. Data represents mean ± SEM. ****p < 0.0001. A non-parametric t-test was used for statistical analysis with n = 3 biologically independent samples). *See Supplementary* Figures 7-11 *for extended nanobody characterization and CAR construct optimization*.

To validate these findings experimentally, 15 combinatorial variants were generated and evaluated them in vitro using the transduction efficiency assay. Similar to in silico results, the quadruple mutant (K182A+Y207A+Y352A+K354A) demonstrated nearly complete abrogation of receptor binding activity. Notably, the two key residues (K182A+Y352A) were sufficient to effectively eliminate receptor binding (**Supplementary Fig. 11a-c**). This double mutant was further validated across multiple cell lines, including Jurkat T cells and healthy donor-derived CD3^+^ T cells, consistently showing complete loss of receptor binding (**Supplementary Fig. 11d, e**). These results thus establish an efficient strategy for receptor detargeting while minimizing the number of required mutations.

Integrating these in silico, in vitro, and in vivo results, we generated a fully optimized PIRYV envelope with reduced immunogenicity and receptor binding deficiency (referred as; ePIRYV^RBD^; RBD: receptor binding deficient) for in vivo CAR-T cell generation (**Fig. 2c**). To create an effective T cell targeting system, we developed T cell receptor-specific humanized nanobodies using our proprietary de novo nanobody generation pipeline (CelNFo) against CD3, CD4, CD5, CD7, CD8, and TCRα (**Supplementary Fig. 12a, b 7** (**Table 6**), along with established T cell ligands CD80 and CD252.

All nanobodies were synthesized with myc tags for identification, and evaluated for surface expression in HEK293T producer cells by transient transfection which showed surface expression (**Supplementary Fig. 12c**). Subsequently, we generated lentiviral vectors encoding GFP reporters using these engineered envelopes for T cell transduction studies. Among all tested ligands, CD3 and CD7 consistently demonstrated the highest transduction efficiency, and these two nanobodies were selected for further development. These T cell-targeting nanobodies mediated dose-dependent transduction in CD3^+^ T cells when incorporated with ePIRYV^RBD+nbC^^3/7^ (**Fig. 2d-f and Supplementary Fig. 12d**).

### Generation of Functional Bispecific BCMA/CD19 CAR-T Cells with viroVbots

Multiple myeloma was selected as the target indication for in vivo CAR-T cell development, based on our recent study in this area.^37^ Accordingly, we designed a bispecific CAR construct capable of simultaneously engaging BCMA and CD19, with a human CD19 scFv and a humanised BCMA as described by us previously.^37,38^ Clinical evidence further indicates that bispecific CARs co-targeting BCMA and CD19 yield favorable therapeutic outcomes,^39^ supporting the rationale for a dual-antigen targeting strategy. Computational analysis confirmed the binding affinity of human (hu)CD19 CAR to CD19 ectodomain (**Supplementary Fig. 13**). The final BCMA/CD19 bispecific CAR construct (designated bi-CAR; **Fig. 2g**) incorporated dual 4-1BB/ICOS co-stimulatory domains. The selection of the 4-1BB/ICOS co-stimulatory domains was based on our previous study.^38^ Upon evaluation of transduction efficiency, ePIRYV^RBD+nbC^^3/7^ demonstrated dose-dependent transduction efficiency (MOI 2.5) in CD3-activated T cells as well as in purified CD4^+^ and CD8^+^ T cell populations (**Fig. 2h & Supplementary Fig. 14a**).

To determine whether transduction efficiency was maintained in non-activated T cells, we compared the activation status of CD3/CD7-targeted lentiviral vectors with standard CD3/CD28 Dynabeads and found efficient transduction by ePIRYV^RBD+nbC^^3/7^ envelope in the absence of additional activation molecules (**Fig. 2i, j & Supplementary Fig. 14b**). Importantly, substantial transduction efficiency (>20%) was observed in patient-derived PBMCs (**Supplementary Fig. 14c, d**).

In vitro anti-tumor cytotoxicity assays demonstrated efficient activity against multiple target cell types, including MM1.S and MM.1R multiple myeloma cells, and CD19-expressing Raji lymphoma and NALM6 cells (**Fig. 2k, l and Supplementary Fig. 14e, f**). Functional characterization revealed sustained release of granzyme B and cytokines associated with CAR-T cell activity (**Supplementary Fig. 14g-i**). Importantly, T cell immunophenotyping confirmed preservation of memory T cell subsets (**Supplementary Fig. 14j**). These findings establish ePIRYV^RBD+nbC^^3/7^ as an efficient platform for generating T cell-targeted CAR with potent anti-tumor activity and preserved immunological function. We designated this optimized platform viroVbot and advanced to preclinical evaluation.

### Preclinical testing of viroVbots in humanized animal models

For preclinical validation, immunodeficient transgenic xenograft mouse model was engrafted with human hematopoietic stem cells (HSCs) following previously established protocols.^40^ The bi-CAR with ePIRYV^RBD+nbC^^3/7^ was used as the transgene delivery system (designated viroVbot1) and administered to mice at various concentrations, with the lowest concentration being 0.25 x10^6^ TU/mouse. Blood samples were collected at regular intervals, and CD3^+^ T cells were evaluated for CAR molecule expression by flow cytometry (**Fig. 3a**). ViroVbot1 demonstrated superior T cell transduction efficiency compared to ePIRYV^wt^ pseudotyped control lentiviral vectors expressing the same transgene across all analyzed tissue samples, including blood, bone marrow, and spleen respectively. (**Fig. 3b-d**).

**Figure 3:**
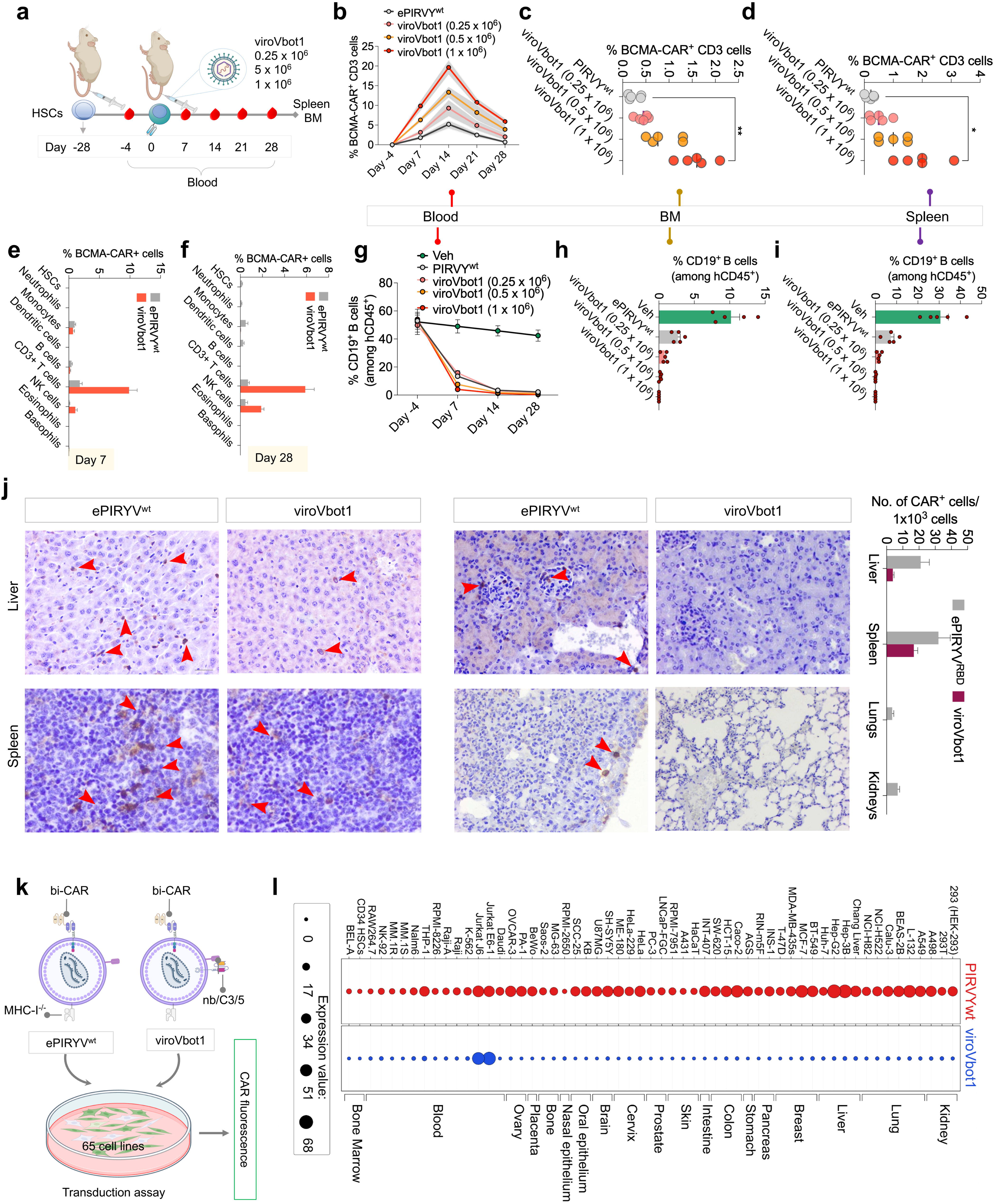
viroVbot1 achieves efficient in vivo T cell transduction with minimal off-target effects. **a,** Experimental timeline of in vivo CAR-T delivery in humanized xenograft model. NSG mice received intravenous engraftment of human hematopoietic stem cells (HSCs) on days - 28 with prior busulfan treatment. viroVbot1 (ePIRYV^RBD+nbC3/7^ encoding bi-CAR transgene) was administered intravenously at doses ranging from 0.25 × 10⁶ to 1 × 10⁶ TU/mouse on day 0. Blood, was collected at days -4, 7, 14, 21, and 28 post-vector administration and bone marrow (BM), and spleen at the end of the experiment. **b**, Kinetics of CAR-T cell expansion in peripheral blood. Percentage of BCMA-CAR^+^ CD3^+^ T cells over time (Day -4 to 28). **c**, **d,** CAR-T cell frequency in **(c)** bone marrow and **(d)** spleen at Day 28. Each point represents pooled samples from 15 animals (n = 5 from 3 pooled in each group). **e, f,** CAR-T cell transduction efficiency in peripheral blood cells determined by flow cytometry by collecting the samples at day 7 and day 28. **g**, Flow cytometry analysis of CD19^+^ cells among hCD45^+^cells showing rapid B cell depletion over time. **h, i,** Similarly in bone marrow and spleen. **j,** Representative IHC images of major organs (spleen, liver, lungs, and kidneys) showing CAR expression as indicated by red arrowheads with quantitative analysis (n=6 images); Bar graph shows CAR+ cells per 1×10^3^ cells by tissue in visceral tissues. **k,** Schematic of vector production of the two lentiviral vectors used: (left) ePIRY^wt^ with MHC-I^-/-^ modification and bispecific BCMA/CD19 CAR; (right) viroVbot1 with TcrBM-detargeted envelope (ePIRY^RBD+nbC3/7^), with the same bispecific CAR. Both vectors were used to transduce a panel of 65 distinct human-derived cell lines to assess CAR expression at MOI 2.5. **l,** CAR expression profile across 65 cell lines (ex vivo transduction assay). Dot plot showing CAR fluorescence (n=3 biologically independent samples). Data represents mean ± SEM. **p < 0.01; *p < 0.05. A non-parametric t-test was used for statistical analysis between groups. Scale bar; d: 200 μm.

To evaluate off-target effects of viroVbot1, CAR expression across all blood cell populations and major organs was assessed. As anticipated from the CD7 nanobody targeting, efficient bi-CAR expression was detected not only in T cells but also in NK cells, with lower, but detectable level expression in monocytes and B cells at early time points post-administration (day 7). At later time points, CAR expression in monocytes declined substantially and became undetectable in B cells, while expression in T and NK cells was maintained (**Fig. 3e, f**). Functional analysis revealed pronounced B cell depletion in the blood, spleen, and bone marrow, confirming the cytolytic activity of CAR-T cells generated in vivo by viroVbot1 (**Fig. 3g-i**).

Analysis of major organs by immunohistochemistry (IHC) revealed moderate CAR expression in the spleen, and liver. Importantly, weak signal was detected also in lungs and kidneys of mice treated with ePIRYV^wt^, whereas no signal was observed in viroVbot1 group **(Fig. 3j & Supplementary Fig. 15a**). Weak, residual signal was detected in the ePIRYV^wt^ in other organs, and again no signal was observed in viroVbot1 group. These IHC findings were further confirmed by RT-qPCR analysis of the vector copy number (VCN) in these tissues (**Supplementary Fig. 15b**). These results suggest that CAR proteins present on HEK293T cell surfaces during virus production may contribute to uptake by other immune cell populations, particularly B cells expressing CD19 receptors, potentially leading to undesired long-term effects. In parallel, uptake of lentiviral vectors by monocytes/macrophages and liver hepatocytes suggests these cells may serve as cellular sinks, limiting the availability of CAR carrying vectors in circulation for efficient T cell transduction and potentially causing unanticipated adverse effects. Similar phenomena have been documented by others research groups.^25^

To explicitly confirm that retargeted viroVbot1 specifically transduces T cells rather than other cell types, we evaluated these molecules in vitro using organ-specific cell lines (**Fig. 3k**). Evaluation of 65 different cell lines after 7 days of transduction revealed that only small fraction of specific cell lines were transduced, particularly B cells, monocytes, and macrophages with viroVbot1 (**Fig. 3l**). Given that liver accumulation represents a major safety concern for viral-based in vivo therapies, we subsequently focused on mitigating off-target uptake of viroVbots to enhance safety and targeting specificity.

### CAR-engineered producer cell lines mitigate B cell off-targeting of viroVbots

To minimize non-specific lentiviral vector uptake by B cells, macrophages, and hepatocytes, a systematic approach was implemented. First, we confirmed that viroVbot1 uptake by B cell lines expressing either CD19 or BCMA occurred through specific receptor-mediated mechanisms rather than passive uptake processes. We utilized both wild-type and CRISPR-Cas9 knockout versions of Raji cells and CD19/BCMA knockout MM.1S cells, which we had previously developed (**Fig. 4a)**.^41^

**Figure 4:**
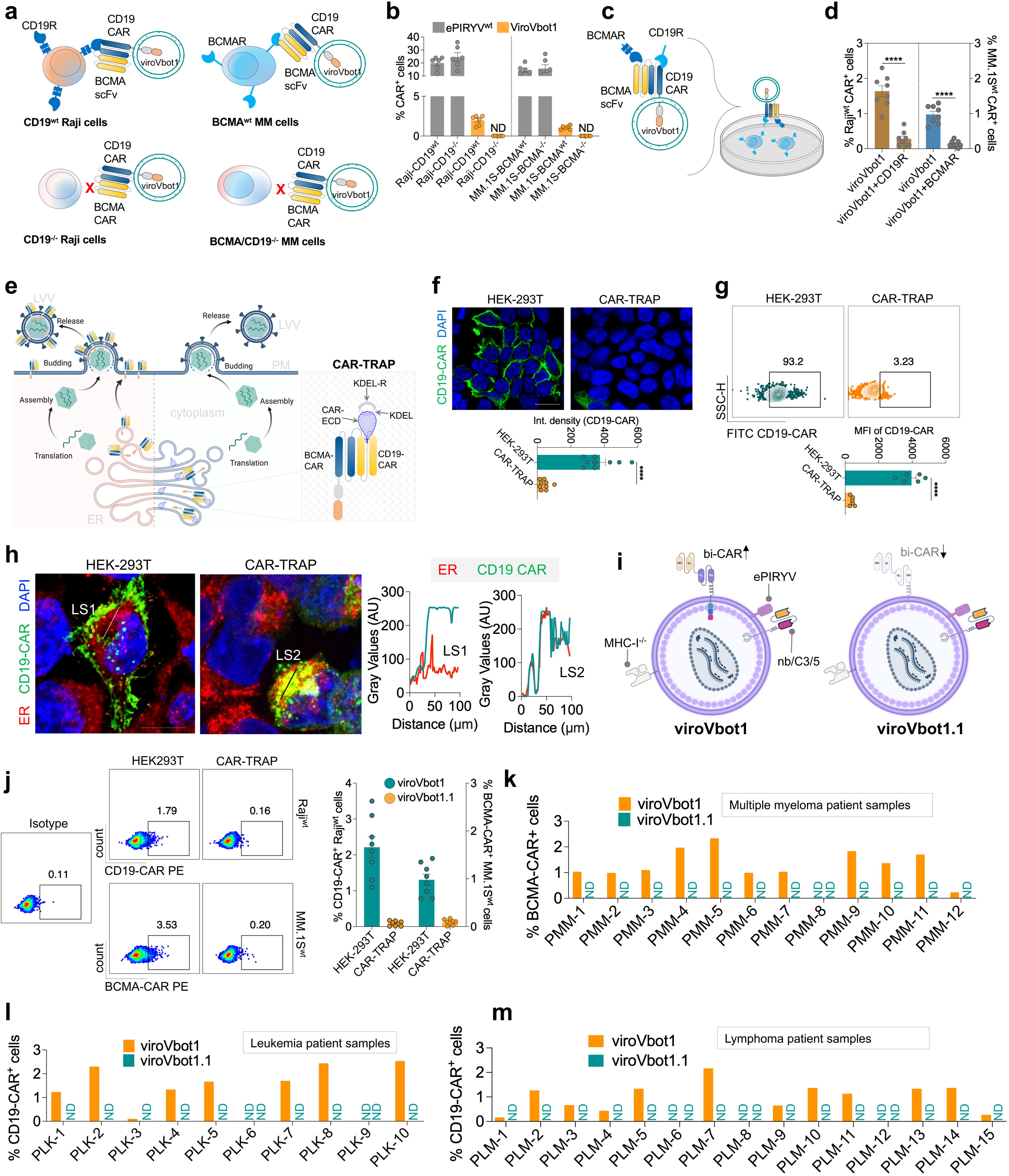
CAR-TRAP producer cells prevent off-target B cell transduction through ER retention. **a,** Schematic showing receptor-mediated CAR uptake by B cells with wild-type (CD19^wt^ Raji, BCMA^wt^ multiple myeloma (MM) phenotype and CRISPR-Cas9 knockout (BCMA/CD19^-/-^ MM, CD19^-/-^ Raji) phenotype. Cell lines were transduced with viroVbot1 or wild type PIRYV **b,** Flow cytometry showing ex vivo CAR transduction the wild type and knockout cells (n=6). **c,** Schematic of receptor competition assay showing that pre-incubation of viroVbot1 with recombinant BCMA (BCMAR) and CD19 (CD19R) extracellular domain proteins before transduction of wild-type Raji and MM.1S cells. **d,** Transduction efficiency (% CAR^+^ cells) in Raji and MM.1S cells, with viroVbot1 incubated with vehicle (PBS) or viroVbot1 incubated with CD19R or BCMAR (n=8 biologically independent samples). **e,** Schematic showing CAR glycoprotein displayed on wild-type producer cell line (left) is incorporated onto the lentiviral envelope in contrast to CAR-TRAP producer cell (right) where CD19-ECD-KDEL retains the CAR in ER lumen rather than plasma membrane. **f**, Representative Immunofluorescence image of HEK293T control cells (left), show robust CD19-CAR (green) throughout the cell surface; CAR-TRAP-expressing cells (right) show minimal surface CD19-CAR fluorescence. Bottom panel is quantification of CD19-CAR intensity density (integrated intensity per cell area) in HEK293T versus CAR-TRAP (n=9 cells). **g,** Contour plots of CAR (FITC-CD19) surface expression under non-permeabilising conditions with quantitative analysis shown as MFI (n=6), **h,** Representative Immunofluorescence image showing intracellular ER-localized CAR (anti-CD19-CAR, green) colocalized with calnexin (ER marker, red) in CAR-TRAP cells, along with the lines scans (LS) showing co-localisation of ER signal with CD19 CAR. **i,** Illustration of viroVbot1 particle produced from wild type producer cell and viroVbot1.1 particle produced from CAR-TRAP cells with bi-CAR as transgene. **j,** Dot plots showing % BCMA or % CD19 CAR expression in Raji^wt^ or MM.1S^wt^ cells with LVV obtained from HEK293T or CAR-TRAP cells. Quantification bar graphs (right) under same conditions (n= 8). **k,** BCMA CAR expression in patient multiple myeloma samples (PMM-1 to PMM-12) showing % of BCMA/CAR^+^ cells after transduction with viroVbot1 versus viroVbot1.1. **l-m,** Similarly, BCMA CAR expression in patient-derived leukemia (PLK-1 to PLK-10) and lymphoma (PLM-1 to PLM-15) samples. Data represents mean ± SEM. ****p < 0.0001. A non-parametric t-test was used for statistical analysis between groups. Scale bar; f: 50 μm, n: 10 μm.

Lentiviral vectors were generated with either ePIRYV^wt^ or viroVbot1 expressing bi-CAR transgenes and used to transduce B cells and MM.1S cells. As expected, modest uptake of viroVbot1 was observed in both CD19^wt^ and BCMA^wt^ cells, which was prevented in knockout cells (**Fig. 4b**). To further confirm that lentiviral vector uptake by receptor-expressing cells occurred via CAR-receptor-mediated processes, we pre-incubated viroVbot1 with recombinant BCMA and CD19 proteins before transducing wild-type receptor-expressing cells. This competition approach resulted in significant reduction of viroVbot1 uptake (**Fig. 4c, d**). These results confirm that B cell uptake of CAR transgene was not passive but actively mediated by interactions between CAR molecules and CD19 or BCMA receptors. This led us to develop a strategy for generating lentiviral vectors in producer cells without surface expression of CAR molecules.

To prevent BCMA/CD19 CAR protein expression on lentiviral vector surfaces, a previously developed strategy was used, which involves N-terminal tagging of the CD19 extracellular domain with an endoplasmic reticulum retention signal (KDEL). This approach involved stable expression of CD19 extracellular domain in producer cell lines with KDEL retention signals, ensuring that constitutively produced extracellular domain remained sequestered within the ER lumen (**Fig. 4e**). Surface CAR expression was analyzed by immunofluorescence and flow cytometry in HEK293T and CD19-ECD-KDEL (designated CAR-TRAP) cells (Fig. 4f, g). CD19-ECD-KDEL expression was subsequently assessed by immunofluorescence. No surface expression was detected under standard surface staining conditions, while expression was restricted to the ER under permeabilization conditions (**Fig. 4h**). These CAR-TRAP producer cells were subsequently used to generate bi-CAR expressing lentiviral vectors, designated viroVbot1.1 (**Fig. 4i**).

Testing in CD19-expressing B cells and BCMA-expressing multiple myeloma cells confirmed that KDEL-mediated retention effectively limited bi-CAR surface expression on producer cell lines, completely abrogating transduction of these CD19 and/or BCMA expressing target cells (**Fig. 4j**). Importantly, viroVbot1.1 vector showed no significant transduction when tested with patient-derived myeloma cells as well as in B cells from leukemia and lymphoma patients (**Figure 4k-m**). These results thus demonstrate that restricting CAR protein expression in producer cells effectively limits CAR surface display during LVV release, preventing detrimental B cell uptake.

### CD47 expression and liver-specific microRNA targeting attenuate viroVbot delivery to macrophages and hepatocytes

We employed a liver-specific microRNA-122 target sequence (5xmiR-122T) to promote transgene degradation in hepatocytes. The miR-122 target site sequence was expressed downstream of CD3ζ, with bi-CAR under EF1α promoter control (**Fig. 5a**). Expression levels remained unchanged in T cells between 5xmiR-122T-containing transgenes and vector control with non-targeting sequence (**Supplementary Fig. 16a**). Huh-7 hepatocytes showed markedly reduced bi-CAR expression (**Fig. 5b**). Primary hepatocytes exhibited reduced CAR expression, confirming hepatocyte-specific silencing (**Supplementary Fig. 16b**). 5xmiR-122T preserved transduction efficiency in primary T cells and anti-tumor activity against BCMA-expressing cells (**Supplementary Fig. 16c, d**). Since monocytes and macrophages also demonstrated lentiviral vector uptake, we developed additional strategies to limit expression in these cell types. A plasmid encoding CD47 with gp41 was used to stably introduce CD47 into CAR-TRAP producer cells, and surface expression was confirmed by flow cytometry and immunostaining (**Fig. 5c, d and Supplementary Fig. 17a, b**).

**Figure 5.**
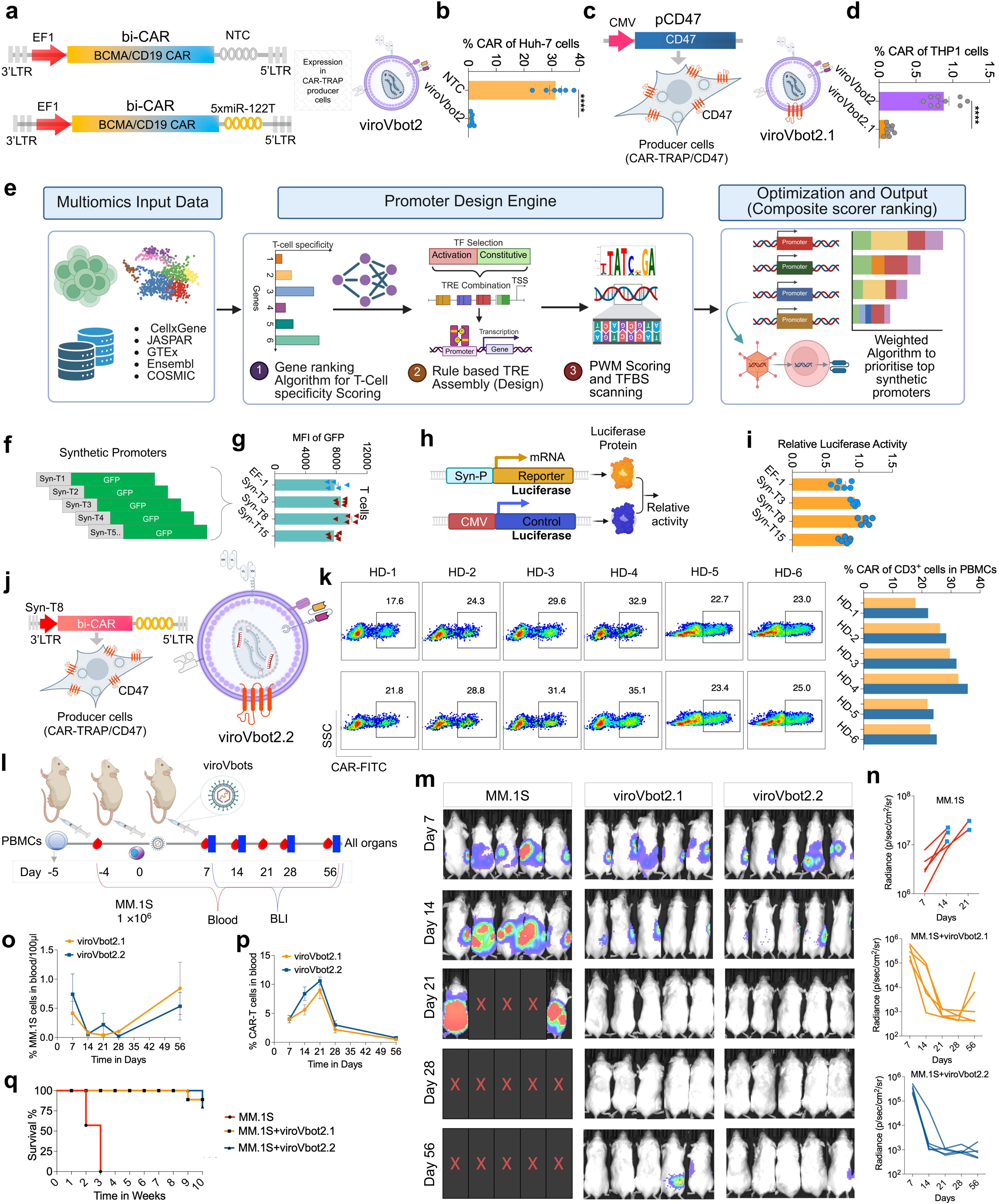
Layered engineering of viroVbots with PromoterForge-derived T cell promoters, liver specific transgene detargeting, and CD47 shielding. **a,** Schematic of miR-122-mediated hepatocyte-specific transgene silencing strategy incorporating five tandem miR-122 target sequences (5×miR-122T) in the 3’ untranslated region (UTR) downstream of CD3ζ costimulatory domain of bi-CAR transgene (viroVbot2). **b,** Flow cytometry analysis of % CAR expression (BCMA CAR-PE) in Huh-7 hepatoma cells transduced with LVV containing miR-122 (viroVbot2) or non-targeting control shRNA (NTC) with bi-CAR as transgene. **c,** Schematics of CD47 overexpression strategy in CAR-TRAP producer cells for macrophage evasion (viroVbot2.1). **d,** Flow cytometry analysis of % CAR expression in THP1 cells transduced with either viroVbot2 or viroVbot2.1. **e,** Schematic flow diagram of PromoterForge pipeline *(details of the pipeline are provided in methods)*. **f,** Candidate synthetic promoters fused upstream with GFP reporter cassette in reporter-plasmid format. Each synthetic promoter represents a unique combination of core promoter elements, TFBS motifs and enhancer arrangements. **g,** Flow cytometric analysis of GFP expression as MFI in primary human T cells transduced with constructs in which GFP is driven by the EF-1α promoter or the indicated synthetic promoters (Syn-T3, Syn-T8, Syn-T15) (n=5 biologically independent samples). **h,** Schematic of the dual-luciferase reporter assay used to validate promoter activity. **i,** Relative luciferase activity in T cells transfected with reporter constructs containing EF-1α or the indicated synthetic promoters (Syn-T3, Syn-T8, Syn-T15), normalized to the CMV control (n=6 biologically independent samples). **j,** Schematic of the viroVbot2.2 construct and LVV design. The transfer vector encodes bi-CAR under the Syn-T8 promoter with mir-122. Producer cells co-express the CAR-TRAP system and CD47, generating viroVbot2.2 particles displaying CD47 on the envelope. **k,** Flow cytometry analysis of CAR expression on CD3⁺ T cells in PBMCs from six healthy donors (HD-1 to HD-6) following ex vivo transduction with viroVbot2.1 (top row) and viroVbot2.2 (bottom row). Numbers in gates indicate the percentage of CAR-FITC⁺ cells. The bar graph shows the %CAR⁺ of CD3⁺ cells per donor. **l,** Scheme of the humanized mouse model. NCG mice were engrafted with human PBMCs (day -5), inoculated with 1×10⁶ MM.1S multiple myeloma cells (day -4), and treated with a single dose of viroVbots (day 0). Peripheral blood was collected and BLI performed on days 7, 14, 21, 28, and 56; all organs were harvested at endpoint (day 56). **m** Representative BLI images of MM.1S tumor burden in mice treated with PBS control (MM.1S), viroVbot2.1, or viroVbot2.2 at the indicated time points. Red “X” denotes deceased animals. **n,** Quantification of whole-body BLI radiance (p/sec/cm^2^/sr) over time for each treatment group. Each line represents an individual mouse (n=5 per group). **o,** Percentage of circulating MM.1S tumor cells per 100 µl of blood over time in mice treated with viroVbot2.1 (orange) or viroVbot2.2 (blue). Data shown as mean ± SEM. **p,** Percentage of CAR⁺ T cells in peripheral blood over time in the same treatment groups, demonstrating in vivo expansion kinetics and contraction of CAR-T cells. **q,** Kaplan-Meier survival curves of mice bearing MM.1S tumors and treated with PBS (red), viroVbot2.1 (orange), or viroVbot2.2 (blue); n = mice per group, log-rank test. Data represents mean ± SEM. ****p < 0.0001. A non-parametric t-test was used for statistical analysis between groups.

These CD47-expressing producer cell lines were used to generate bi-CAR lentiviral vectors, designated viroVbot2.1. Evaluation in monocytic and macrophage cell lines demonstrated robust decreases in lentiviral vector uptake (**Supplementary Fig. 17c, d**). These results were confirmed in PBMCs from multiple myeloma patients, showing significant reductions in viroVbot2.1 uptake by monocytes compared to viroVbot2 (**Supplementary Fig. 17e**). Combination of 5xmiR-122T-mediated hepatocyte silencing, CAR-TRAP producer cells, and CD47 expression effectively minimized off-target lentiviral vector delivery.

### PromoterForge enables the design of T cell-specific promoters

To further enhance safety and minimize off-target expression, we developed an AI-based synthetic promoter pipeline for T cells, termed PromoterForge (**Fig. 5e and** *Methods*). PromoterForge was designed based on three guiding principles: (1) incorporation of transcription factor binding sequences with high T cell expression; (2) inclusion of promoter core regions exhibiting T cell-specific activity; and (3) minimal activity in non-T cell types. To minimize transgene packaging constraints, we limited all synthetic promoters to lengths under 200 base pairs. Using this pipeline, we generated 15 T cell-specific synthetic promoters (**Table 7**). GFP screening identified optimal T cell-specific promoters (**Fig. 5f**). Multiple promoters demonstrated T cell-specific activity with minimal off-target expression (**Supplementary Fig. 18a, b**). The best-performing promoter (Syn-T8) was subsequently evaluated in T cells alongside more than 20 additional cell lines (**Fig. 5g and Supplementary Fig. 18c**). Syn-T8 activity was further confirmed using dual-luciferase system, demonstrating superiority over EF-1α (**Fig. 5h, i**). Importantly, robust promoter activity was also observed across 12 patient-derived T cell samples, demonstrating modestly higher activity than endogenous EF-1α promoter activity (**Supplementary Fig. 18d**). Syn-T8 exhibited superior T cell-specific activity compared to all tested alternatives. Detailed analysis of PromoterForge development and validation is provided in the **Supplementary Information-I**.

We next combined the Syn-T8 promoter with viroVbot2.1 to generate optimized lentiviral vectors, designated viroVbot2.2 (**Fig. 5j**). Transduction efficiency (at MOI 2.5) was >20% across all analyzed samples, including healthy control-derived PBMCs and patient-derived T cells (**Fig. 5k & Supplementary Fig. 19a**). Anti-tumor activity evaluation comparing viroVbot2.1 and viroVbot 2.2 confirmed that tumor control was maintained with the T cell-specific synthetic promoter (**Supplementary Fig. 19b, c**).

These viroVbots were evaluated in in vivo studies using humanized mouse models. Tumor-bearing multiple myeloma humanized mouse models were developed and administered healthy donor-derived PBMCs (**Fig. 5l**). Blood samples were collected at regular intervals for analysis, and all organs were harvested at study termination for CAR analysis. Efficacy analysis evaluated anti-tumor responses through bioluminescence imaging, tumor-related symptoms, and CAR-T cell presence in blood. Both viroVbot2.1 and viroVbot2.2 significantly eliminated tumors over time (**Fig. 5m, n**). Early tumor clearance occurred in both viroVbot2.1 and viroVbot2.2 recipients. Detection of CAR-T cells confirmed robust in vivo CAR-T cell generation by both the viroVbots, which correlated with clearance of MM.1S cells (**Fig. 5o, p**). However, several mice showed tumor relapse over time, leading to reduced survival (**Fig. 5-m-q**). Analysis of blood and organs revealed no organ-related toxicity or CAR-T cell presence in any studied organs or non-T cell types (**Supplementary Fig. 20**). Particularly, viroVbot2.2 was not detected in any other analyzed cell types except T cells. These results demonstrate robust in vivo CAR-T cell generation specifically in T lymphocytes without off-target effects, with CAR-T cells detectable only in blood.

### Minimal IL-7R chimera with sBCMA neutralization induces durable CAR-T cell proliferation and enhanced anti-tumor response

Though highly promising, in vivo CAR-T cell generation faces different challenges compared to ex vivo manufacturing, including lower transduction efficiency that results in reduced initial CAR-T cell numbers, requiring enhanced strategies to promote T cell persistence, proliferation, and anti-tumor activity. Therefore, to enhance in vivo CAR-T cell durability and anti-tumor responses, we developed chimeric signaling receptors designed for dual functions; inducing T cell proliferation through cytokine signaling and neutralizing soluble BCMA as decoy receptors. We generated de novo sBCMA-targeting binders with high binding affinity for soluble BCMA, structural stability, but reduced affinity for membrane-bound forms (**Table 8**). The top three peptides were selected for in vitro validation (**Fig. 6a, Supplementary Fig. S21a & Supplementary Information-II**). We utilized IL-7Rα and synthesized minimal intracellular domains designed to retain STAT5 activation, based on previously validated functional studies.^42^ Three minimal IL-7Rα domains were created and combined with the selected sBCMA binding peptides (designated sBC-Ps) using G4S linkers and CD8α transmembrane domains, resulting in 12 chimeric signaling receptor variants. These receptors were individually evaluated in vitro in primary T cells to assess STAT5 phosphorylation using PathScan Phospho-Stat5 ELISA assays (**Fig. 6a & Supplementary Fig. S22a**).

**Figure 6:**
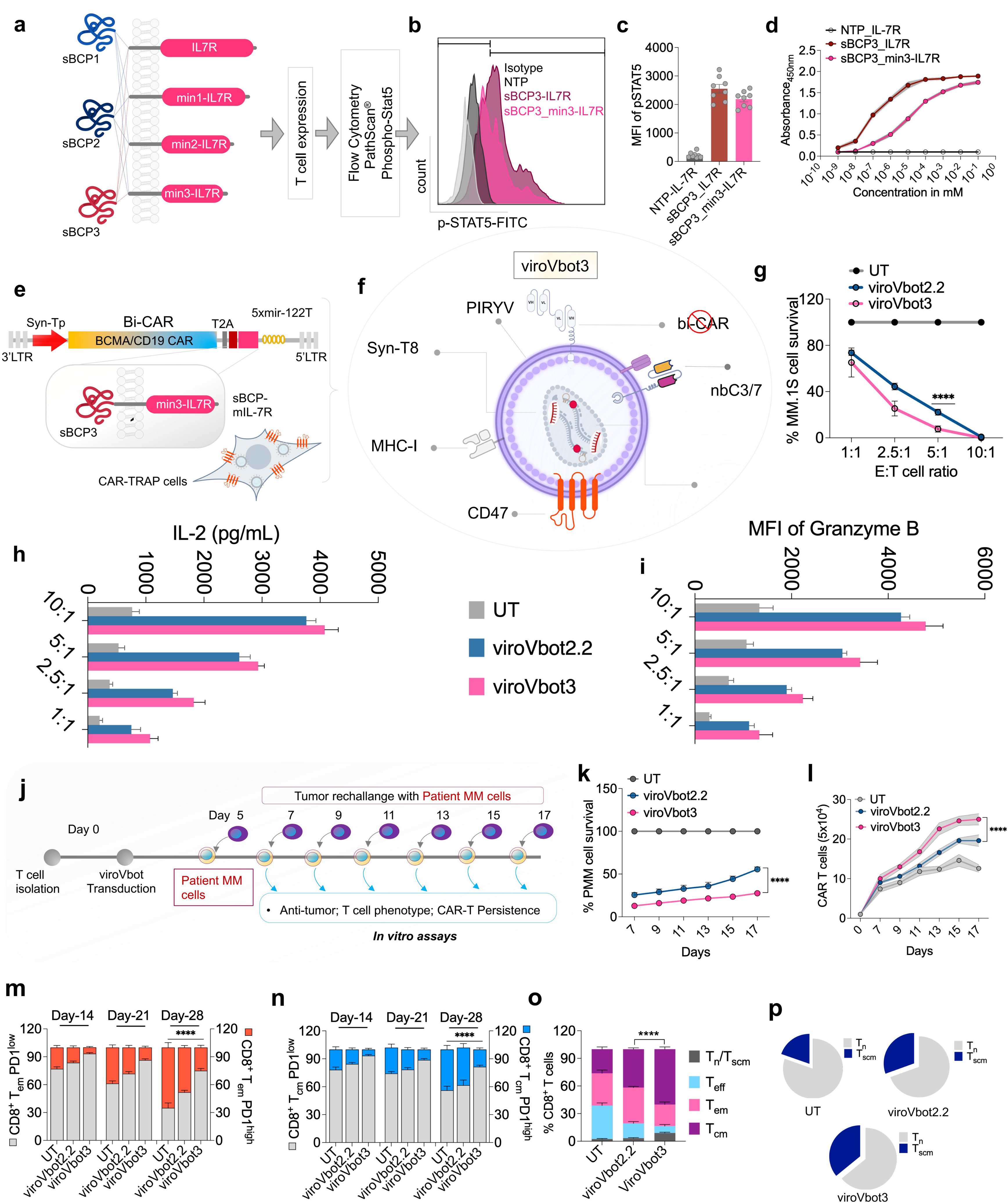
A dual CAR with soluble BCMA-targeting receptor for enhanced T cell persistence and tumor control. **a,** Schematic of the screening strategy for selecting a synthetic IL-7R agonist. Three synthetic binding proteins (sBC-P1, sBC-P2, sBC-P3) were paired with full-length IL-7R or minimized IL-7R variants (min1-, min2-, min3-IL7R), expressed in T cells, and assessed for downstream STAT5 phosphorylation using the PathScan Phospho-STAT5 assay. **b,** Representative flow cytometry histograms of phospho-STAT5 (p-STAT5) in T cells expressing the indicated constructs, compared with isotype and non-transduced (NTP) controls. Anti-p-STAT5 antibody was used followed by Alexafluor 488 and acquired in FITC, channel. **c,** Quantification of pSTAT5 MFI across conditions, showing comparable STAT5 activation by sBCP3-IL7R and the minimized sBCP3_min3-IL7R receptor relative to NTP-IL7R control (n=8 biologically independent samples). **d,** Dose-response curves of ligand-induced receptor activation measured by ELISA across a concentration range (mM) for NTP-IL7R, sBCP3-IL7R, and sBCP3_min3-IL7R (n=5 biologically independent samples). **e, f** Schematic of the viroVbot3 transfer vector and producer cell design. The bi-CAR (BCMA/CD19) cassette is driven by the Syn-Tp promoter, linked via T2A, and detargeted from hepatocytes by 5× miR-122 target sites in the 3′ UTR to co-express the synthetic sBCP3-min3-IL7R (sBCP-mIL7R) cytokine receptor module. **g,** In vitro cytotoxicity assay showing % MM.1S tumor cell survival at increasing effector-to-target (E:T) ratios following co-culture with untransduced T cells (UT), viroVbot2.2-, or viroVbot3-generated CAR-T cells (n=5 biologically independent samples) after 24 h. **h,** IL-2 secretion (pg/mL) by CAR-T cells co-cultured with MM.1S target cells at the indicated E:T ratios (n=5 biologically independent samples). **i,** Similarly, intracellular Granzyme B expression (MFI) in CAR-T cells across the same E:T ratios (n=5). **j,** Schematic of the serial tumor-rechallenge assay. **k,** Percentage of Patient MM (PMM) cell survival over time during serial rechallenge in co-cultures with UT, viroVbot2.2, or viroVbot3 CAR-T cells. **l,** Absolute CAR-T cell counts (5×10^4^) during serial rechallenge, demonstrating superior expansion and persistence of viroVbot3-generated CAR-T cells. **m,** Bar graph of frequency of PD1^low^ (gray) versus PD1^high^ (orange) populations within CD8⁺ effector memory (T_EM_) cells at days 14, 21, and 28 of co-culture for the three groups. **n,** Similarly, CD8⁺ central memory (T_cm_) cells at the same time points (n=5 biologically independent samples). **o,** Memory subset distribution (% of CD8⁺ T cells); naive/stem-cell memory (T_n_/T_scm_), effector (T_eff_), effector memory (T_em_), and central memory (T_cm_), across UT, viroVbot2.2, and viroVbot3 groups. **p,** Pie charts showing the relative proportions of T_n_ (gray) and T_scm_ (blue) compartments within CD8⁺ T cells (n=5 biologically independent samples). Data represents mean ± SEM. ****p < 0.0001. A non-parametric t-test was used for statistical analysis between groups.

sBCP-mIL-7R (designated sBC-P3_min3-IL7R) with optimal pSTAT5 activation, was selected for further development (**Figure 6b, c**). Dose-response study using purified sBCMA confirmed dose-dependent pSTAT5 activation (**Figure 6d**). To validate receptor functionality under physiological conditions where sBCMA is secreted by multiple myeloma cells, we utilized supernatants from MM.1S cells and bone marrow aspirates from patient samples. Under both conditions, sBCMA presence optimally activated pSTAT5 in CAR-T cells, confirming efficient signaling by these chimeric receptors (**Supplementary Fig. 22b-d**). We subsequently created transgenes simultaneously expressing bi-CAR and sBCP-mIL-7R, generating lentiviral vectors in CAR-TRAP cells designated viroVbot3 based on our previously established process.^38,43^ Transduction efficiency remained comparable to viroVbot2.2 (**Fig. 6e, f & Supplementary Fig. 22e**). Patient-derived CD3^+^ T cells were used for functional activity evaluation. Anti-tumor activity demonstrated robust cytotoxicity mediated by viroVbot3 against MM.1S cells and patient-derived multiple myeloma cells (**Fig. 6g & Supplementary Fig. 22f**).

viroVbot3 induced robust granzyme B and IL-2 production when co-cultured with patient-derived myeloma cells, with somewhat higher levels observed with viroVbot3, reaching statistical significance in most samples (**Fig. 6h, i**). To assess long-term persistence of these viroVbot variants, we used a tumor rechallenge (TR) model evaluating patient-derived CD3^+^ T cells upon re-challenge with patient-derived myeloma cells (**Fig. 6j**). viroVbot3 demonstrated superior anti-tumor activity compared to viroVbot2.2, which correlated with significantly higher T cell persistence under TR conditions (**Fig. 6k, l**). This improved persistence was accompanied by reduced exhaustion of memory cell populations in the viroVbot3 group, with corresponding increase in central memory cells and naive and memory stem T cells (**Fig. 6m-p**). These findings demonstrate that IL-7R signaling significantly enhances T cell anti-tumor activity and improves long-term CAR-T persistence.

### viroVbot3 demonstrates safety and efficacy across multiple xenograft models and B cell malignancies

The preclinical efficacy of viroVbot3 was evaluated in humanized mouse models engrafted with patient-derived PBMCs, using the lowest viroVbot3 dose (0.25 x 10^6^ TU/mouse) with 5 mice per group (**Fig. 7a**). Mouse models were initially developed by engrafting MM.1S cells under TR conditions. Secondary tumor challenge was administered on day 40 post-CAR-T cell administration, using MM.1S cells lacking BCMA and CD19 expression (BCMA^-/-^/CD19^-/-^) and expressing only GPRC5D. Instead of PIRYV pseudotyped lentiviral vectors, we used eCOCV-G and eVSV-G-based vectors for re-dosing, which had been optimized for reduced immunogenicity (*Figure 1*). This approach will ensure that if mice (and eventually humans in clinical translation) developed antibodies against PIRYV envelope, alternative COCV-G or VSV-G mutant-based pseudotypes could be used for re-dosing. viroVbot3 achieved sustained tumor clearance (**Fig. 7b, c**). The survival kinetics revelated higher survival rates of the mice treated with viroVbot3 (**Fig. 7d**) and the BCMA-positive CAR-T cells were detected in blood at all studied time points (**Fig. 7e**).

**Figure 7:**
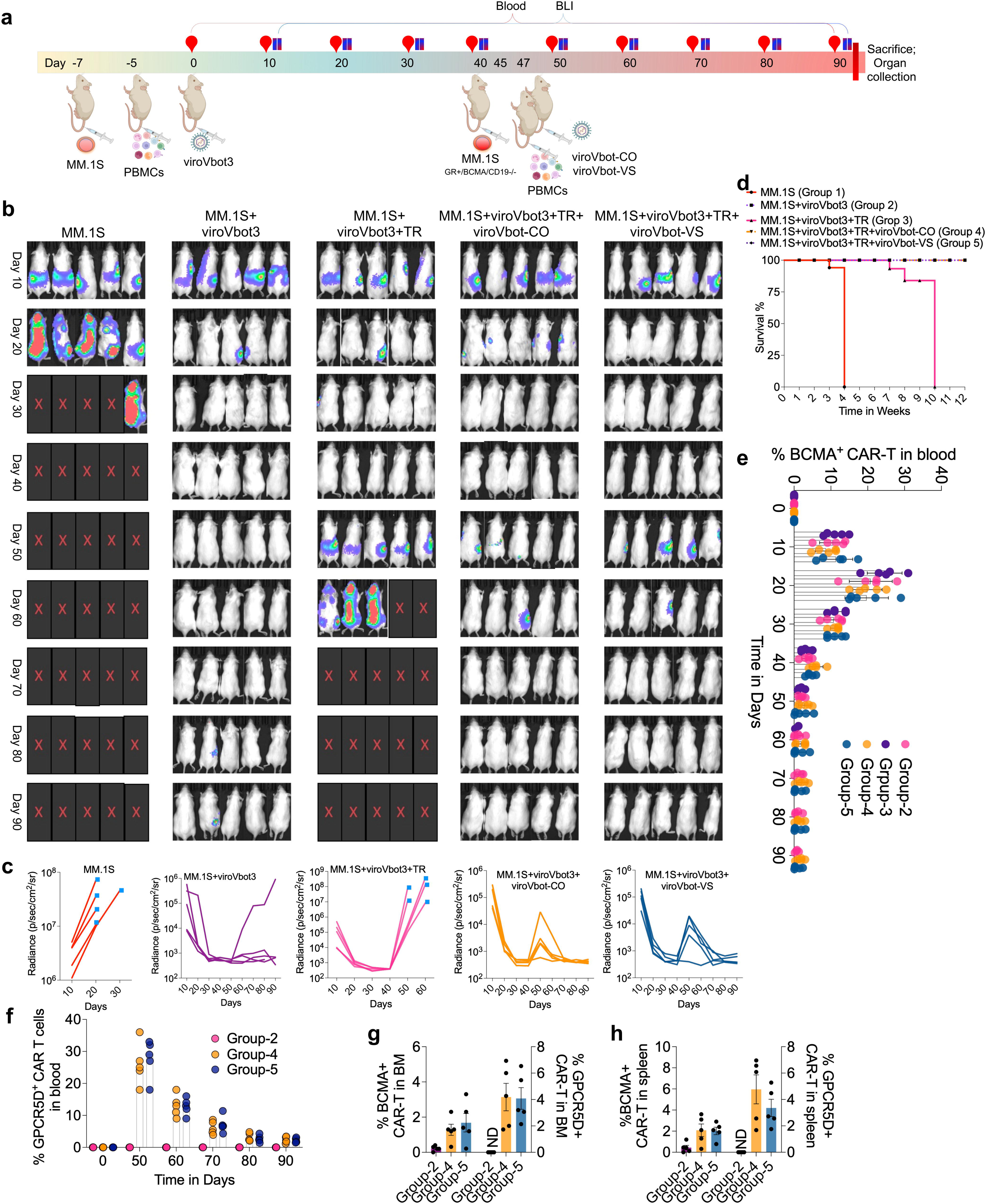
viroVbot3 achieves durable tumor control with redosability via heterologous envelope pseudotypes. **a,** NCG mice received MM.1S cells (day -7), human PBMCs (day -5), and viroVbot3 (day 0). On day 45, mice were rechallenged with BCMA negative but expressing GPRC5D (GR) MM.1S (GR⁺/BCMA⁻/CD19⁻) and given a second PBMC infusion plus viroVbot-CO or viroVbot-VS on day 47. Blood and BLI were collected at indicated time points; organs were harvested at day 90. **b,** Representative BLI images of tumor burden over time (days 10-90) in five treatment groups as indicated (n=5 mice in each group). **c,** Quantification of whole-body BLI radiance (p/sec/cm^2^/sr) over time for each group; each line represents an individual mouse. **d,** Kaplan-Meier survival curves of the five groups (Group-1 to Group-5) across 12 weeks. Log-rank test (n=10 mice in each group). **e,** Longitudinal flow cytometric quantification of BCMA⁺ CAR-T cells (% of T cells) in peripheral blood from days 0-90 across Groups 2-5, showing initial expansion and contraction kinetics of the first-line CAR-T population. **f,** Frequency of GPRC5D⁺ CAR-T cells (% of T cells) in peripheral blood across Groups 2, 4, and 5, demonstrating expansion of the second-line viroVbot-CO/VS-derived CAR-T cells following antigen-loss rechallenge. **g,** Quantification of BCMA⁺ (left axis) and GPRC5D⁺ (right axis) CAR-T cells in bone marrow (BM) at endpoint (day 90) for Groups 2, 4, and 5. ND, not detected. **h,** Similarly in spleen. Data represents mean ± SEM. A non-parametric t-test was used for statistical analysis between groups.

After 40 days, TR was performed using BCMA^-/-^ but GPRC5D expressing MM.1S cells, followed by PBMC administration from the same patients and viroVbot3 treatment with either COCV-G (viroVbot3.1) or VSV-G (viroVbot3.2). All tumor-rechallenged mice treated with viroVbot3.1 or viroVbot3.2 showed positive responses. Notably, mice treated with viroVbot3 without TR also survived beyond 50 days. Evaluation of BCMA CAR-T cells revealed persistent cell populations in surviving mice from the viroVbot3 group at 90 days (**Fig. 7e**). Further, in TR groups treated with viroVbot3.1 or viroVbot3.2, GPRC5D CAR-T cells remained present until the last analysis at 90 days (**Fig. 7f**).

To determine off-target effects, we analyzed all major organs and blood cell populations. No CAR-T cells were detected in any organs except minimal populations in spleen, and bone marrow, with no CAR-T cell or CAR transgene expression detected elsewhere (**Fig. 7g, h & Supplementary Fig. 23a-c**). Similarly, no CAR expression was observed in any analyzed blood cell populations other than T and NK cells (**Supplementary Fig. 23d**). Further serum biochemistry analysis revealed no evidence of systemic toxicity, with all measured parameters remaining within normal physiological ranges (**Supplementary Fig. 24**).

To expand viroVbot technology applications to solid tumors, we developed a gastric cancer model based on our previous study.^38^ Mice were initially treated with viroVbot3 encoding humanized CAR targeting CLDN18.2 (designated viroVbot3S). We similarly generated viroVbots with COCV-G (viroVbot3.1S) or VSV-G (viroVbot3.2S) envelope proteins. The ASG gastric engraftment animal model was engrafted with PBMCs followed by designated viroVbot3S administration. Subsequently, mice received viroVbot3.1S and viroVbot3.2S at one-month intervals (**Fig. 8a**). Results indicate that viroVbots efficiently eliminated tumors and increased mouse survival; however, effects were not sustained as significant decline in circulating CAR-T cells occurred over time, resulting in decreased survival (**Fig. 8b-d**). To overcome this reduced efficacy, we re-administered mice with viroVbot3.1S, achieving higher sustained efficacy with increased survival and decreased tumor volumes over time (**Fig. 8b-d**). CAR-T cell analysis revealed improved persistence of BCMA CAR-T cells with re-dosing (**Fig. 8e**). Further, re-dosing increased circulating CAR-T cell numbers in blood.

**Figure 8:**
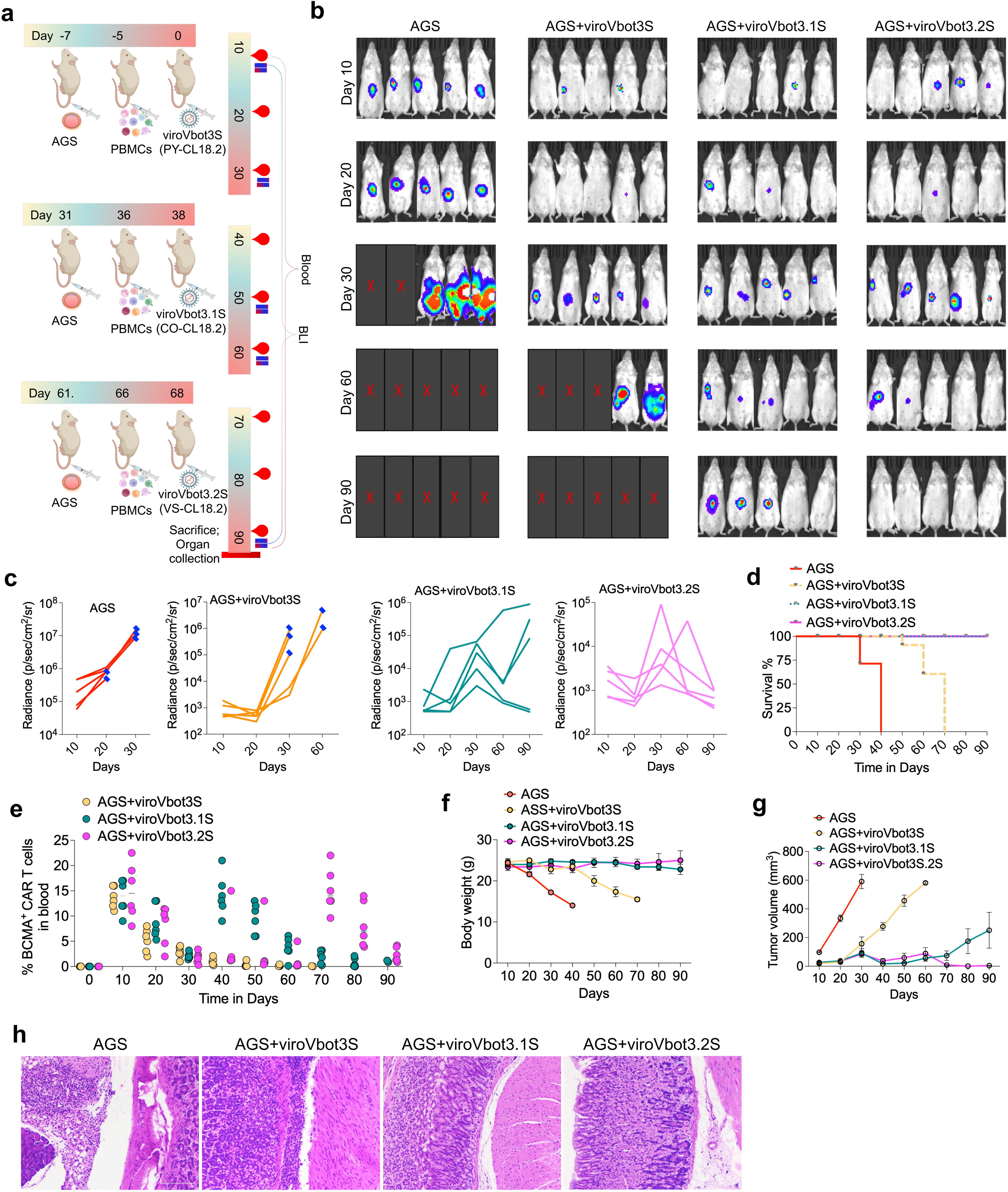
viroVbot3 extends efficacy to solid tumors via Claudin18.2-targeting CAR. **a,** Schematic of the serial antigen-escape rechallenge model in AGS gastric cancer. NCG mice were injected with AGS tumor cells (day -7), engrafted with human PBMCs (day -5), and treated with viroVbot3S (PY-CL18.2) on day 0. Two subsequent rounds of tumor rechallenge with AGS cells (days 31 and 61) were followed by additional PBMC infusions (days 36 and 66) and sequential dosing of viroVbot3.1S containing eCOCV-G (CO-CL18.2, day 38) and viroVbot3.2S containing eVSV-G (VS-CL18.2, day 68), with CLDN18.2-targeted CAR. Peripheral blood and BLI were collected at the indicated time points. **b,** Representative BLI images of AGS tumor burden over indicated time points across four groups. Red “X” denotes deceased animals. **c,** Quantification of whole-body BLI radiance (p/sec/cm^2^/sr) over time for each group; each line represents an individual mouse. **d,** Kaplan-Meier survival curves of mice. **e,** Longitudinal flow cytometric quantification of CLDN18.2-CAR⁺ T cells (% of circulating T cells) in peripheral blood from days 0-90 across the three viroVbot-treated groups. **f,** Body weight (g) of mice over the course of the experiment as a measure of overall tolerability. **g,** Tumor volume (mm³) measured by caliper over time across the four groups. **h,** Representative haematoxylin sections of tumor/gastric tissue from each group at endpoint, showing dense tumor infiltrate in untreated AGS controls and progressive tumor clearance with restored tissue architecture in viroVbot3.1S- and viroVbot3.2S-treated mice. n=5 mice per group. Data represents mean ± SEM. A non-parametric t-test was used for statistical analysis between groups. Scale bar; d: 200 μm.

To find whether a third viroVbot dose would further enhance in vivo CAR-T cell efficacy, viroVbot3.2S was administered in the mice. The third viroVbot injection completely eradicated tumors, with all mice maintaining good health at the final data collection point (**Fig. 8f, g**). Tumor assessment of recovered tissues showed significant reductions in gastric tumor mass and no toxicity (**Fig. 8h**). The viroVbot platform achieves durable, redosable in vivo CAR-T cell generation across both hematologic and solid tumor models, with envelope diversification (PIRYV, COCV-G, and VSV-G mutants) allowing sequential administration to sustain antitumor activity while maintaining a favorable off-target profile.

## Discussion

In vivo CAR-T cell generation is a revolutionary paradigm shift toward scalable, affordable, and accessible cellular immunotherapy, but significant immunological and safety barriers persist.^44^ Early lentiviral and lipid nanoparticle (LNP) platforms have demonstrated functional in-body CAR-T generation and tumor control in preclinical studies, and the ESO-T01 and HN2301 Phase I trial confirms clinical feasibility.^12,45,46^ However, ESO-T01 also revealed substantial toxicity challenges, with Grade 3 cytokine release syndrome occurring in 80% of patients, pointing up that immunogenicity remains a critical bottleneck in in vivo CAR-T therapy. A fundamental limitation stems from wild type VSV-G-pseudotyped vectors, despite efficiency in ex vivo applications, VSV-G undergoes complement-mediated neutralization systemically, induces robust anti-envelope antibody responses that prevent re-dosing, and triggers substantial adaptive immune responses that constrain initial transduction and durability.^19,36,44,47^ To address these critical limitations, we developed viroVbot, a next-generation lentiviral platform.

To overcome VSV-G-mediated immunogenicity, we developed CIMMEXA, an AI-driven platform to identify naturally occurring viral envelopes with reduced MHC-epitope density. Computational analysis of over 1,500 pH-dependent viral envelopes identified Piry virus glycoprotein (PIRYV) as substantially lower in immunogenic burden than VSV-G, COCV-G, and VSAV-G. Unlike previous approaches using pH-independent envelopes,^25,27^ we selected pH-dependent envelopes to ensure fusion requires endocytosis and endosomal acidification, providing a critical safety layer that prevents transduction of non-target cells through transient surface binding. Assessment of pre-existing immune responses in 65 healthy volunteers revealed that PIRYV induced the lowest CD4+ and CD8+ T cell responses, while neutralizing antibody kinetics in immunized mice confirmed minimal anti-PIRYV activity compared to robust VSV-G responses. Critically, PIRYV demonstrated resistance to serum neutralization, consistent with published preclinical data.^19,47^ We further applied CIMMEXA to rationally optimize even low-immunogenic envelopes by designing targeted amino acid substitutions. Computationally optimized VSV-G, COCV-G, and VSAV-G variants showed substantially attenuated immunogenicity while preserving transduction efficiency, establishing a generalizable strategy for envelope engineering beyond conventional approaches.

Previous in vivo CAR-T platforms using retargeted viral envelopes have been limited by incomplete elimination of native viral tropism, reliance on single-receptor targeting, and lower CAR-T transduction, resulting in suboptimal targeting specificity and off-target transduction.^28,48,49^ To overcome these limitations, our dual-targeting strategy combined with an immunogenicity-optimized, receptor-binding-deficient PIRYV envelope (ePIRYV^RBD^) with humanized CD3- and CD7-specific nanobodies (nbCD3/7) showed specific and higher in vivo transduction efficiency. This is the first study to identify critical residues within the low-immunogenic envelope PIRYV receptor-binding domain, which are responsible for LDL receptor family binding, and their mutation completely abolished native receptor binding. Comparative structural assessment with VSV-G and COCV-G further confirmed complete detargeting, including identification of an additional residue required for full receptor ablation in COCV-G. Although previous studies have used wild-type COCV-G, which exhibits off-target transduction,^22^ our approach fully uncouples viral entry from endogenous receptor usage and redirects transduction through two independent T-cell antigens, thereby enhancing targeting precision and cellular coverage while minimizing off-target uptake. Furthermore, the use of compact humanized nanobodies offers superior stability, reduced aggregation propensity, lower immunogenicity, and improved manufacturability compared with conventional scFv-based retargeting strategies.

Dose-dependent transduction studies demonstrated robust and efficient T and NK cell targeting across multiple in vitro and in vivo models. However, a critical clinical safety concern, is inadvertent transduction of malignant B cells through CAR-mediated bridge-binding on viral particles as highlighted by previous studies. ^22,28,29^ This may let leukemic clones to acquire CAR expression and promote tumor escape. To prevent it, we developed CAR-TRAP producer cells that sequester CAR transgenes in the endoplasmic reticulum via KDEL-retention signals, blocking surface display.^50^ This mechanism is simple compared to other previous approaches such as TRiP system,^25^ and completely eliminated transduction of patient-derived multiple myeloma cells and primary B cell populations, providing evidence for the additional layer of safety and addressing a major clinical risk of in vivo CAR-T cell therapy.

Off-target liver and macrophage accumulation due to LDL-R-mediated hepatocyte uptake and Kupffer cell sequestration is also a critical factor that limits therapeutic efficacy and also raising safety concerns.^22,25,51^ We addressed this through a dual-pronged approach by combining liver-specific microRNA silencing with macrophage-inhibitory surface engineering. Integration of previously established miR-122 targeting sequences in our transgene design enables hepatocyte-specific silencing while preserving robust T cell expression, which is a particularly elegant solution that leverages natural tissue-specific microRNA expression patterns without compromising promoter strength.^52–55^ Simultaneously, we incorporated CD47 expression on producer cell surfaces to inhibit macrophage uptake and reduce innate immune activation, which has been validated in previous studies^56^ Our extensive biodistribution analysis reveals negligible CAR expression in liver, lungs, and kidneys compared to control vectors, demonstrating the remarkable efficacy of this safety architecture.

Another, major obstacle of the in vivo CAR-T cell approach is that the promoters such as EF1α drive CAR expression in any transduced cell type, creating toxicity risks from ectopic expression in non-T cell populations.^57^ To address this, we developed PromoterForge, which is an AI-based synthetic promoter discovery pipeline. Through PromoterForge, we generated T cell specific synthetic promoters, which showed T cell-specific activity and were significantly better compared to conventional EF1α across patient-derived T cell samples. This transcriptional innovation transforms CAR-T safety by providing cell-type-specific expression without compromising transduction efficiency. Recent studies have similarly pursued T cell-specific transgene restriction through synthetic promoter design.^58^ However, PromoterForge uniquely combines machine learning-driven promoter discovery with T cell-specific transcription factor profiling to achieve superior specificity and expression levels compared to conventional approaches, while maintaining the transduction efficiency required for clinical efficacy.

CAR-T cell durability remains a fundamental limitation affecting long-term clinical efficacy in a subset of patients.^59^ While in vivo CAR-T approaches may mitigate ex vivo culture-induced exhaustion, recent studies indicate that some patients may still relapse, potentially due to lower persistence.^45,60^ To enhance durability, we engineered chimeric signaling receptors combining IL-7Rα intracellular domains with de novo designed sBCMA-targeting peptides. As IL-7Rα is known to drive memory T cell formation.^61^ This dual-function design provides simultaneous proliferation signals through STAT5 activation while neutralizing soluble BCMA, a known T cell suppressive factor in multiple myeloma microenvironments.^37^ The viroVbot3-transduced T cells exhibited markedly enhanced STAT5 phosphorylation, significantly reduced exhaustion markers, and superior persistence upon tumor rechallenge. Notably, viroVbot3-transduced T cells demonstrated central memory and naïve stem cell phenotypes characteristic of durable, long-lived anti-tumor responses in xenograft models, suggesting enhanced potential for sustained clinical efficacy.

A critical limitation expected to emerge in previous in vivo CAR-T platforms is the inherent biological challenge of redosing patients, as anti-envelope antibodies mediate viral neutralization and prevent therapeutic responses in tumor-rechallenged conditions.^44^ We overcame this through envelope diversification. We sequentially administered viroVbot3 with alternative envelopes (COCV-G-pseudotyped viroVbot3.1 and VSV-G-pseudotyped viroVbot3.2) that maintained robust anti-tumor efficacy upon tumor rechallenge. This approach allowed repeated dosing without neutralization-mediated loss of transduction. Expanding beyond hematologic malignancies, we also demonstrated the application of our platform to solid tumors by targeting CLDN18.2 in gastric cancer models based on our previous ex-vivo CAR molecules.^38^ Sequential triple dosing with envelope diversification achieved complete tumor eradication with sustained survival, demonstrating that redosing progressively enhanced CAR-T persistence and anti-tumor activity. These results establish a generalizable strategy to overcome the inability to sustain therapeutic CAR-T cell numbers through repeated dosing while maintaining anti-tumor efficacy, a longstanding limitation in solid tumor CAR-T responses.^27,62^

Off-target CAR uptake in organs can lead to long-term consequences. However, our extensive preclinical safety analysis shows minimal CAR-T cell organ infiltration and negligible off-target cell transduction, demonstrating that our multi-layered safety approach provides durable protection. CAR-T cells remain restricted to blood and lymphoid tissues, with negligible presence in liver, lungs, kidneys, and other critical organs. CAR expression is confined to T and NK cells, with no detectable transduction of B cells, monocytes, macrophages, or other off-target populations at any time point. Together, these independent safety mechanisms provide reliable redundancy, ensuring a robust safety profile for our platform.

In conclusion, the viroVbot platform reported in this study redefines in vivo CAR-T therapy by eliminating the manufacturing complexities of autologous approaches while delivering the safety and efficacy required for clinical translation. By generating CAR-T cells directly within the patient, this technology offers a scalable, accessible alternative poised to broaden the reach of cell-based immunotherapy. Future work should prioritize early-phase clinical trials with rigorous immunogenicity monitoring to define its therapeutic potential across diverse malignancies and patient populations.

## Methods

### CIMMEXA^TM^ Based Immunogenicity Prediction of Viral Envelope Proteins

All available pH-dependent glycoprotein sequences belonging to the family Rhabdoviridae were retrieved from the NCBI database, yielding a total of 22,562 sequences. The vesicular stomatitis virus glycoprotein (VSV-G) was selected as the reference sequence for homology-based identification of related envelope proteins. Homologous sequences were identified using three complementary sequence similarity search algorithms; BLASTP, PSI-BLAST, and DELTA-BLAST, with a minimum sequence identity threshold of 30%. Results from all three searches were merged, and redundant entries were removed to generate a final non-redundant dataset comprising 641 unique VSV-G homologs for downstream analyses. Functionally conserved homologs were subsequently aligned using ClustalW to assess sequence conservation and identify structurally relevant regions. Full-length envelope protein sequences were then processed using a sliding-window approach to generate overlapping peptide libraries ranging from 9 to 13 amino acids in length, ensuring comprehensive coverage of the MHC class I epitope landscape. Each peptide was evaluated against a comprehensive panel of more than 11,000 HLA alleles using three independent MHC class I binding prediction methods: (i) a pan-allelic artificial neural network (ANN) model trained on experimentally validated ligand elution and binding affinity datasets; (ii) a position-specific scoring matrix (PSSM) approach derived from mass spectrometry-based HLA ligandome data; and (iii) a transformer-based deep learning model trained on a large consolidated peptide-MHC interaction dataset. The use of multiple prediction algorithms minimized method-specific biases and enhanced the robustness of immunogenicity assessment.

Raw predictions from individual algorithms were integrated using the Logistic Probabilistic Ensemble (LPE), generating standardized binding probability matrices for each peptide. LPE parameters were optimized using 59,998 experimentally validated peptide-HLA pairs from the Immune Epitope Database (IEDB) spanning >150 HLA alleles (HLA-A, B, C families). Five-fold cross-validation yielded AUC ROC = 0.9348 ± 0.0015, with 89.2% sensitivity and 91.1% specificity. Composite immunogenicity scores were calculated by aggregating LPE consensus probabilities across all peptide-HLA pairs with weighted contributions. Peptides were classified as strong binders, weak binders, or non-binders. Systematic immunogenicity risk profiles were generated for each viral envelope protein and ranked to identify candidates with lowest predicted immunogenic burden.

Strong binding epitopes were mapped and cross-referenced against different virus glycoproteins. Amino acid positions with high-affinity epitopes were targeted for rational substitutions. Three-dimensional models of wild-type and optimized envelope variants were generated using homology modeling. Molecular dynamics simulations (500 ns) were performed with snapshots collected every 10 ps. Protein stability was assessed via root-mean-square deviation (RMSD; acceptance threshold <2.0 Å). Mutations were retained only if secondary structure elements, particularly in the receptor-binding domain, remained intact.

### Computational Design of Humanized Nanobodies

De novo humanized nanobodies targeting the antigen was generated using CelNFo, an in-house computational pipeline developed at Cellogen Therapeutics that combines framework humanization with structure-guided CDR design to generate antigen-specific VHH candidates. Humanized nanobody frameworks were first constructed by substituting camelid framework residues with their human germline-derived equivalents at non-hallmark positions. Conserved VHH hallmark residues governing domain stability and solubility were retained without modification. The resulting humanized frameworks were used as fixed structural scaffolds in all subsequent design steps.

De novo CDR generation was performed using Rfantibody (Bennett et al., Nature, 2025), a computational antibody design framework built on antibody-specialized RFdiffusion models. The experimentally determined or predicted three-dimensional structure of the target antigen was provided as structural input. RFdiffusion was then used to generate nanobody backbone conformations docked against the antigen surface, with humanized framework residues constrained throughout the sampling process. CDR backbone conformations were iteratively sampled and refined to identify geometries exhibiting high shape and physicochemical complementarity toward the target epitope. Amino acid sequences for each generated backbone were designed using ProteinMPNN, which optimizes residue identities conditioned on the backbone geometry. Multiple sequence variants were produced per backbone to increase sequence diversity across the candidate pool. Structural validation of the designed sequences was carried out using an antibody-specialized RoseTTAFold2 model integrated within the RFantibody workflow. Designed sequences were further evaluated for structural stability, folding, and antigen-binding compatibility using Molecular Dynamic Simulation.

### Receptor Detargeting & Molecular Dynamics

In silico analysis was performed on the crystal structure of wild-type (wt) PIRYV bound to LDL-R to identify residues in proximity to charged residues on the receptor. Contact residues between PIRYV and LDL-R were identified through distance-based contact analysis (<4.0 Å), and opposite-charge substitutions at these residues were introduced to destabilize the binding between PIRYV and the receptor and visualized in PyMOL. Binding free energy changes were quantitatively estimated using MM-GBSA (Molecular Mechanics Generalized Born Surface Area) calculations integrated with MD trajectory analysis. RMSD and per-residue RMSF measurements from three independent MD simulation replicates (500 ns each, 310 K, TIP3P water, 0.15 M NaCl) were calculated to assess trajectory convergence, ensemble stability, and local conformational flexibility of mutated residues and surrounding interface regions.

### T Cell-Specific Synthetic Promoter Design (PromoterForge Platform)

PromoterForge was developed for designing T cell-specific synthetic promoters by integrating data from five public databases. 33.2 million human cells from the CellxGene Census were analysed, including 14.3 million T cells across 12 subtypes (e.g., CD8^+^ cytotoxic, CD4^+^ helper, Treg, naïve, memory, γδ, NKT, and MAIT cells), to identify genes with enriched T cell specific expression. GTEx v8 data from 17,382 samples across 54 tissues were used to evaluate off-target expression in vital organs. Validated transcription factor (TF) binding motifs were obtained from JASPAR 2024, native promoter sequences (500 bp upstream of the transcription start site) from Ensembl GRCh38, and oncogene annotations from COSMIC v101 for safety screening.

Candidate genes were ranked using a composite score combining specificity (0.40), expression (0.25), log2 fold-change (0.20), and detection rate (0.15), where specificity was defined as T_cell_mean / (T_cell_mean + Non_T_cell_mean). The top 100 genes were classified as Novel, Pan T Cell, T Cell Specific, or Common. Twenty-five curated T cell TFs (specificity 0.78-0.99) were grouped into an activation-responsive pool (NFATC1, NFATC2, NFKB1, RELA, JUN, FOS, IRF4) for inducible expression and a constitutive pool (FOXP3, TBX21, GATA3, TCF7, LEF1, BCL11B, RUNX1) for stable baseline expression. Synthetic promoters were assembled in a modular architecture [Distal TREs]-[Spacer]-[Proximal TREs]-[Spacer]-[Core Promoter], using 2-4 copies of each JASPAR-validated motif separated by 6-8 bp spacers (AGTCAGCT) and a TATA box (TATAAA) as the core element. Final constructs ranged from 100 to 300 bp to remain compatible with viral vector cargo limits.

Each promoter was evaluated using five metrics (0-10 scale): Expression = min (10, Total_TREs × 1.2 + Activation_TREs × 0.5); Specificity = (Σ TF_Specificity × Copies / Total_TREs) × 10; Inducibility = (Activation_TREs / Total_TREs) × 10; Safety = max(5, 10 - Constitutive_TREs × 0.8); and Quality = 10 × (1 - |GC_content - 0.5| - Length_penalty). These were combined into a composite score weighted as Specificity (35%), Expression (25%), Inducibility (15%), Safety (15%), and Quality (10%), prioritizing T cell specificity as the principal design objective. The platform was implemented as an interactive web application comprising five modules: a Gene Atlas of 100 ranked T cell-specific genes, a Synthetic Promoter Library of 20 optimized designs, a Custom Promoter Designer with real-time scoring, a Promoter Analyzer for scanning user-supplied sequences, and a TRE Library of the 25 curated TFs. All source data were derived from public repositories, and complete tables are provided in **Supplementary Information-I**.

### BCMA-Targeting Binder Design

The extracellular domain, stalk region, and half transmembrane segment of BCMA were modeled according to sequence motifs established in prior literature.^63^ Computational binder design was performed using RfDiffusion (Baker laboratory) targeting hotspot residues at the interface between extracellular domain and transmembrane region. This positioning favored specificity for soluble BCMA while minimizing cross-reactivity with membrane-anchored forms. Diffusion-based scaffold generation produced diverse protein architectures, followed by sequence optimization and co-folding predictions with AlphaFold2. Designs were filtered stringently based on predicted aligned error values (PAE_interaction < 15) and root-mean-square deviation (< 2.0 angstroms) to the target binding geometry. Five high-confidence designs were subjected to affinity maturation using FoldX-based binding free energy calculations. Computational mutagenesis was performed systematically, generating and evaluating combinations of point mutations at the binding interface. Top-ranking candidates were selected for atomistic molecular dynamics simulations to assess binding stability and conformational stability.

Binding affinities of designed complexes were evaluated through umbrella sampling and free-energy perturbation methods on co-folded models. For final recombinant constructs, membrane-embedded systems were prepared using CHARMM-GUI with appropriate lipid parameterization. All-atom molecular dynamics simulations using AMBER22 assessed long-term conformational stability, transmembrane helix insertion, and structural integrity of chimeric receptors in lipid bilayer environment.

Selected sBCMA binders were fused via flexible G4S linker to the CD8 transmembrane domain and three different minimal intracellular domains of IL-7 receptor alpha-chain, generating 12 chimeric receptor combinations designed for ligand-induced STAT5 phosphorylation upon target engagement.

### Plasmid Design and Synthesis

VSV-G was obtained from Addgene (#12259), while related vesiculovirus glycoproteins (VSNJ-G; CPV-G; ISV-G; CARV-G; COCV-G; MRV-G; VSM-G; VSJ-G; VSP-G; PIRYV; and VSAV-G, as shown in ***Figure 1b***) were obtained from NCBI (**Table 9**). All the wild type and computationally optimized envelope variants with key mutations were synthesized by GenScript. All envelope expression constructs were assembled into the pHCMV backbone between EcoRI and KpnI restriction sites under CMV promoter control. The bispecific BCMA/CD19 CAR construct (bi-CAR) incorporating dual 4-1BB/ICOS costimulatory domains was assembled into self-inactivating lentiviral transfer plasmids (based on our previously generated vector backbone;^38^ driven by EF1α or the T cell-specific synthetic promoters generated by PromoterForge. Five tandem miR(microRNA)-122 target sequences were inserted into the 3’ UTR downstream of CD3ζ by synthesizing along with the bi-CAR plasmid. A non-targeting miR sequence was used based on our previous study.^64^ GPRC5D-targeting CAR (GC5B390) and Claudin 18.2-targeting CAR with Whitlow-engineered scFv linkers were developed as described previously.^38^ All constructs were verified by Sanger sequencing across coding sequences and cloning junctions.

Producer cell engineering plasmids included CAR-TRAP (CD19-ECD-KDEL with ER retention signal) and CD47 (NCBI NM_001777) expression constructs were cloned into pEGF-puro using Nhe1 and BsrG1 under CMV promoter. HEK293T cells were transfected with expression plasmids and at 48 h post transfection, cells were subjected to puromycin/blasticidin (0.8-2 ug/mL) selection until non-transfected control cells were eliminated. An empty-vector control producer cell line (CAR-TRAP plus empty vector) was generated in parallel under identical selection conditions. CAR constructs combining full and minimal IL-7Rα intracellular domains with de novo soluble BCMA-targeting peptides, G4S linkers, and CD8α transmembrane domains were synthesized commercially and cloned as previously mentioned.^38^ In-house lentiviral packaging system was generated using transgene from the plasmids originally developed Addgene vectors: pMD2.G/VSVG (envelope), pRSV-Rev (Rev protein), and pRRE (gag/pol), with codon optimization and kanamycin selection markers. Transfer vectors encoding GFP/luciferase reporters or CAR transgenes under CMV/EF1α or T cell-specific promoters were designed with self-inactivating LTRs and puromycin/blasticidin resistance markers as previously described.^38^ T cell receptor-specific humanized nanobodies (anti-CD3, -CD4, -CD5, -CD7, -CD8, -TCRα) and T cell ligands (CD80, CD252; NCBI IDs 941, 7292) were synthesized with N-terminal myc tags and cloned into pHCMV backbone (EcoRI/KpnI) under CMV promoter control.

### Lentiviral Vector Production and Concentration

Self-inactivating (SIN) replication incompetent lentiviral vectors pseudotyped with wild-type and engineered vesiculovirus envelope glycoproteins were produced by transient transfection of HEK293T cells as previously described.^38,43^ Briefly, 5 × 10^6^ HEK293T cells were seeded in 10 cm tissue culture plates. After 24 h, cells were transfected using Lipofectamine 2000 or PEI with the following plasmids: (1) lentiviral genome plasmid encoding a transgene (luciferase; CAR; GFP) as mentioned in the results; (2) packaging plasmid (in-house generated); (3) envelope glycoproteins (in-house synthesized wild type and mutated and VSV-G with empty vector as control; Addgene #12259). Vector-containing supernatant was harvested 48 and 72 h post-transfection, clarified by centrifugation, and concentrated 100-fold by ultracentrifugation (Beckman coulter) at 22,000 rpm for 2 h at 4°C using a sucrose cushion (20% w/v) or also the LVV concentration as performed using an in-house developed LVV concentrator (LV^DOTM^). Further, a GMP grade vector batch for viroVbot3 was generated using the chromatography systems (ACHROM BASE; Genetix) and Tangential Flow Filtration system (Tanfil 100; Rocker Scientific Co., Ltd; Genetix). Vectors were resuspended in either in the cell culture media or TSSM buffer (100 mM sodium chloride, 20 mM Tris pH 7.3, 10 mg/mL sucrose, 10 mg/mL mannitol), aliquoted, and stored at -80°C until use. Vector particle concentration was quantified by HIV-1 p24 capsid antigen ELISA (Takara or ABL) according to the manufacturer’s instructions and transducing titer (TU/mL) was determined by serial dilution on target cell lines by flow cytometry, as described previously.^38,43^

### Cell line culture and Screening Assay

All cell lines (**Table 9**) were cultured in appropriate media according to manufacturer’s guidelines and maintained at 37°C in 5% CO_2_ and 95% humidity. Cell lines were routinely tested for mycoplasma contamination and used within 15 passages after thawing.

viroVbot1 vectors encoding bispecific BCMA/CD19 CAR transgene were evaluated against a panel of 65 organ and tissue-specific cell lines encompassing hematopoietic, lymphoid, myeloid, hepatic, endothelial, epithelial, and specialized cell types to establish T cell-selective targeting specificity and identify off-target transduction risks. Cell lines were cultured in optimal media as indicated in **Table 9**, harvested to 2-5 × 10^5^ cells/mL, and transduced with viroVbot vectors at normalized particle inputs (MOI 2.5) for 7 days at 37°C in 5% CO^2^. CAR transgene expression was assessed by flow cytometry, with isotype controls and/or wild type vector transduced cell lines analyzed in parallel.

### CRISPR-Cas9-Mediated Knockout Cell lines

Raji B lymphoma cells and multiple myeloma-derived cell lines with targeted knockouts of BCMA and CD19 were generated using CRISPR-Cas9 genome editing following our previously established protocol.^41^ Guide RNA sequences for all knockout targets are provided in **Table 9**. MHC-I knockout HEK293T cells were similarly generated using the same CRISPR-Cas9 methodology, with gRNA design targeting B2M gene (Beta-2-Microglobulin) to eliminate surface MHC-I expression. All knockout clones were validated by flow cytometry and confirmed to be MHC-I negative.

### Confocal Microscopy and Colocalization Analysis

Nanobodies and surface-displayed ligands were immunostained and imaged as previously described.^65,66^ Briefly, fixed and permeabilized cells were stained with fluorophore-conjugated antibodies against myc-tag (nanobodies), then visualized on a Zeiss LSM 900 confocal microscope using sequential scanning at 405 nm (DAPI) and 488 nm (Alexa Fluor 488). Z-stacks were acquired at 0.5 µm intervals with 1 Airy unit pinhole diameter. Colocalization between CAR and nanobody signals was quantified using Pearson’s correlation coefficient (minimum n = 20 cells per condition) in ImageJ/Fiji.

CAR-TRAP localization (intracellular ER-retained versus surface-absent) was confirmed by confocal microscopy using anti-G4S linker antibody and rabbit anti-calnexin (1:500, Cell Signaling; #2679) as ER marker, or by flow cytometry comparing permeabilized versus non-permeabilized staining. Surface CD47 expression on CD47-overexpressing producer lines was similarly validated by confocal immunofluorescence (rat anti-CD47, 1:200) and flow cytometry.

### Primary T Cell Isolation and Enrichment

Peripheral blood mononuclear cells (PBMCs) were isolated from whole blood by density gradient centrifugation using Ficoll-Paque as per manufacturer’s protocol. Primary CD3^+^ T cells were enriched from PBMCs using negative immunomagnetic selection (Miltenyi pan T cell isolation kit or Stemcell Technologies), following the procedure as previously described.^38,43^ Enriched CD3^+^ T cells (purity >95%) were resuspended in TexMACS medium supplemented with 2.5% human serum, GlutaMAX, and cytokines (10 ng/mL IL-7 and IL-15) at 1 × 10⁶ cells/mL. For subset-specific studies, CD4^+^ and CD8^+^ T cells were further isolated from CD3^+^ populations via positive selection using EasySep kits (Stemcell Technologies), with post-enrichment purity >95% confirmed by flow cytometry.

### Patient-Derived Multiple Myeloma Cells

Fresh bone marrow aspirates were enriched for plasma cells via negative-selection immunomagnetic enrichment (RosetteSep Human Multiple Myeloma Cell Enrichment Cocktail) as previously described.^37^

### T Cell Activation and Transduction

Primary CD3^+^ T cells were activated with anti-CD3/CD28 Dynabeads (1:1 ratio, 24 h) as a positive control, then transduced with lentiviral vectors and analyzed for activation markers (CD69) by flow cytometry. Patient-derived PBMCs were similarly assessed for CAR expression by flow cytometry at day 7 post-transduction. Transduction efficiency with ePIRYV^RBD+nbCD3/7^ vectors was directly compared to standard CD3/CD28 stimulation to determine whether T cell pre-activation was required for efficient gene transfer, as previously described.^38,43^

### Cytokine and Functional Assays

Secreted IFN-γ, IL-2, TNF-α, and Granzyme B were quantified by ELISA in CAR-T:target coculture supernatants and mouse serum, as previously described. ^38,43^ Intracellular Granzyme B was measured by flow cytometry after permeabilization and anti-Granzyme B staining. Similarly, T cell memory subsets were characterized using antibody panels as described previously. Further, cytotoxicity against luciferase-expressing MM.1S, MM.1R, Raji, and NALM6 target cells was assessed by bioluminescence killing assay at varying E:T ratios (4-24 h), as previously described. Similarly, cell death analysis by flow cytometry based 7-AAD staining was performed as described previously.^38,43^

### Biochemical and Receptor Validation Assays

#### STAT5 Phosphorylation Assay

Primary T cells (1 × 10^6^) were incubated with titrated sBCMA peptides for 30 m at 37°C, then fixed and permeabilized using Intracellular Fixation and Permeabilization Buffer Set (Cat no.: 88882400, Thermo Fisher, USA). Cells were stained with anti-phospho-STAT5 (Tyr694) antibody (Cat No.: MA5-14822, Thermo Fisher, USA) and isotype control (Cat no.: 12471482, Thermo Fisher, USA) for 60 m at 4°C. Analysis was performed by flow cytometry (BD FACS) using FlowJo v10 software, as previously described.^38,43^ Further, Primary T cells were stimulated with titrated recombinant soluble BCMA or sBCP3_min3-IL7R. Phospho-STAT5 (Tyr694) levels were quantified using the PathScan® Phospho-Stat5 Sandwich ELISA Kit (Cell Signaling Technology, #7113) per manufacturer’s instructions.

#### Receptor Competition and Binding Assays

Recombinant BCMA (BCMAR) and CD19 (CD19R) extracellular domain proteins (0.5-1 µg/mL) were pre-incubated with viroVbot particles for 1 h at 37°C, then used to transduce wild-type Raji and MM.1S cells. Transduction efficiency was quantified at 48 h post-infection by flow cytometry. The flow cytometry analysis confirmed that BCMAR or CD19R binding to the viroVbot1 which has BCMA scFv and CD19 scFv significantly reduced the uptake of these vectors.

#### Vector Copy Number Analysis

Vector copy number (VCN) per diploid genome was determined by quantitative PCR (qPCR) in peripheral blood, spleen, bone marrow, liver, lung, kidney, brain, and tumor tissue. Genomic DNA was extracted using DNeasy Blood & Tissue Kit (Qiagen) with on-column RNase A treatment. Each sample was run in technical triplicate with a standard curve generated from serial dilutions of CAR-positive reference DNA. VCN per diploid genome was calculated as 2 × (vector copies / reference gene copies).

### Animal Experiments

Animal experiments were conducted at Adgyl Lifesciences, Bengaluru, Karnataka, and at the CSIR-Institute of Chemical Biology, Kolkata, West Bengal, India, following institutional ethical guidelines and approval from the Institutional Animal Ethics Committee. Animal care and euthanasia procedures complied with the Committee for the Purpose of Control and Supervision of Experiments on Animals guidelines and institutional regulations.

As described by us previously,^38,43^ NOD.Cg-*Prkdc*^scid^ *Il2rg*^tm1Wjl^/SzJ (NSG) and NOD-*Prkdc*^em26Cd^^52^ *Il2rg*^em26Cd^^22^/NjuCrl (NCG) mice with or without MHC I/II KO mice, both male and female, aged 6 to 8 weeks, were obtained from The Jackson Laboratory and Charles River Laboratories respectively. BALB/c mice aged 6 to 8 weeks were sourced from institutional breeding colonies. All mice were housed in specific pathogen-free barrier facilities equipped with individually ventilated cages, high-efficiency particulate air filtration, 12 h light to dark cycles, and maintained at constant temperature (22°C) with ad libitum access to food and water. Experimental and control animals were co-housed whenever possible under identical husbandry and enrichment conditions. Cage changes and animal handling were performed in Class II biosafety laminar flow workstations to maintain aseptic conditions and minimize contamination. Mice were euthanized by carbon dioxide inhalation followed by cervical dislocation.

### Animal Ethics Statement

All animal experiments were approved by the Institutional Animal Ethics Committee (IAEC) of Adgyl Lifesciences, Bengaluru, Karnataka and CSIR-Institute of Chemical Biology, Kolkata, West Bengal, India under Approval No. AD-G3538 and ICB/IAEC/2025/October/9. The study conducted in accordance with CCSEA guidelines (Government of India) and ARRIVE 2.0 guidelines. Animal housing, husbandry, welfare monitoring, analgesia, anesthesia, and humane endpoints were implemented throughout the study to minimize pain and distress.

### Assessment of Neutralizing Antibody Response In Vivo

Female C57BL/6 mice (n = 12) were immunized subcutaneously with 10 µg of adjuvanted recombinant envelope glycoproteins (rEnv) from VSV-G, VSAV-G, COCV-G, or PIRYV formulated in Montanide ISA 51, followed by a booster on Day 14. On Day 28, mice received a subcutaneous challenge with lentiviral vector (LVV) pseudotyped with the corresponding envelope glycoprotein (1 × 10⁶ transduction units). Blood was collected on Day 56, and serum was isolated by centrifugation and stored at -20°C until analysis. This assay procedure was followed as previously described.^19^

Prior to neutralization testing, pooled serum samples underwent absorption with 1 × 10^7^ HEK293T cells per 150 µL serum on ice for 1 h to remove non-specific antibodies directed against host cell antigens. The absorbed serum was then subjected to serial dilution in plain OptiMEM medium, generating 12 concentrations ranging from 1:2 to 1:6,250,000. Each dilution (10 µL) was mixed 1:1 (v/v) with lentiviral vector pseudotypes at 4 × 10^5^ TU/mL, incubated at 37°C for 1 h to allow antibody-vector interaction, and subsequently applied to HEK293T cells seeded in 96-well plates at 2 × 10^4^ cells per well in 150 µL medium. Following 48 h of infection, cells were harvested by trypsin digestion and analyzed by flow cytometry to determine the percentage of green fluorescent protein-positive cells. Transduction efficiency at each serum dilution was calculated relative to vector-only controls, and the resulting data were plotted against reciprocal dilution factors using GraphPad Prism v10.3.

### Immunogenicity prediction and ex vivo immune screening

#### Ex vivo immune screening

Recombinant envelope proteins (VSNJG, VSV-G, COCV-G, PIRYV, VSAV-G, and others) were produced in HEK293T cells and purified. Peripheral blood was collected from 65 healthy donors (informed consent, ethical approval obtained) and PBMCs were isolated by Ficoll-Paque density gradient and cryopreserved. Cryopreserved PBMCs were thawed, stimulated with 4 μg/mL recombinant envelope protein or PBS control for 18 h at 37 °C in the presence of anti-CD40 (2 μg/mL), then fixed and analysed by flow cytometry on a BD FACS Lyric. Antigen-specific CD4⁺ T cell responses were quantified using CD154 expression, and CD8⁺ T cell responses using CD137 expression, with memory subsets identified by CCR7/CD45RA (T_em_ and T_cm_). Data were analysed in FlowJo v10 and plotted as heatmaps (Prism v9).

#### In vitro infectivity and cytotoxicity assays

Wild-type and computationally-optimised envelope variants were tested in infectivity assays using pseudotyped viral particles at serially diluted relative difference factor (RDF) values (1 × 10^5^ to 1 × 10⁰) on HEK293T target cells expressing cognate receptors, with infectivity quantified by luciferase activity. Killing capacity of envelope-specific CD8⁺ T cells was assessed by co-culture of flow-cytometry-sorted CD137⁺ CD8⁺ T cells with infected HEK293T targets at an effector-to-target ratio of 10:1, with target cell viability monitored over 48-72 h. Cytokine production (IFN-γ, TNF-α) was measured in supernatants and sera by multiplex Luminex assay.

#### Antigen-specific T cell

Antigen-specific CD4⁺ and CD8⁺ T cell responses were quantified by flow cytometry as the percentage of CD154⁺ and CD137⁺ cells, respectively, relative to PBS control wells. Statistical significance was determined between stimulated and unstimulated groups. The protocol for these assays was followed as mentioned previously.^36^

### Humanized Mouse Models

To establish disseminated multiple myeloma, MM.1S-luciferase cells (1 × 10^6^ cells per mouse) were injected via the lateral tail vein on Day -4. Health donor-derived human PBMCs (1 × 10^7^ cells per mouse) were engrafted intravenously on Day -5 (one day before tumour inoculation). On Day 0, mice were randomised by baseline tumour burden (bioluminescence imaging, BLI) and were treated with a single intravenous dose of viroVbot2.1 or viroVbot2.2, or with vehicle control (*Figure 7*). Tumour burden was monitored by serial whole-body BLI (IVIS Spectrum, PerkinElmer) on Days 7, 14, 21, 28, and 56, with peripheral blood sampled on the same days for flow-cytometric quantification of circulating MM.1S cells, total human CD45⁺ chimerism, and CAR⁺ T cell frequency among CD3⁺ cells. At Day 56, animals were sacrificed and bone marrow, spleen, liver, lungs, and kidneys were harvested for end-point flow cytometry, immunohistochemistry, and vector-copy-number analysis. viroVbot vectors were administered intravenously via tail vein injection in a total volume of 200 μL

Haematological relapse/antigen-escape model (*Figure 7*). To model sequential antigen escape, MM.1S-luciferase cells (1 × 10^6^ cells per mouse) were injected intravenously on Day -7, followed by intravenous engraftment of human PBMCs (1 × 10⁷ cells per mouse) on Day -5. Mice received a single intravenous dose of viroVbot3 (0.25 × 10^6^) on Day 0 (Group 2). Subsequent groups received an antigen-loss tumour rechallenge with GPRC5D⁺/BCMA⁻/CD19⁻ MM.1S cells on Day 45 (Group 3, “TR”), with two further treatment arms receiving either viroVbot-CO (Group 4) or viroVbot-VS (Group 5) intravenously on Day 47. BLI and peripheral blood flow cytometry (for BCMA⁺ and GPRC5D⁺ CAR T cells) were performed every 10 days through Day 90, with terminal collection of bone marrow and spleen for CAR T phenotyping.

*CD34^+^ HSC-humanised model for vector dose response and B cell aplasia (Figure 3)*. Busulfan treated (25 mg/kg/day x 2 injected intraperitoneally) NSG mice were engrafted by intrahepatic injection with conditioned whole blood-derived CD34⁺ haematopoietic stem cells (5 × 10^5^ cells per mouse). Animals were aged for 28 days to allow multi-lineage reconstitution (HSCs delivered at Day -28 relative to vector administration), with a baseline blood draw at Day -4 to confirm human CD45⁺, CD3⁺, and CD19⁺ engraftment. On Day 0, mice received a single intravenous dose of viroVbot1 at one of three dose levels (0.25 × 10^6^, 0.5 × 10^6^, or 1 × 10^6^ transducing units per mouse) or ePIRYV^wt^ control vector. Peripheral blood was sampled on Day -4, 7, 14, 21, and 28 for serial quantification of BCMA-CAR⁺ CD3⁺ T cells and CD19⁺ B cell depletion among hCD45⁺ cells. Terminal bone marrow and spleen were collected at Day 28 for end-point CAR-T enumeration and confirmation of B cell aplasia, and biodistribution was assessed in liver, spleen, lungs, and kidneys by immunohistochemistry and vector copy number.

#### Solid tumour model

PBMC-humanised AGS gastric adenocarcinoma model with sequential antigen-switching re-treatment (*Figure 8*). To evaluate viroVbot activity against a CLDN18.2⁺ solid tumour, NCG mice were inoculated intraperitoneally with luciferase-labelled AGS human gastric adenocarcinoma cells as described by us previously.^38^ Two days later (Day -5), donor-derived human PBMCs (1 × 10^7^ cells per mouse) were engrafted intravenously to provide the effector T cell substrate for in situ viroVbot transduction. On Day 0, mice were randomised by baseline BLI tumour burden and received a single intravenous dose of viroVbot3S (1x 10^6^; encoding the PY-CL18.2 CAR architecture).

To model the clinical reality of relapse and to test sequential antigen-switching, a re-treatment regimen was applied to surviving animals: on Day 31, MM.1S-luciferase cells were re-introduced intravenously as a secondary tumour challenge to evaluate cross-protection; PBMCs were re-engrafted on Day 36 (1 × 10⁷ cells per mouse); and viroVbot3.1S (CO-CL18.2) was administered intravenously on Day 38. A third cycle was performed on Days 61/66/68 with MM.1S rechallenge, PBMC re-engraftment (1 × 10⁷ cells), and viroVbot3.2S (VS-CL18.2) administration, respectively. Tumour burden was monitored by serial BLI and by external caliper measurement of any palpable tumour mass (volume calculated as length × width² × 0.5) on Days 10, 20, 30, 60, and 90. Peripheral blood was sampled at each BLI time point through Day 90 for flow-cytometric quantification of BCMA⁺ and CLDN18.2-targeted CAR -T cell kinetics among hCD45⁺ CD3⁺ cells. Body weight was recorded twice weekly throughout the study as a global tolerability readout, and any animal losing >20% of its baseline body weight or meeting predefined humane endpoint criteria (hunched posture, ruffled fur, reduced mobility, signs of xenogeneic graft-versus-host disease) was euthanised. At terminal sacrifice (Day 90 for surviving animals, or at humane endpoint), spleen, bone marrow, liver, kidneys, lungs, and stomach were harvested for histopathology, with formalin-fixed paraffin-embedded sections processed for haematoxylin and eosin staining to evaluate tumour architecture, lymphocytic infiltration, and the integrity of normal gastric epithelium adjacent to AGS lesions.

### CAR -T Cell Analysis and Tissue Distribution

Peripheral blood was collected at study-specific intervals (Days 0, 7, 14, 21, 28 for Fig. 3; Days 7, 14, 21, 28, 56 for Fig. 5; Days 10-90 every 10 days for Figs. 7 and 8) by submandibular or retro-orbital sampling under brief isoflurane anaesthesia into K₂EDTA tubes, and erythrocytes were removed using ammonium chloride lysis buffer (BD Pharm Lyse).

CAR surface expression was quantified by flow cytometry using either antigen-specific biotinylated detection reagents (anti-CD19 CAR, Miltenyi #130-129-550; anti-BCMA CAR, Miltenyi #130-127-348) with streptavidin-fluorophore secondary staining, or, for antigen-agnostic detection across all viroVbot constructs including the GPRC5D- and Claudin-18.2-targeted CARs; the anti-G4S linker rabbit monoclonal antibody clone E7O2V (Cell Signaling Technology, unconjugated #71645, Alexa Fluor 488 conjugate #50515, or PE conjugate #38907), which recognises the (Gly_4_Ser)_3_ spacer common to all scFv-based viroVbot CARs. Cells were Fc-blocked (BD Biosciences), stained with a Fixable Viability Staining Assay (ab314684, Abcam) and a multicolour panel for human CD45, CD3, CD4, CD8, CD19, CD20, CD56, CD14, CD15 (with murine CD45 to exclude host cells), acquired on flow cytometer. CAR⁺ cells were reported as a percentage of hCD45⁺CD3⁺ T cells, with absolute counts derived using CountBright beads (Invitrogen) where indicated. All staining, gating, and data-analysis workflows followed protocols previously published by our group.^38,43^ At the end of the study, peripheral blood, and bone marrow were harvested, and the bone marrow were processed as described by us previously,^37^ and processed into single-cell suspensions for flow cytometric analysis using the same panel as above.

For immunohistochemistry (IHC), organs were fixed in 10% neutral-buffered formalin, paraffin-embedded, and sectioned at 5 μm. Following heat-induced epitope retrieval (citrate pH 6.0 or EDTA pH 9.0), endogenous peroxidase quenching, and protein blocking, sections were probed overnight at 4 °C with an anti-G4S linker rabbit monoclonal antibody (CST #71645, 1:200) and detected using an anti-rabbit HRP-conjugated secondary antibody (CST #8114). DAB was used as the chromogenic substrate, sections were counterstained with Mayer’s haematoxylin, and images were acquired by bright-field microscopy at 20× and 40× magnification, as described previously.^65^ Quantitative analysis of the IHC signal was performed as previously described.^65^ Tumour sections were additionally stained with H&E to assess tissue architecture and lymphocytic infiltration. CAR⁺ cell density was quantified across ≥5 random, non-overlapping high-power fields per section.

### Bioluminescence Imaging

Baseline tumor burden was established via bioluminescence imaging of luciferase-expressing tumor cells using the IVIS Spectrum imaging system, as described previously.^38,43^ Mice were imaged as following intraperitoneal injection of D-luciferin (150 milligrams per kilogram) after 10 to 15 m incubation under isoflurane anesthesia. Total flux was quantified using Living Image software and normalized to background. Complete blood counts were performed at baseline and terminal time points using automated hematology analyzers. Tumor burden was assessed at indicated intervals to evaluate in vivo CAR-T cell anti-tumor activity.

### Tumor Rechallenge (In vitro and In Vivo)

For in vitro rechallenge, CAR-T cells were repeatedly stimulated with fresh tumour targets every 2 days over 15-21 days or as indicated, with CAR-T persistence, target lysis, and cytokine secretion measured at each round, as described previously.^38,43^

For in vivo rechallenge (*Figure 7*), antigen-escape tumour relapse model was developed by intravenously injecting MM.1S^GPRC5D+/BCMA−/CD^^19−^ cells (1 × 10⁶ per mouse) on Day 45, which is 45 days after the initial viroVbot3 (BCMA/CD19 bi-CAR) dose, into surviving mice from groups previously treated with viroVbot3. Two days later (Day 47), donor-matched human PBMCs (1 × 10⁷ per mouse) were re-engrafted intravenously to refresh the human T cell pool, and on Day 50 mice received a single intravenous dose of either viroVbot-CO or viroVbot-VS (both encoding a GPRC5D-targeted CAR) to test sequential antigen-switching. Tumour burden (BLI) and circulating GPRC5D-CAR⁺ T cells were monitored every 10 days through Day 90, when terminal bone marrow and spleen were collected for end-point flow cytometry.

### Flow Cytometry Analysis

Cells were analyzed on BD FACS Aria, Accuri, Lyric, or Beckman Coulter CytoFLEX instruments and data analyzed using FlowJo v10 or CytExpert software as described previously.^38,43^ CAR transgene expression was detected using CAR-specific detection reagents: CD19 CAR detection (Miltenyi Biotec #130-129-550), BCMA CAR detection (Miltenyi Biotec #130-126-727), anti-4-1BB monoclonal antibodies for 4-1BB-containing constructs, anti-G4S linker antibody for GPRC5D (#62405, Cell Signaling Technologies) and CLDN18.2 CARs, or Protein L-APC (CST #29480) for CD20 CAR. T cell immunophenotyping used fluorophore-conjugated antibodies against CD3, CD4, CD8, CD27, CCR7, CD45RA, and CD62L to identify memory subsets and exhaustion markers, and off-target markers CD45, CD19, CD20, CD14, CD15, CD56 were assessed, as mentioned previously.^38,43^ Intracellular staining for Granzyme B (clone GB11, BioLegend) and phospho-STAT5 (Tyr694; Cat no. 12901042, Thermo Fisher) was performed after fixation/permeabilization (Intracellular Fixation and Permeabilization Buffer Set, Cat no. 88882400, Thermo Fisher). Cell viability was determined using 7-AAD staining (BD Biosciences #559925). Gating strategy: lymphocyte singlets; live (7-AAD^-^); CD3+ T cells; CD4^+^/CD8^+^ subsets; CAR^+^ populations. Absolute counts were calculated from frequency data and hemocytometer counts. Complete antibody details are in **Table 9**. Previous protocols.^38,43^

### Human Samples

Peripheral blood mononuclear cells were obtained from two donor populations as described by us previously.^37,38^ Healthy adult volunteers (45 males and 27 females; aged 20 to 55 years) were recruited at Apollo Indraprastha Hospital, New Delhi. These healthy donor samples were used for ex vivo immune screening assays to assess pre-existing immune responses against viral envelope proteins and to evaluate baseline T cell activation profiles. Additionally, peripheral blood mononuclear cells were collected from 37 patients with multiple myeloma. Patient-derived peripheral blood mononuclear cells were used for multiple experimental purposes including evaluation of lentiviral vector transduction efficiency, assessment of CAR-T cell functionality in patient-derived cells, and generation of patient-derived CD3^+^ T cell populations for tumor-specific cytotoxicity assays. Sample collection and processing were conducted in accordance with the Declaration of Helsinki and approved by the Institutional Review Board of the Apollo Hospitals Educational and Research Foundation (IRB Application No. IAH-BMR-064/07-23).

Fresh bone marrow aspirates were also obtained from patients with multiple myeloma for isolation of patient-derived primary multiple myeloma cells. These cells were enriched for plasma cells via negative selection immunomagnetic enrichment and subsequently cultured on HS-5 human bone marrow stromal cell layers in RPMI-1640 medium supplemented with 10 percent fetal bovine serum, 2 millimolar L-glutamine, human cytokines (IL-6, BAFF, and APRIL), and appropriate antibiotics. Patient-derived multiple myeloma cells were used as target cells in cytotoxicity assays, tumor rechallenge studies, and for ex vivo functional evaluation of CAR-T cell responses. All participants provided written informed consent prior to sample collection. Blood and bone marrow samples were surplus specimens that would otherwise have been discarded after routine clinical testing or diagnostic procedures. Cryopreserved peripheral blood mononuclear cells were thawed prior to experimental use and assessed for viability by trypan blue exclusion. All procedures adhered to institutional ethical and regulatory standards and were performed under the supervision of qualified clinical personnel.

### STATISTICAL ANALYSIS

All statistical analyses were performed using GraphPad Prism (v9 and v10.3; GraphPad Software), which was also used to generate bar charts and P-values. Data are presented as mean ± SEM unless otherwise stated. Significance is denoted as *P < 0.05, **P < 0.01, ***P < 0.001, and ****P < 0.0001. Two-group comparisons were performed using non-parametric t-tests, and multi-group comparisons using non-parametric one-way or two-way ANOVA. Survival was analyzed using the Mantel-Cox (log-rank) test. All experiments included positive (as indicated) and vehicle (negative) controls, with a minimum of three biological replicates, and were independently repeated at least three times.

## Data Availability

Source data and processed datasets are provided in the Supplementary Information and Supplementary Tables. De-identified clinical sample data are available from the corresponding authors upon reasonable request, subject to institutional review board approval and a data use agreement. All unique reagents, cell lines, and plasmids generated in this study are available from the corresponding authors upon request.

## Code Availability

All computational scripts, AI-based prediction algorithms (CIMMEXA immunogenicity platform), and analysis workflows are available and will be uploaded in the GitHub repository. Custom Python and R scripts for data analysis, statistical analysis, and visualization are provided with complete documentation. Additional code supporting analyses is available from the corresponding authors upon request.

## Acknowledgements

The authors gratefully acknowledge the support of all members of the TA laboratory. We thank the team at Sarvodaya Hospital, Faridabad, for their assistance in arranging patient samples, and Dr. Garima Nirmal for her invaluable assistance in obtaining human samples. We are deeply grateful to the staff of CSIR-Indian Institute of Chemical Biology (CSIR-IICB), Kolkata, for their essential support in preclinical studies, and the team at Adgyl Lifesciences, Hyderabad, India, for their dedicated assistance with animal studies. The authors also acknowledge the valuable contributions of Dr. Vikram Mathews and Dr. Dinesh Pendharkar for providing critical clinical insights into disease pathogenesis, and Dr. Harenath K Pillai and Dr. Murugaiyan Nedunchezhian for their instrumental work in critical assay development during manuscript preparation. We appreciate the support of the confocal imaging staff at the Central Instrumentation Facility, University of Delhi South Campus, for their technical expertise. We acknowledge the use of Claude for grammar and language refinement during manuscript preparation; however, all scientific content, analyses, and interpretations were generated and verified by the authors. Multiple patent applications related to this work have been filed (202511034051; 202611069689).

## Author Contributions Statement

T.A., R.A., conceived the study, designed experiments. T.A., R.A., and A.I. supervised the research, and wrote the manuscript. D.M. developed the PromoterForge synthetic promoter platform. S.S. designed and validated the bispecific BCMA/CD19 CAR construct. V.C. developed soluble BCMA-targeting peptide binders and chimeric signaling receptors. S.K., J.G., A.S., and N.C. developed humanized nanobodies and contributed to CIMMEXA optimization. S.K., R.S., N.C., and K.H. performed computational immunogenicity predictions and CIMMEXA data analysis. A.K.S., R.C., and U.M. conducted preclinical in vivo efficacy studies and vector validation. M.I.M. assisted in preclinical studies. S.M., M.R., K.H., N.S., D.J., J.H., and S.A. performed in vitro experiments and lentiviral vector development. M.H. contributed to experimental design and data interpretation. P.I. and V.H. provided clinical insights and patient sample context. G.K. selected clinical samples and wrote clinical aspects of the manuscript. S.R. designed lentiviral vectors and performed CRISPR-Cas9 genome editing. All authors reviewed and approved the final manuscript.

