## Supplementary material for "Safe Redosable Low-Immunogenic In Vivo CAR-T Therapy for B Cell Malignancies and Solid Tumors": Supplementary text v2.docx

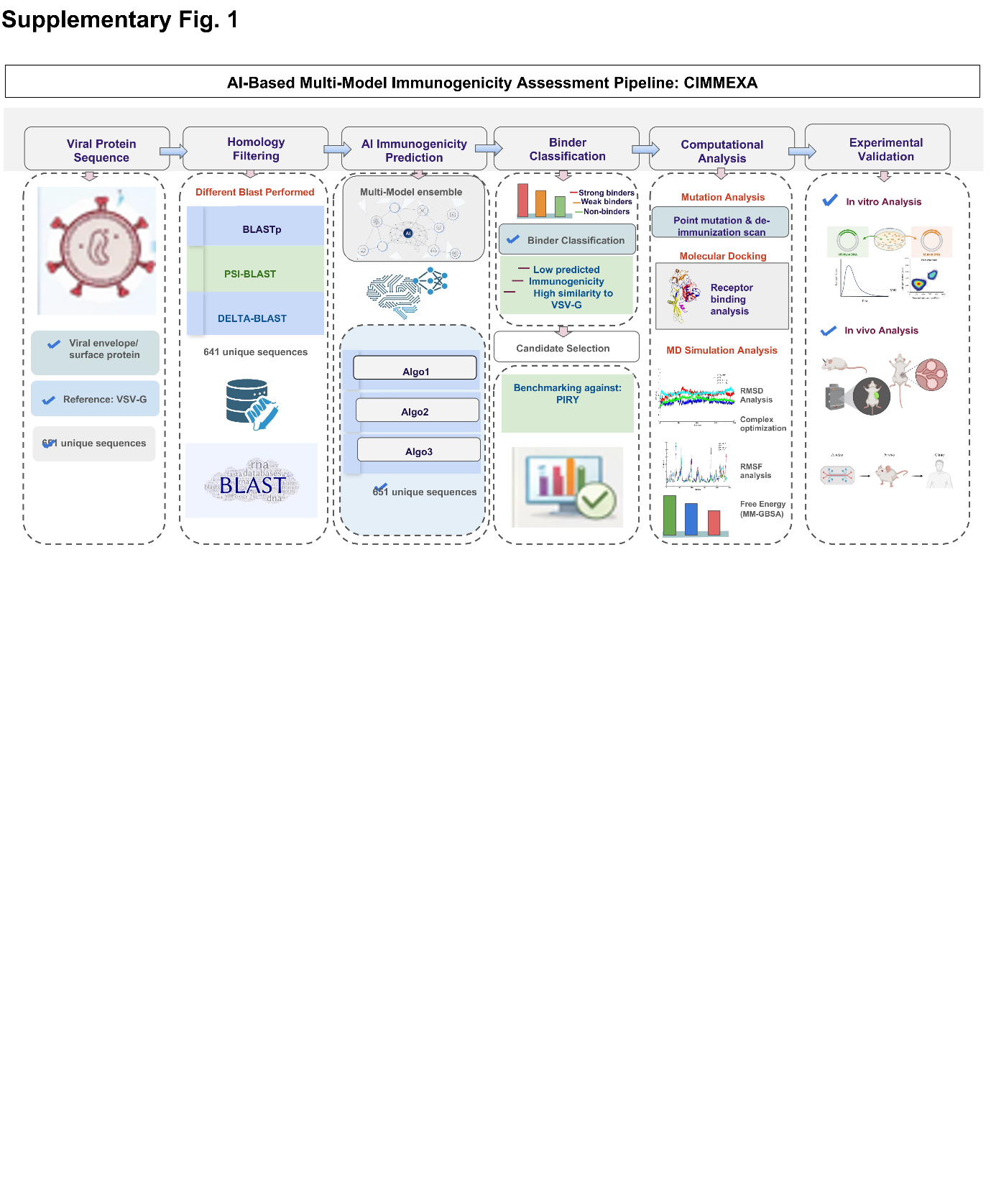

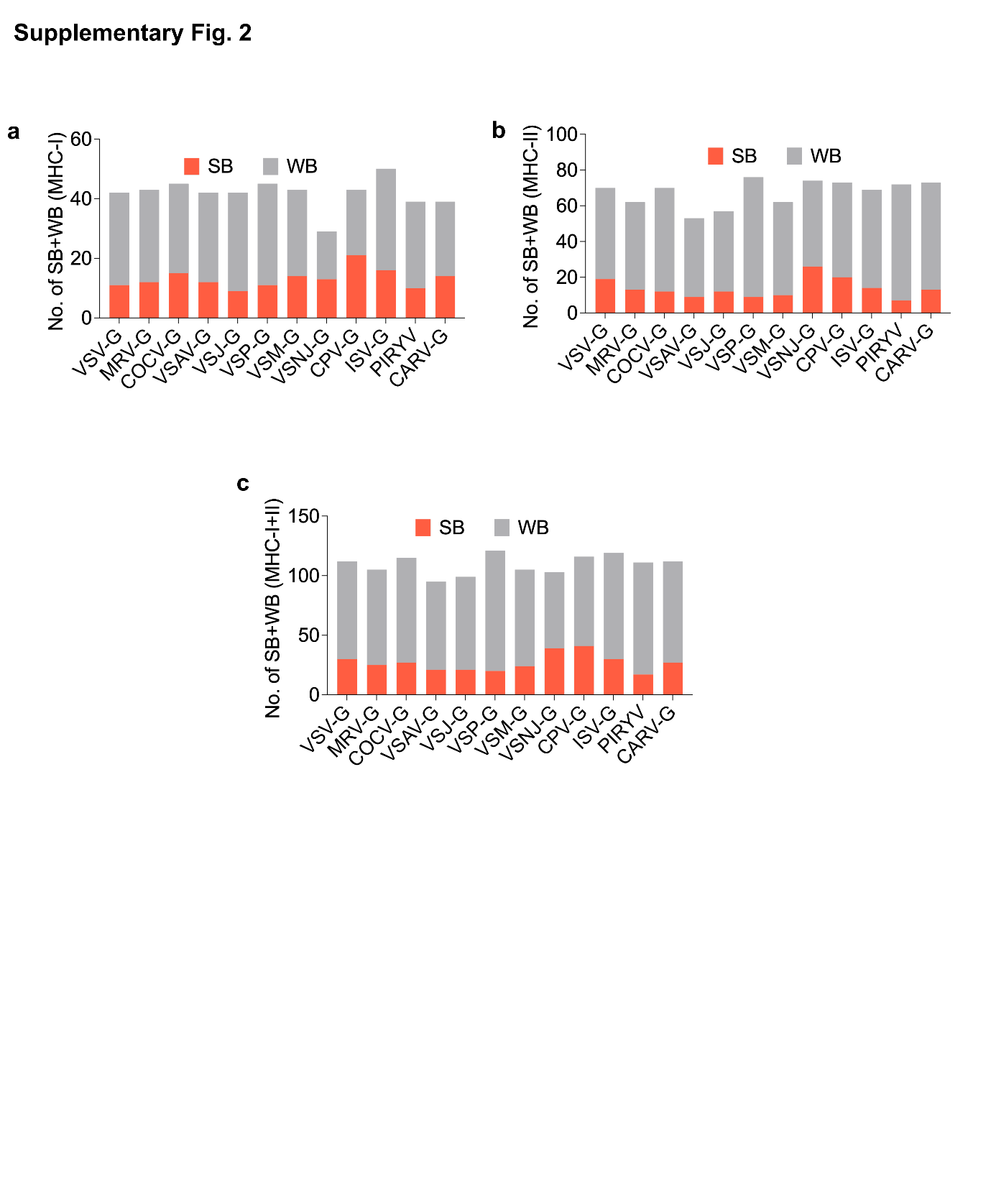

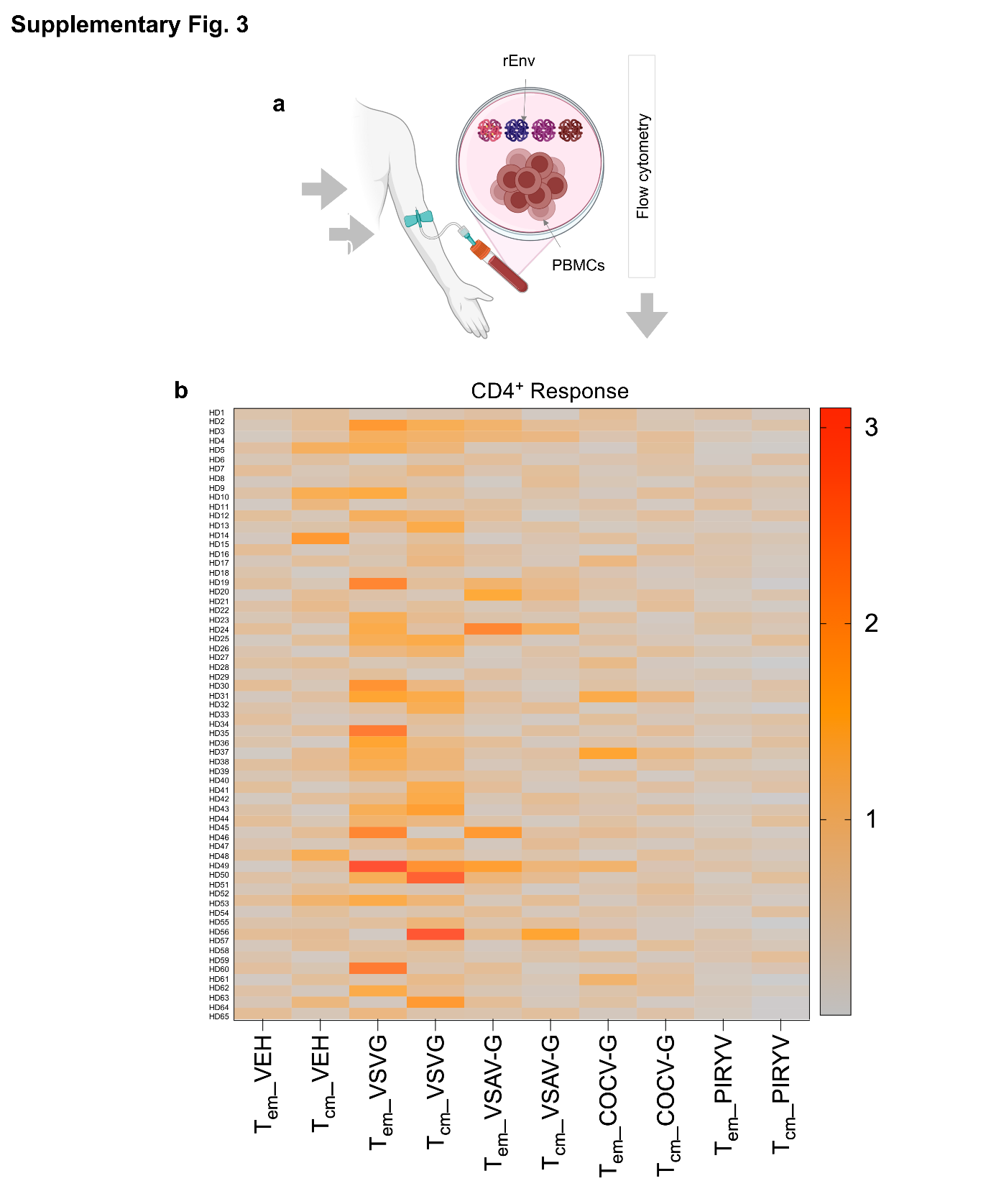

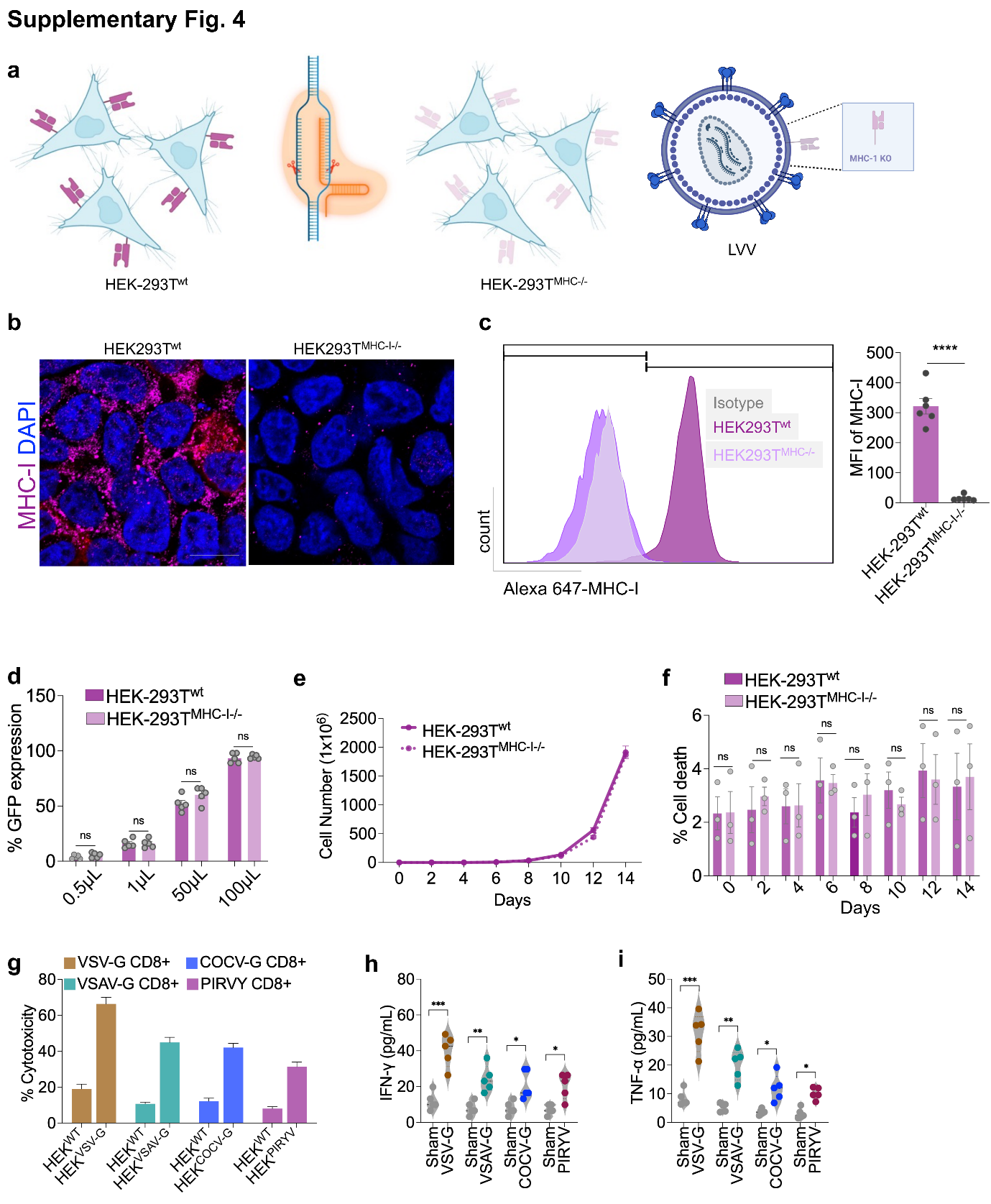

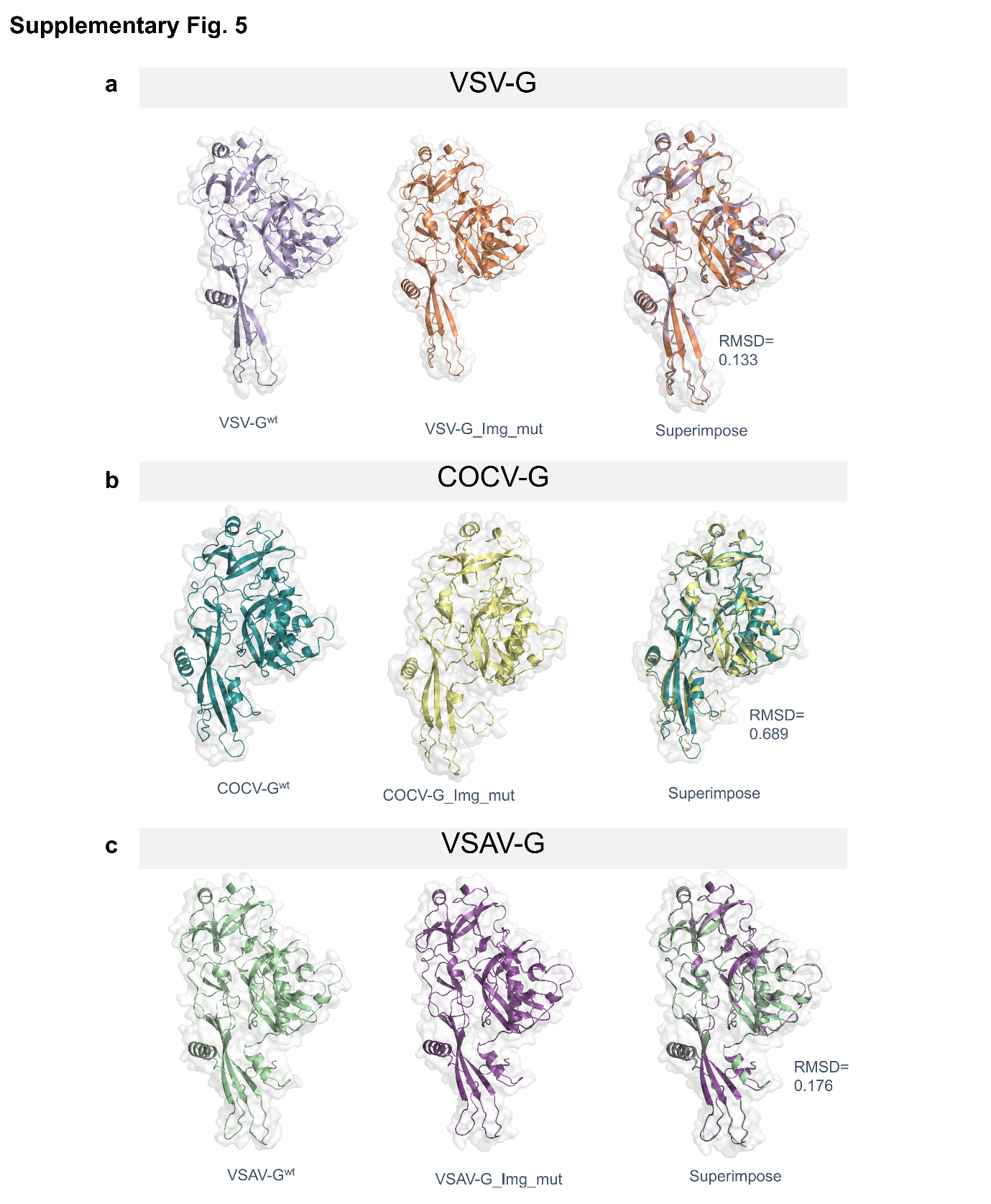

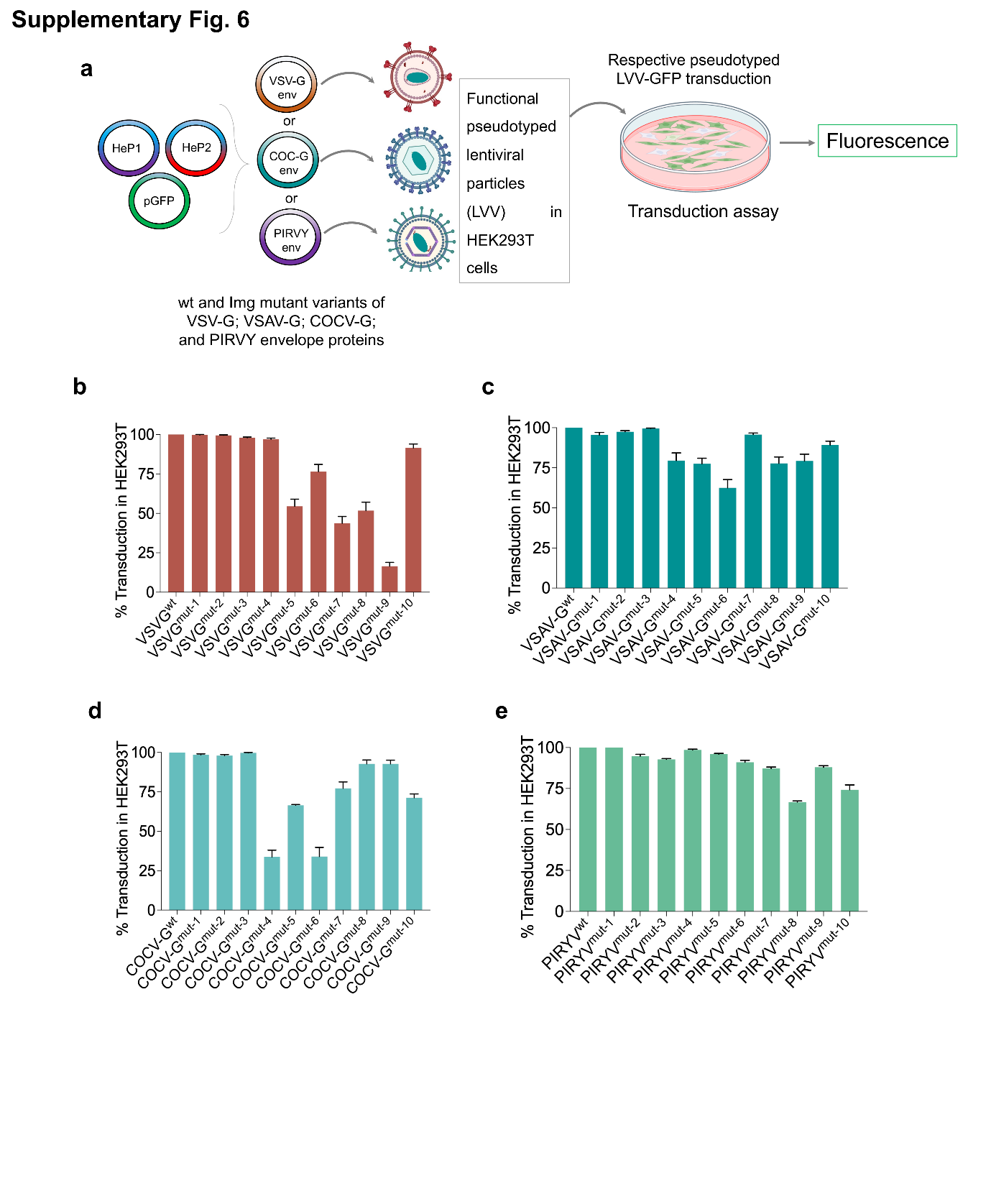

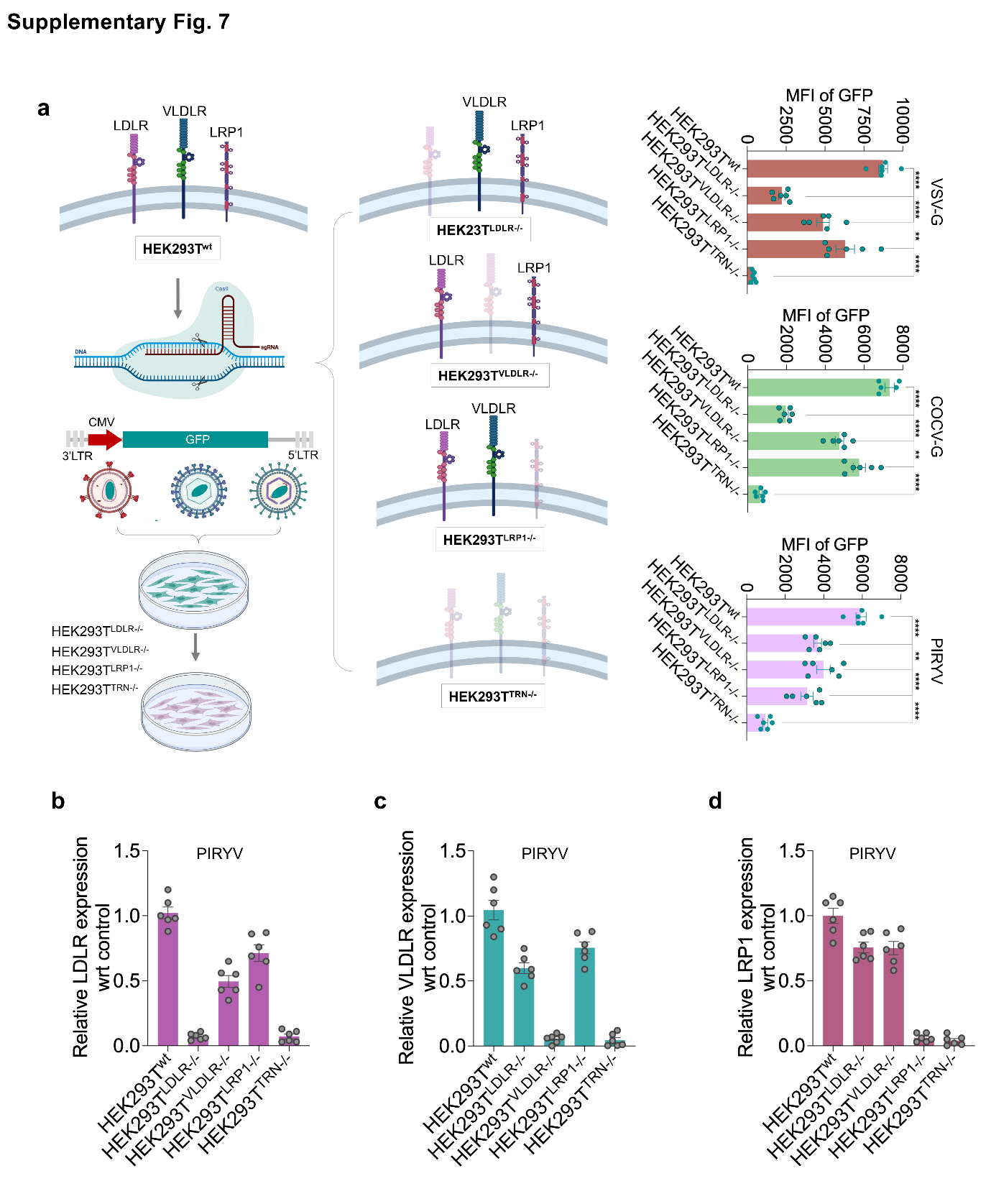

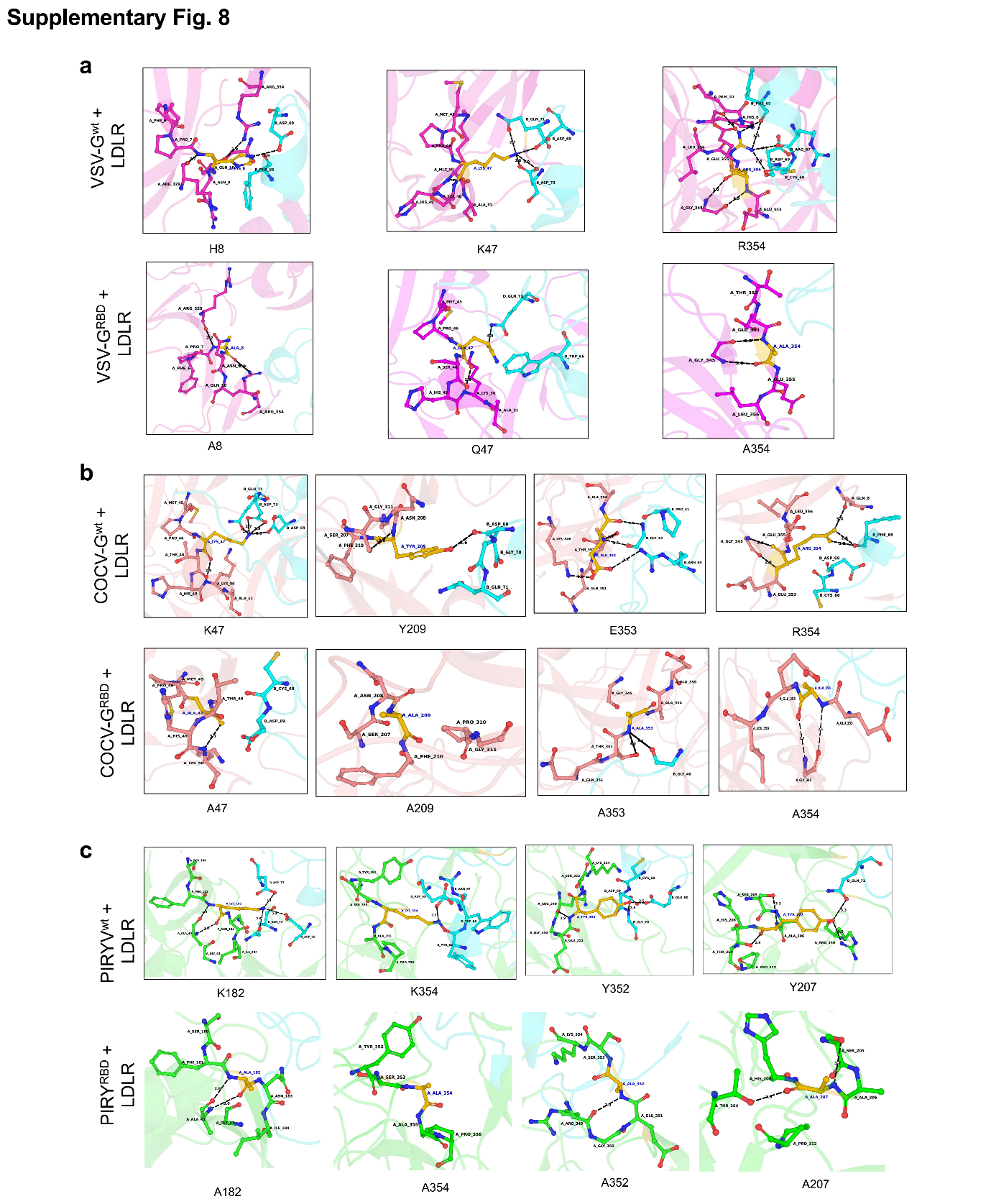

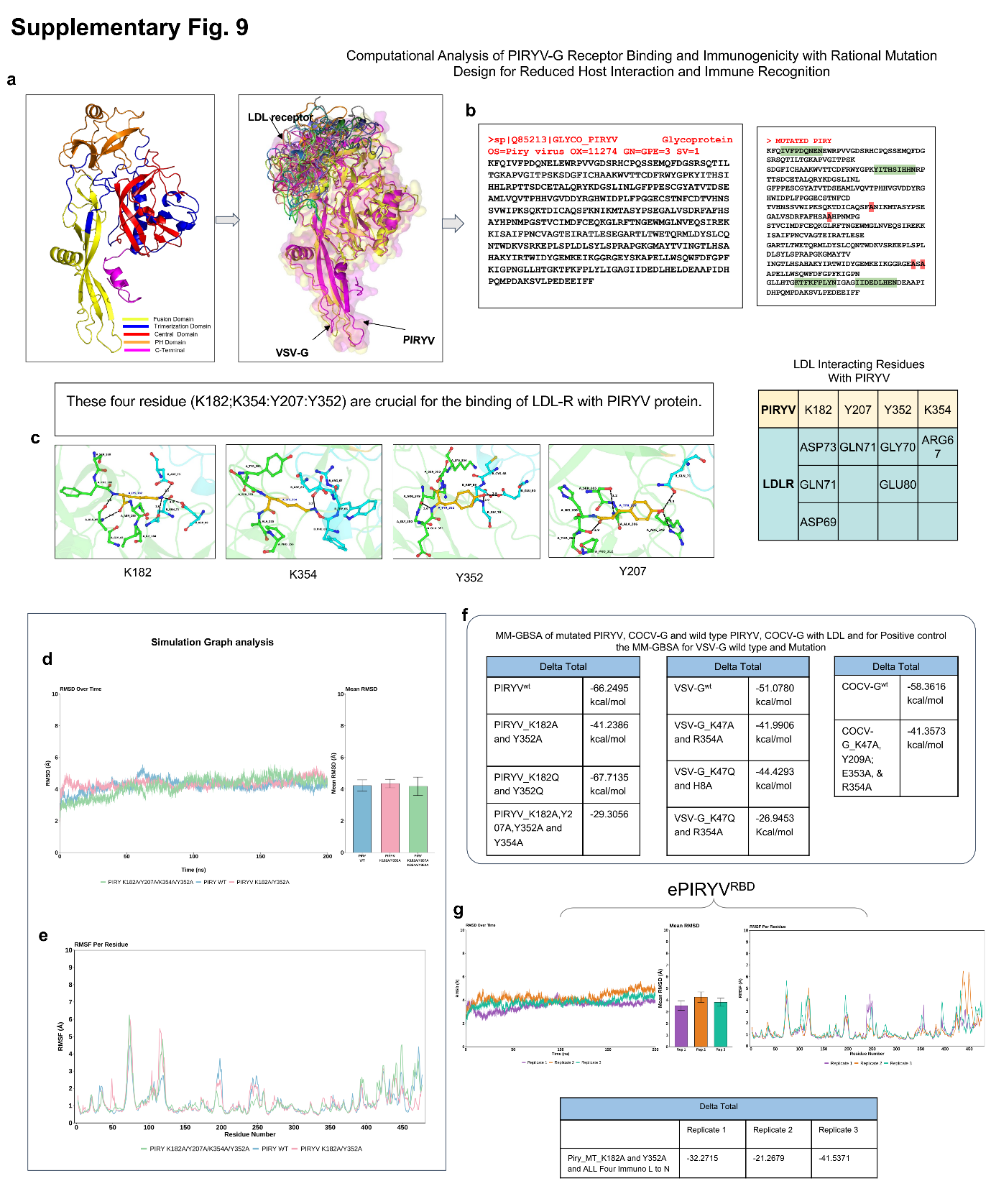

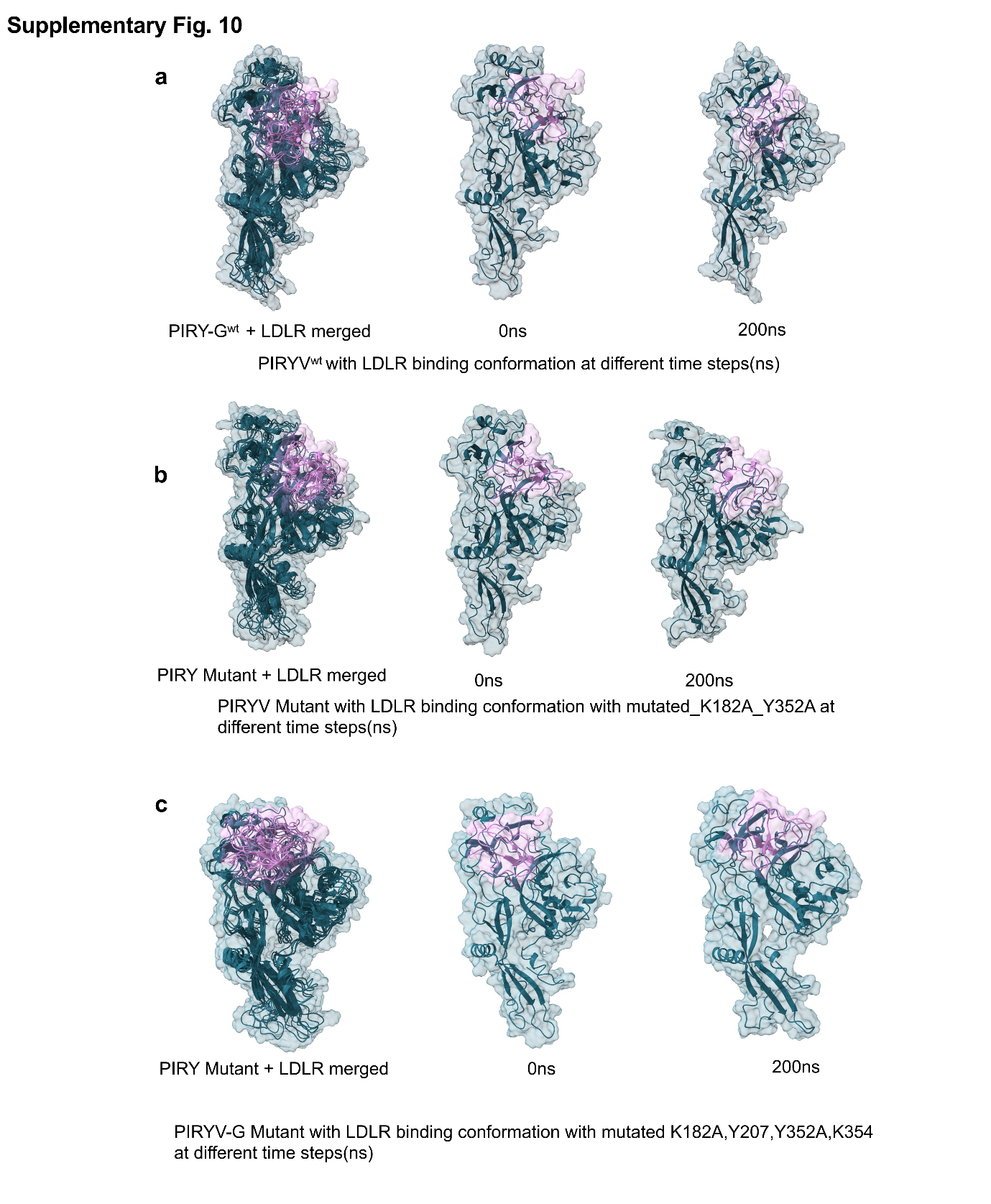

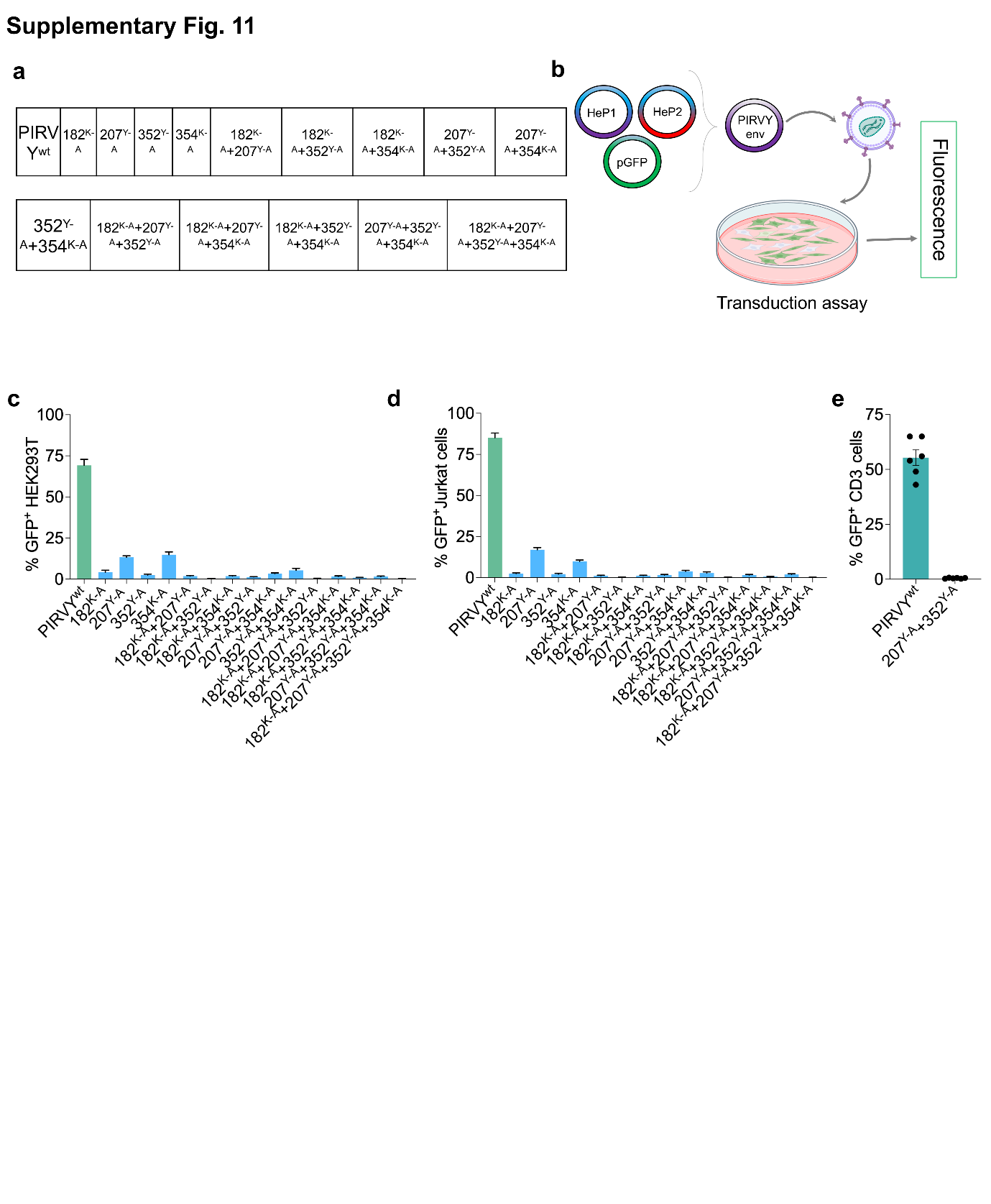

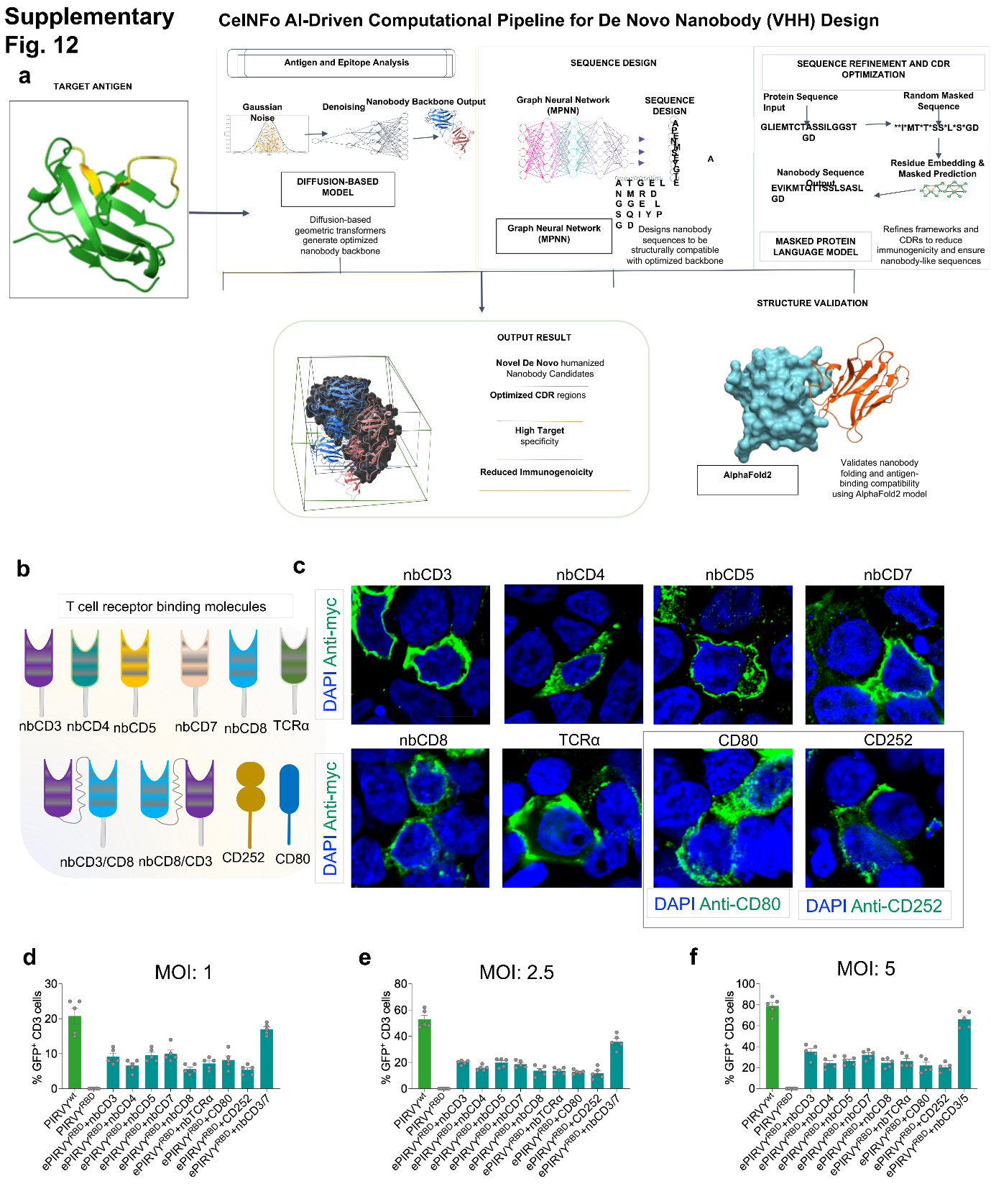

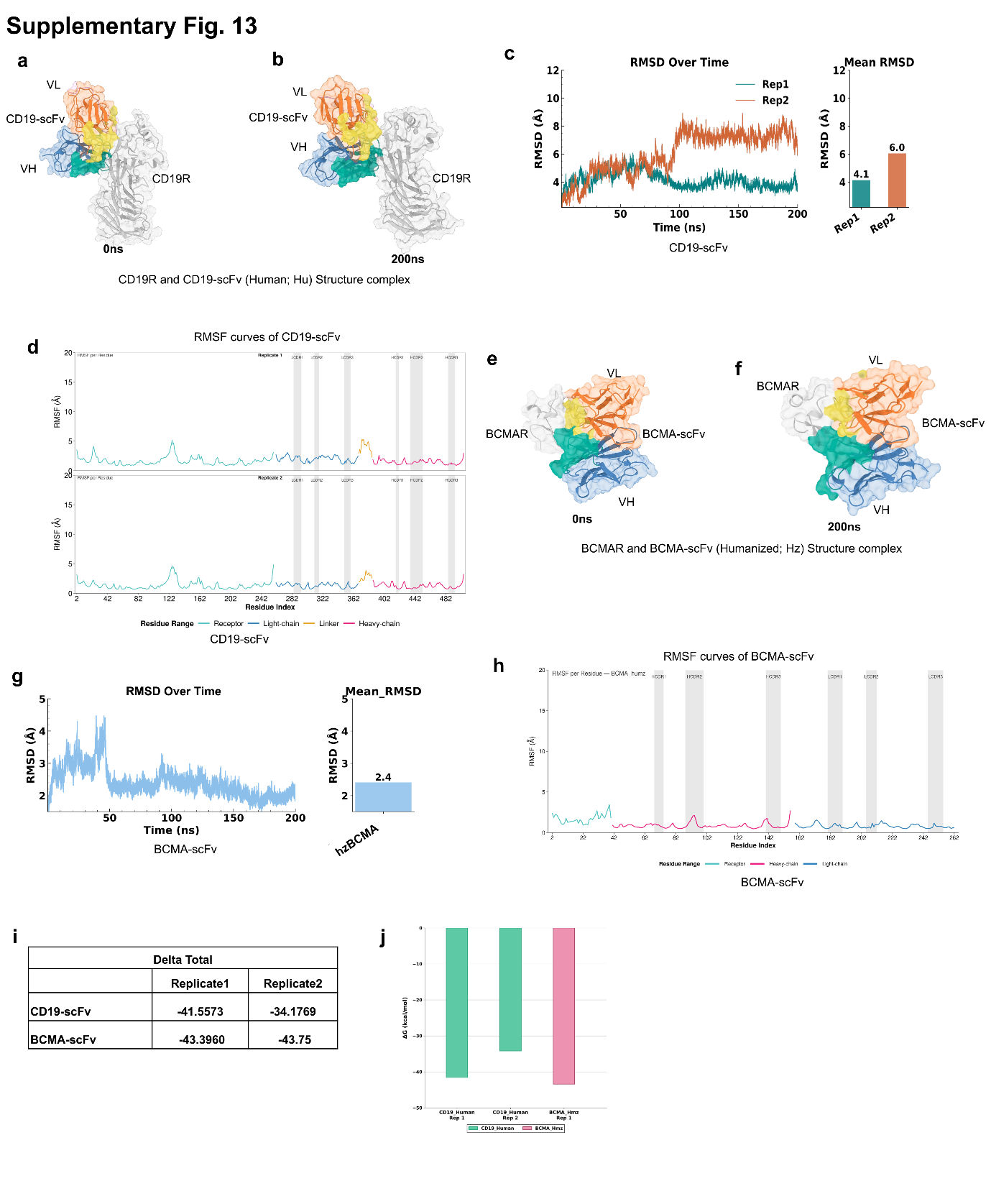

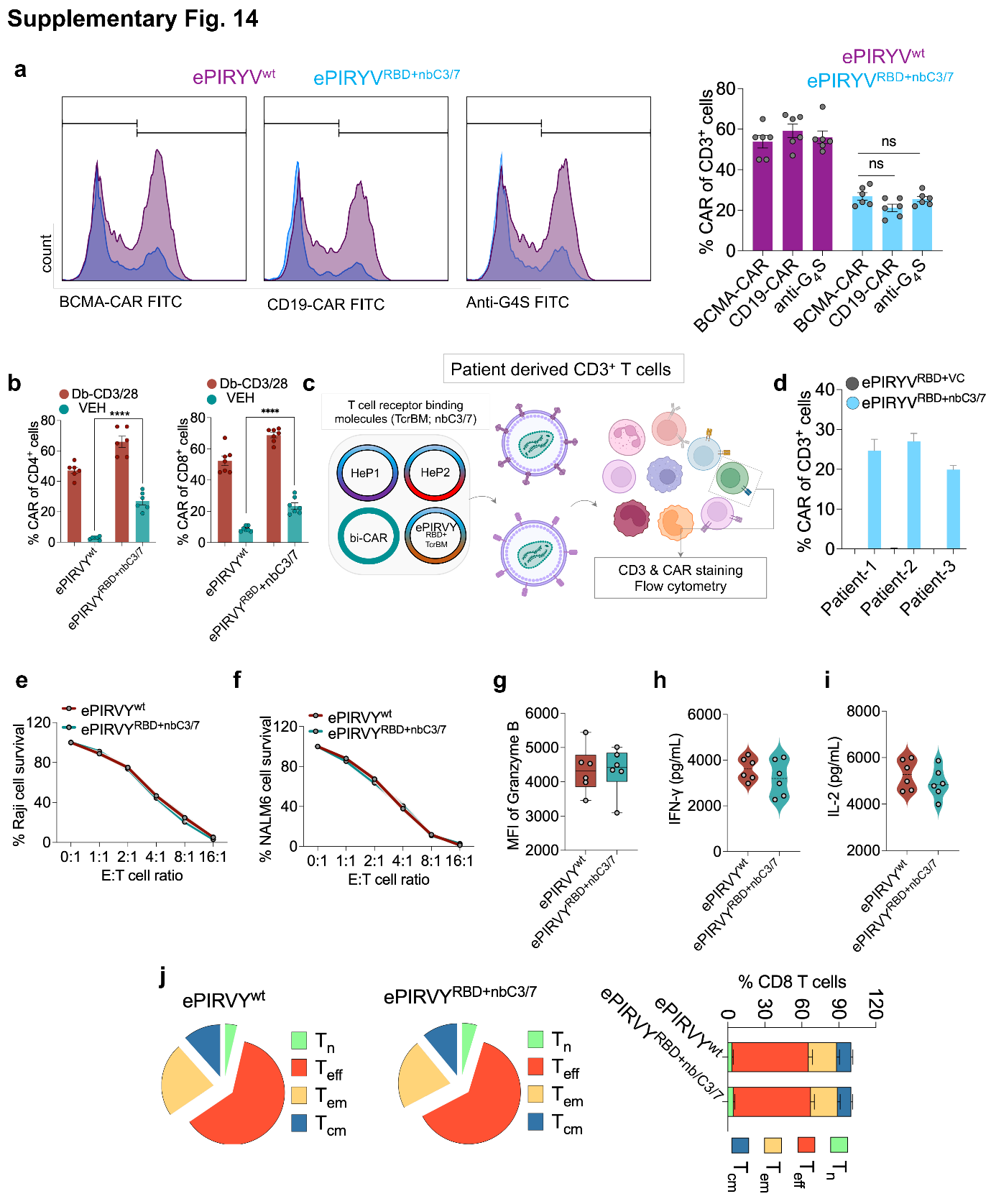

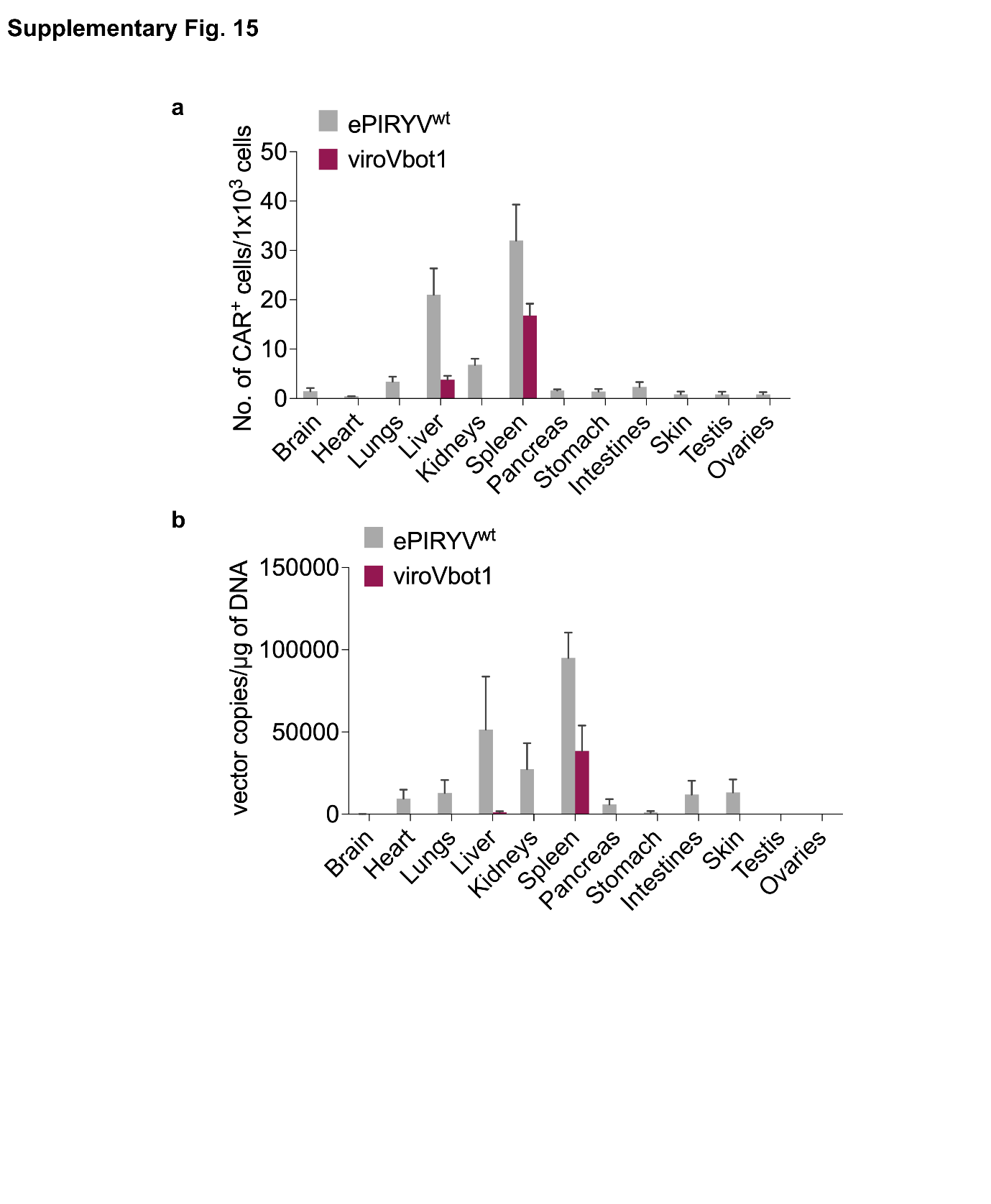

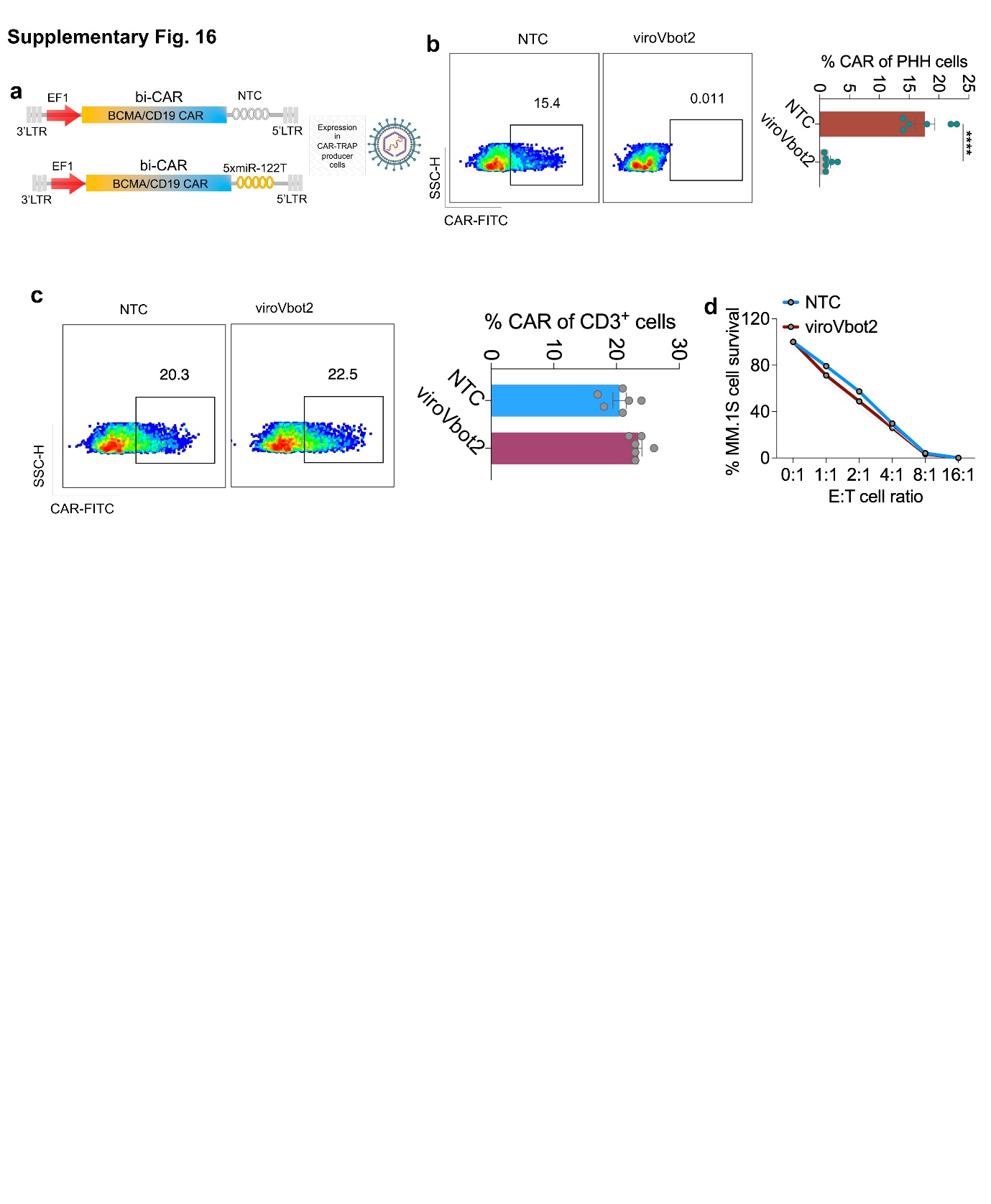

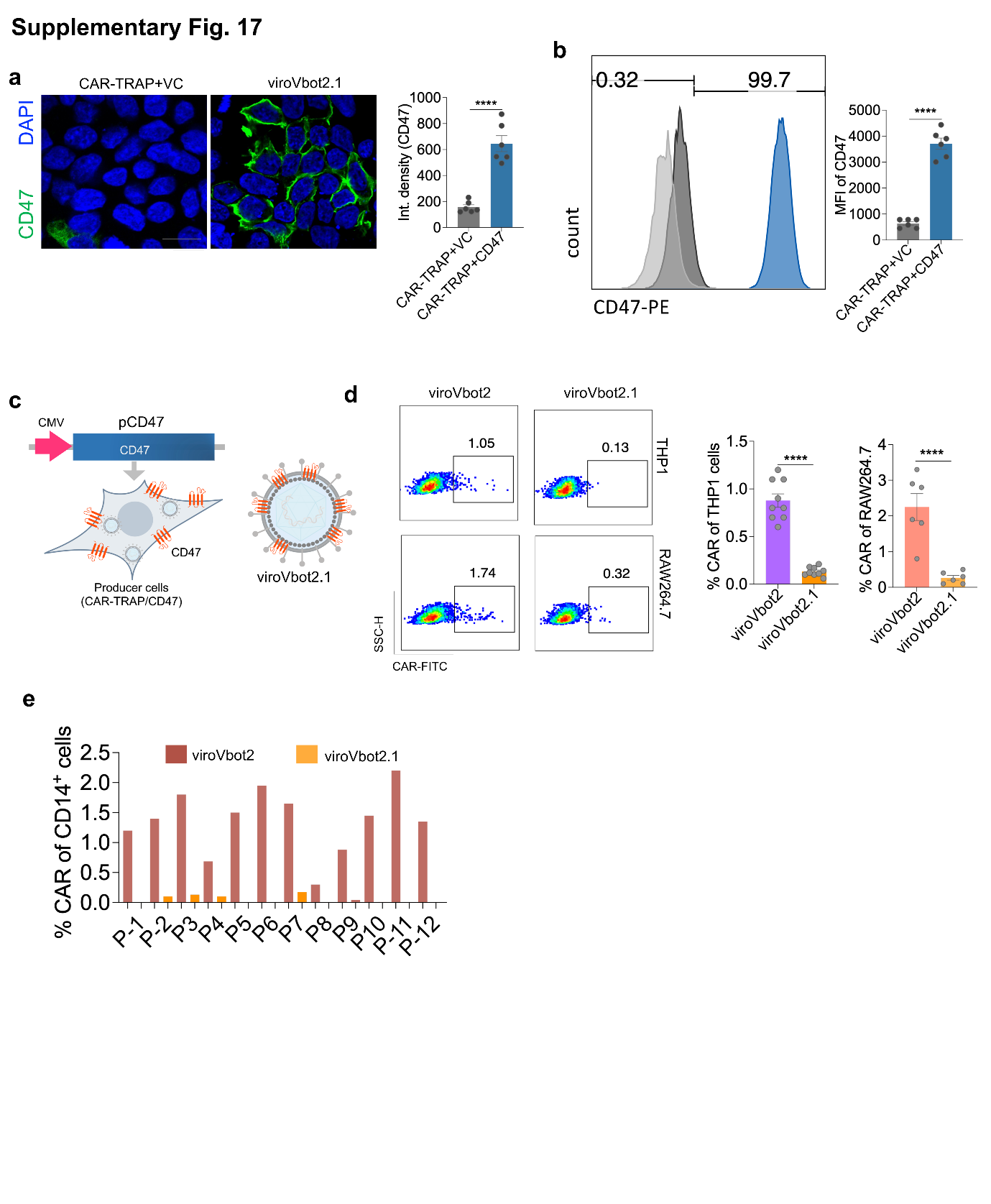

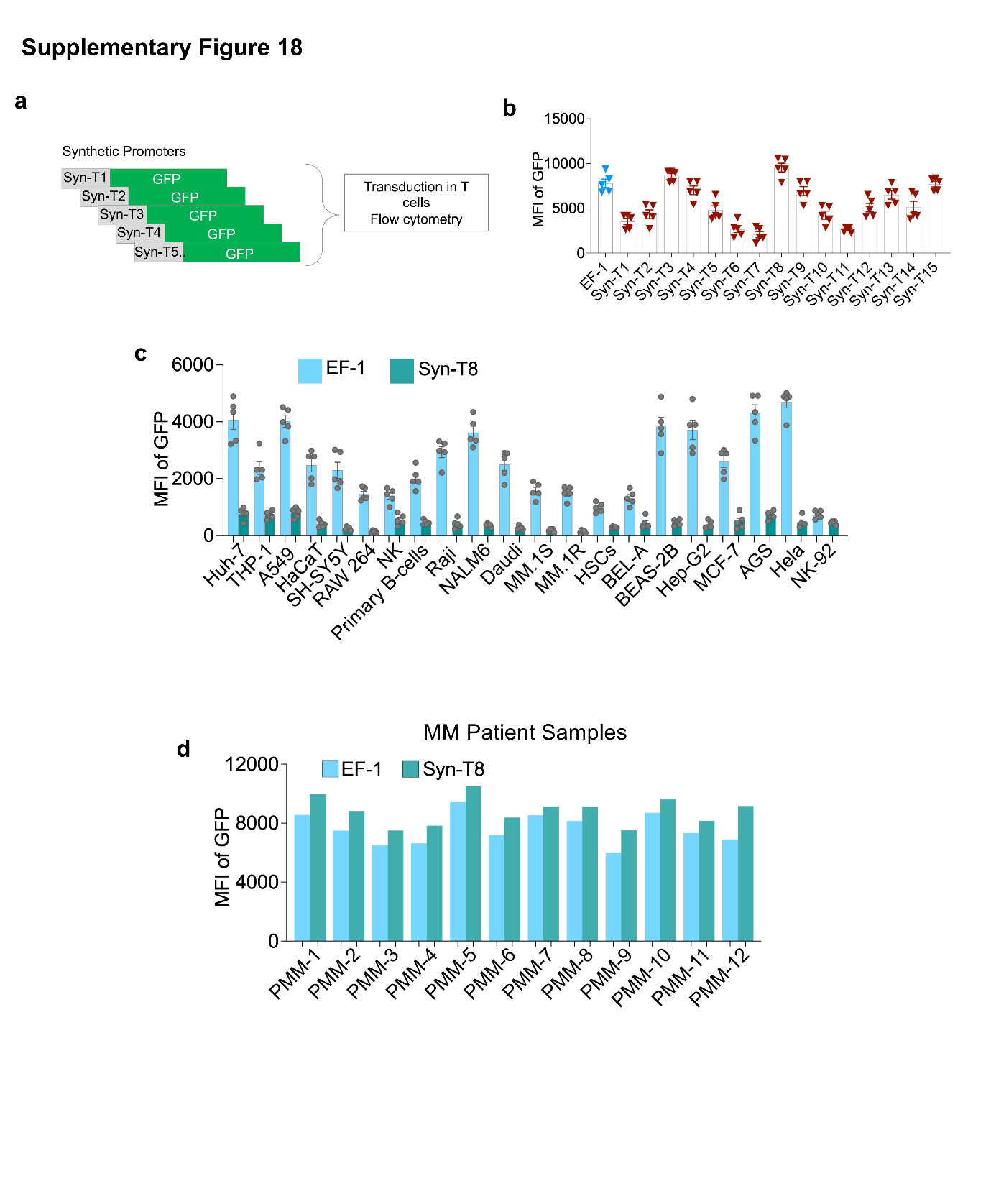

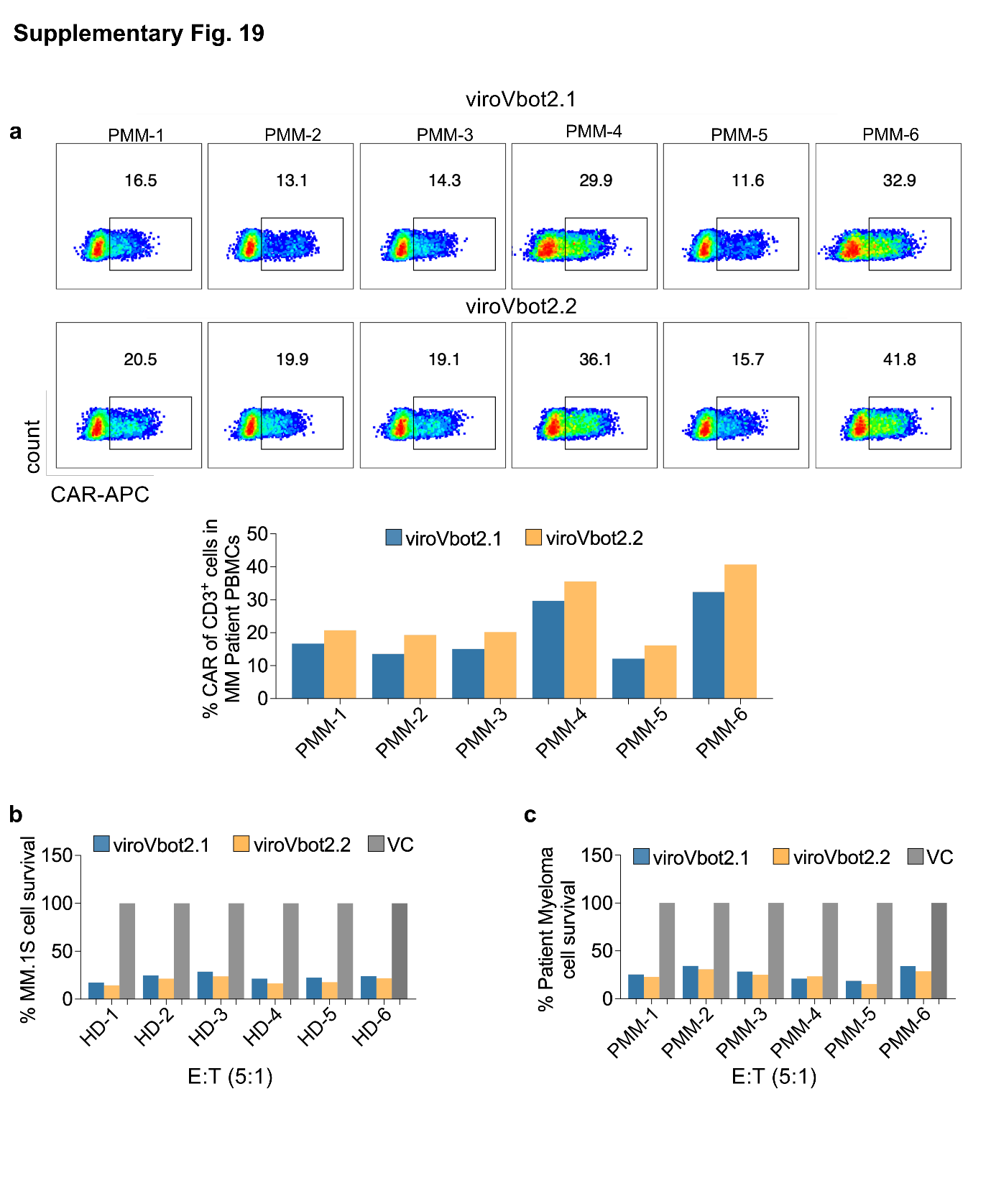

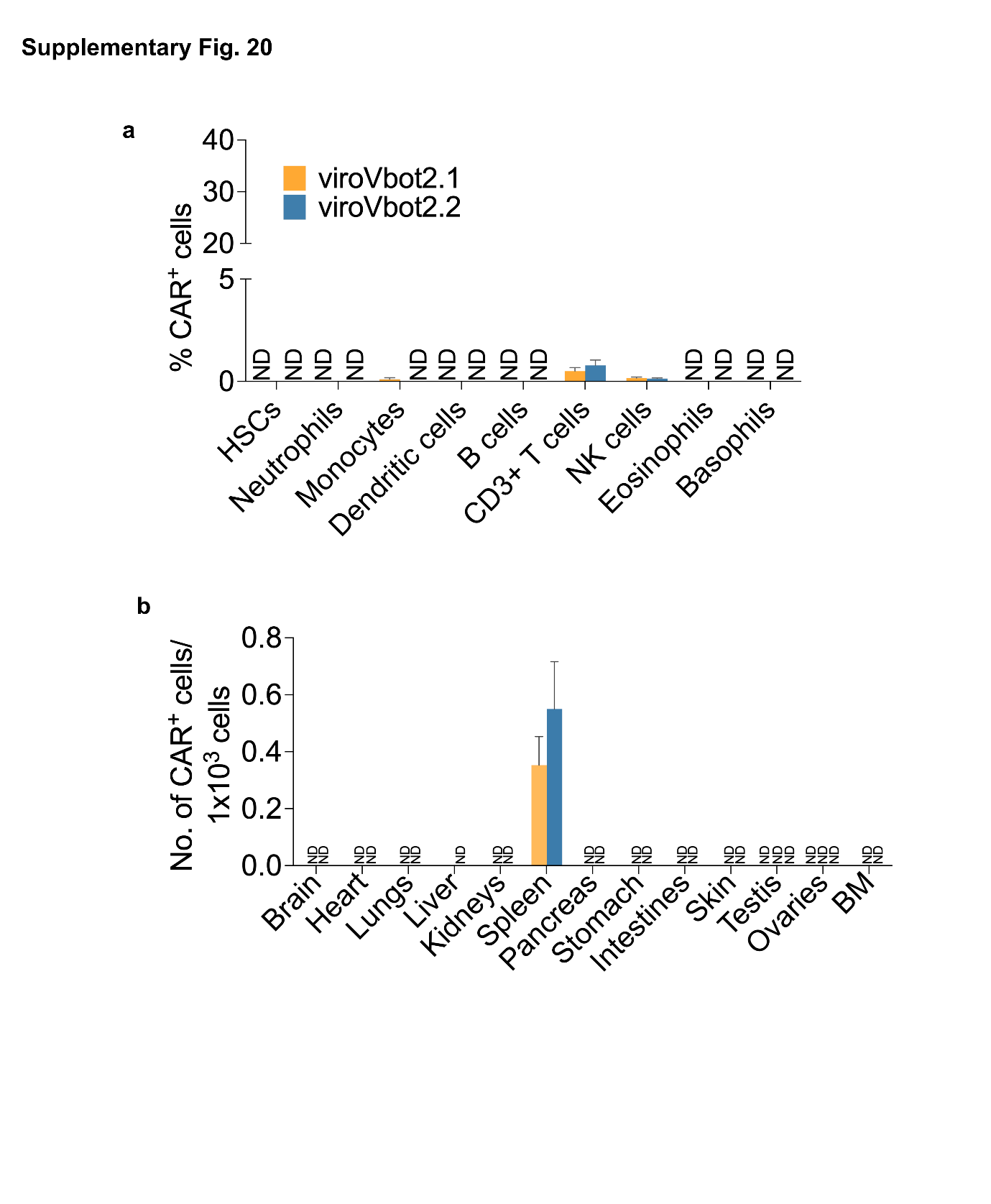

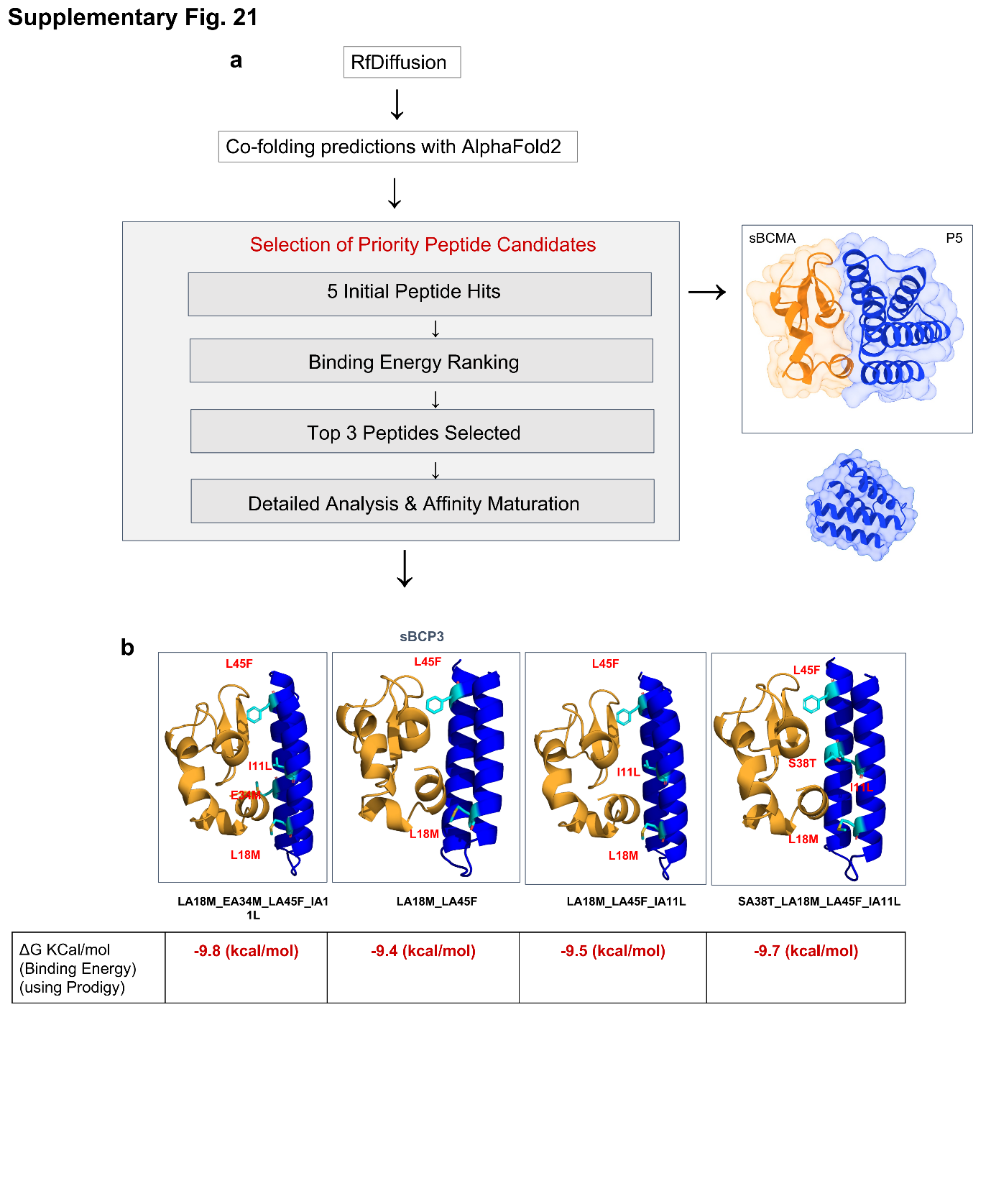

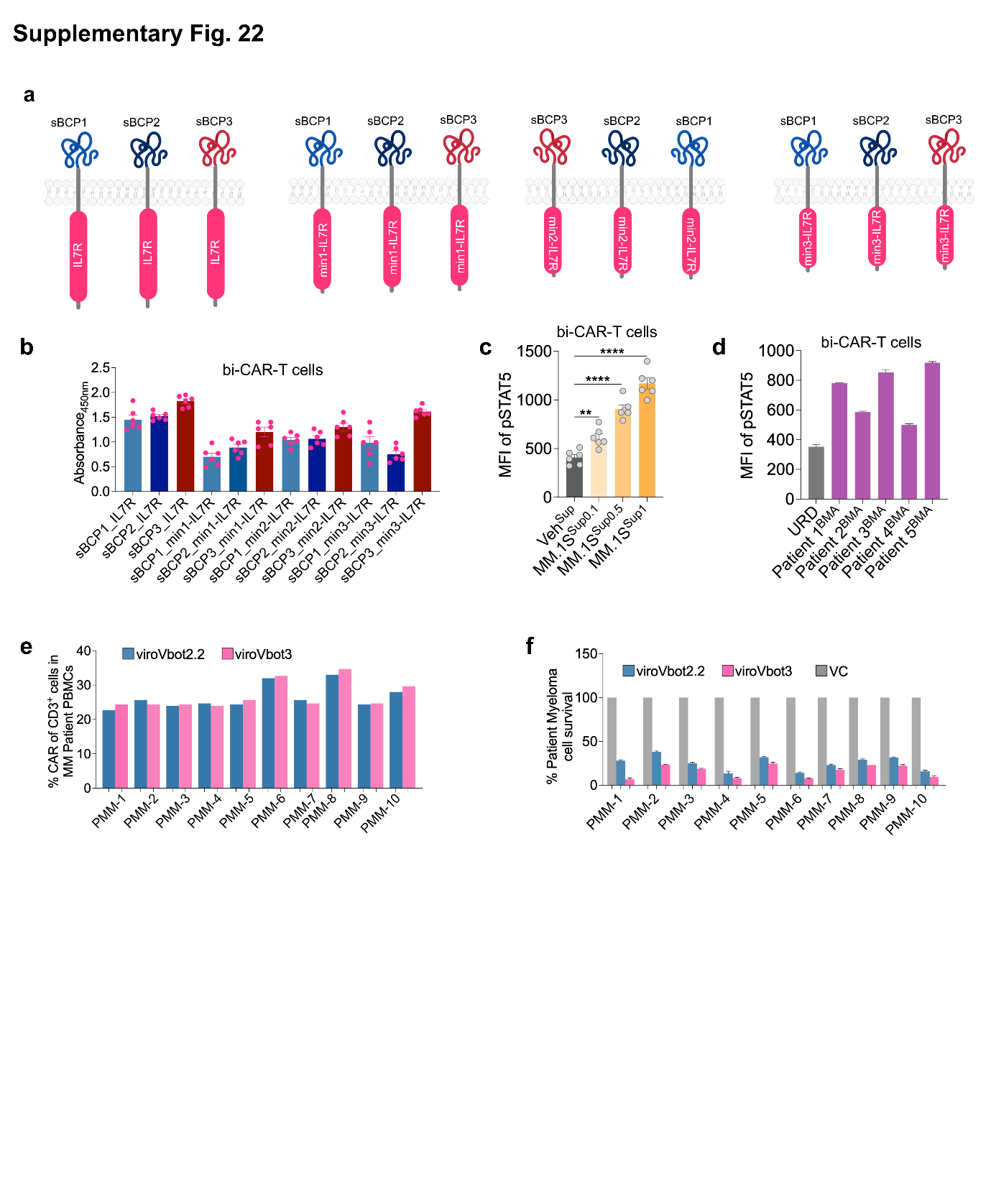

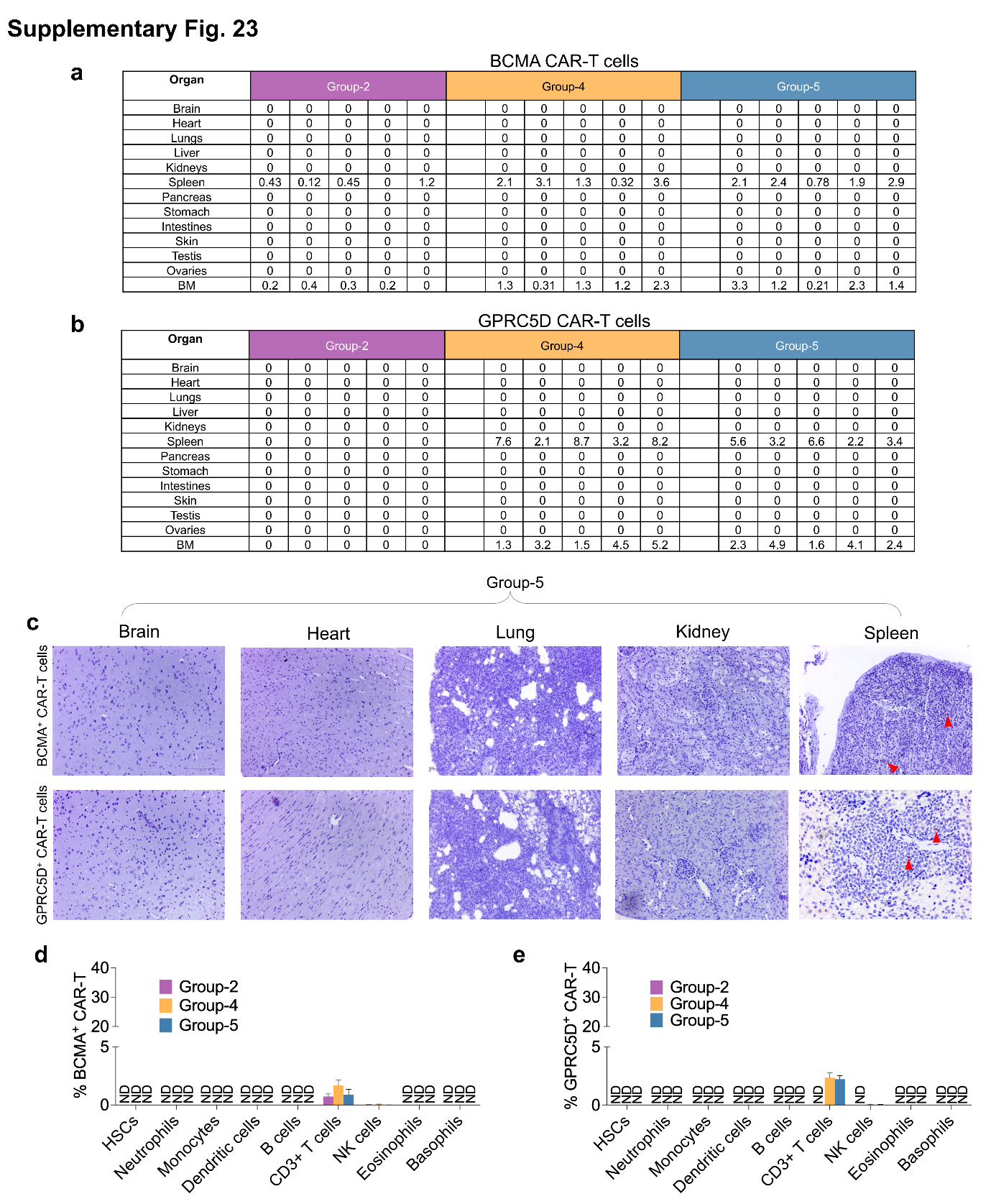

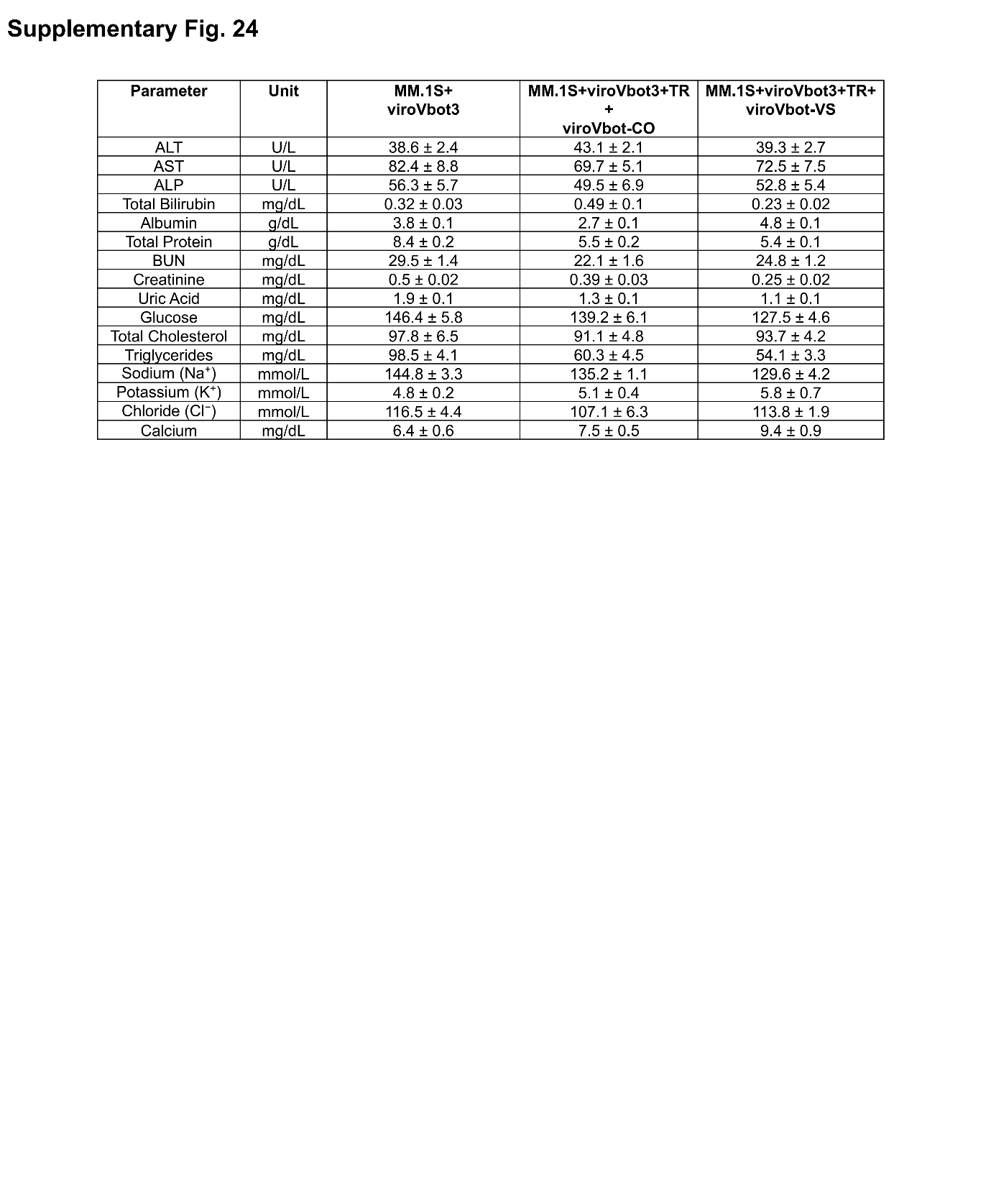

**Supplementary Fig. 1**

**Workflow of the AI pipeline (CIMMEXA) for screening pH-dependent viral envelope proteins**

From 22,562 viral envelope glycoprotein sequences, homology searches identified 641 non-redundant sequences, which were ranked for immunogenicity using the CIMMEXA platform. Following sequential computational prioritization and validation, 12 candidate envelopes were shortlisted for further in vitro evaluation. The in vitro validation lead to the identification of 12 low-immunogenic envelopes that retained lentiviral vector functionality.

**Supplementary Fig. 2**

**MHC-I and MHC-II binding predictions**

**a, Predicted MHC-I strong binders and weak binders across the 12 envelope proteins shortlisted by CIMMEXA. b, Similarly, predicted for MHC-II and c, Combined for MHC-I and MHC-II. Red and gray bars indicate strong binders (SB) and weak binders (WB), respectively.**

**Supplementary Fig. 3**

**T cell immunogenicity profiles across 65 healthy donors.**

**a**, Schematic representation of the workflow for PBMC isolation from healthy volunteer donors followed by flow cytometry analysis. **b**, Heatmaps showing CD154 (CD40L) expression in CD4^+^ T cell subsets, including effector memory (T_em_) and central memory (T_cm_) T cells, compared to an unstimulated vehicle (VEH) control across 65 healthy donors.

**Supplementary Fig. 4**

**Validation of MHC-I knockout HEK293T producer cells and assessment of envelope-specific T cell response**

**a,** Schematic illustration of CRISPR-Cas9-mediated knockout of MHC-I. **b,** Representative confocal microscopy images of MHC-I surface expression (red, Alexa Fluor 647) in CRISPR-generated HEK293T^MHC-I−/−^ cells and wild-type controls (n=7 biologically independent samples). c, Flow cytometry histograms of wild-type and knockout cells with isotype controls, and quantification of mean fluorescence intensity (MFI). **d,** GFP transduction efficiency in knockout versus wild-type producer cells (n=5 biologically independent samples). **e,** Growth kinetics of cells in culture over time. **f,** Quantification of 7-AAD staining assessed by flow cytometry (n=3 biologically independent samples). **g,** Cytotoxicity of virus-specific CD8^+^ T cells from respective envelope-immunized animals against their respective envelope-expressing HEK293T cells and control targets. **h, i,** Cytokine levels (IFN-γ and TNF-α) across VSV-G; VSAV-G; COCV-G; and PIRYV groups. Data is presented as mean ± SEM from three independent biological replicates. Statistical analysis was performed using a non-parametric t-test (*p < 0.05, **p < 0.01, ***p < 0.001, ****p < 0.0001). Scale bar: 50 µm.

**Supplementary Fig. 5**

**Structural comparison of wild-type and immunogenicity mutated envelope proteins**

**a-c,** Ribbon representations of VSV-G wild-type (WT, purple; left), the immunogenicity (Img)
engineered variant (VSV-G_Img_mut, orange; middle), and superimposition of the two structures (right). Structural alignment yielded a root-mean-square deviation (RMSD) of 0.133 Å for VSV-G, 0.689 for COCV-G and 0.176 for VSAV-G, indicating preserved overall protein architecture following Img engineering. Surface representations are shown in grey.

**Supplementary Fig. 6**

**Transduction efficiency of immunogenicity mutated envelope-pseudotyped lentiviral vectors**

**a,** Schematic of the GFP-based transduction assay workflow. **b-e,** Relative transduction efficiency in HEK293T cells for various immunogenicity mutated variants and respective wild type (wt) controls of VSV-G (b), VSAV-G (c), COCV-G (d), and PIRYV (e). GFP fluorescence was quantified 48 h post-transduction. Data is presented as mean ± SEM (n = 4 biologically independent samples).

**Supplementary Fig. 7**

**PIRYV envelope receptor specificity validation**

**a,** Schematic of single and triple-receptor knockout HEK293T cell lines with CRISPR-Cas9 and envelope pseudotyped lentiviral vector transduction in wild-type HEK293T cells (HEK293T^WT^) compared to LDLR knockout (HEK293T^LDLR−/−^), VLDLR knockout (HEK293T^VLDLR−/−^), LRP1 knockout (HEK293T^LRP1−/−^), and triple-receptor knockout (HEK293T^TRN−/−^), transduced with VSV-G; COCV-G; or PIRYV. **b-d,** Relative mRNA expression of LDLR (b), VLDLR (c), and LRP1 (d) in respective knockout cells, transduced with PIRYV and measured by qPCR, with ACTB (β-actin) as the internal control (n=6 biologically independent samples). Data is presented as mean ± SEM.

**Supplementary Fig. 8**

**LDLR binding interface disruption by receptor binding deficient mutations in envelope proteins**

**a-c,** Molecular docking structures of wild-type (wt) and mutant vesiculovirus envelopes complexed with LDLR showing loss of critical binding interactions; receptor binding deficient (RBD). (a) VSV-G (H8/A8, K47/Q47, R354/A354); (b) COCV-G (K47/A47, Y209/A209, E353/A353, R354/A354); and (c) PIRYV (K182/A182, K354/A354, Y207/A207, Y352/A352) are shown with LDLR binding interfaces. Wild-type complexes display hydrogen bonding networks (dashed lines) and salt bridges that are eliminated by mutation, demonstrating specific abrogation of LDLR binding. Stick representations highlight critical residues. Structural analysis explains the RBD phenotype and shows that these mutations directly impair LDLR engagement.

**Supplementary Fig. 9**

**Computational analysis of PIRYV-G de-immunization mutations**

**a,** Ribbon representation of the PIRYV glycoprotein showing distinct structural domains (color-coded). Structural superimposition of PIRYV (magenta) and VSV-G (multicolor) glycoproteins docked onto the LDL receptor, showing overlapping receptor-binding interfaces. **b,** Amino acid sequence of the PIRYV glycoprotein (UniProt: Q85213; GLYCO_PIRYV) with key residues selected for mutagenesis indicated (right). **c,** Molecular docking poses showing predicted binding interactions between PIRYV residues and LDLR contact sites (residue-level views). Summary table of predicted intermolecular contacts between key PIRYV residues (K182, Y207, Y352, K354) and corresponding LDLR residues. **d,** RMSD trajectories and mean RMSD values demonstrating overall structural stability. **e,** MM-GBSA binding free energy (ΔG) calculations. **f,** MM-GBSA binding free energy (ΔG) values calculated from independent simulation replicates for the PIRYV; VSV-G and COCV-G, wild type and mutants. **g,** RMSD and RMSF Plot over 200 ns MD simulations for three independent replicate and MM-GBSA of mutated RBD PIRYV along with final immunogenicity-reduced PIRYV variant (ePIRYV^RBD^).

**Supplementary Fig. 10**

**Molecular dynamics snapshots of PIRYV and LDLR binding dissociation**

**a-c,** All-atom molecular dynamics simulation snapshots (0, 200 ns) comparing wild-type PIRYV with LDLR complex (a) with RBD mutant K182A_Y352A (b) or K182A, Y207A, Y352A, K354A (c). Wild-type complex maintains stable LDLR binding throughout 200 ns trajectory. RBD mutant complex shows progressive loss of binding interface contacts and receptor dissociation by 200 ns. Merged overlay at 200 nanoseconds show structural divergence between stable wild-type and disrupted mutant conformations. Snapshots provide dynamic visualization validating that RBD mutations specifically abrogate LDLR engagement.

**Supplementary Fig. 11**

**Transduction efficiency of PIRYV RBD variants across cell types**

**a,** Summary table of PIRYV RBD mutant variants including wild-type control and targeting critical LDLR binding residues (K182A, Y207A, Y352A, K354A). **b,** Schematic of lentiviral vector transduction workflow showing pseudotyping with PIRYV variants and transduction of target cells. **c-e,** Flow cytometry quantification of GFP-positive cells transduced with PIRYV variant pseudotyped lentiviral vectors: (c) HEK293T cells, (d) Jurkat T cell line, and (e) primary CD3^+^ T cells from healthy donors. Relative transduction efficiency normalized to wild-type PIRYV control (n=6 biologically independent samples). Data is presented as mean ± SEM.

**Supplementary Fig. 12**

**De novo nanobody design using CelNFo AI-driven computational pipeline**

**a,** Workflow of CelNFo pipeline integrating multiple machine learning approaches: antigen and epitope analysis, diffusion-based geometric transformers for optimized nanobody backbone generation, and sequence design using graph neural networks (MPNN). **b,** Schema of de novo designed nanobody candidates targeting CD3, CD4, CD5, CD7, CD8 (lineage markers and activation molecules), T cell receptor alpha (TCRα) with refinement of complementarity-determining regions for high target specificity and reduced immunogenicity. Also, endogenous CD252, CD80 (co-stimulatory molecules) were used. **c,** Immunofluorescence validation of de novo nanobodies and CD252, CD80 in transfected cells showing expression of individual nanobodies (nbCD3, nbCD4, nbCD5, nbCD7, nbCD8) detected via anti-myc antibodies with DAPI nuclear counterstain and for costimulatory molecules (CD80, CD252), anti-CD80 and anti-CD252 antibodies were used (n=6 images in each condition). **d-f,** Bar graph showing flow cytometry analysis of GFP positive CD3 T cells at various MOIs (n=5 biologically independent samples). Data is presented as mean ± SEM.

**Supplementary Fig. 13**

**Conformational stability and binding energetics of the human anti-CD19 and Humanized anti-BCMA antibody-antigen complexes.**

**a, b,** Surface-and-cartoon representations of the CD19-scFv (human scFv) bound to the CD19 ectodomain (CD19R) (grey) at 0 ns (a) and 200 ns (b) of the molecular dynamics (MD) trajectory. Variable light (VL, orange) and variable heavy (VH, blue) domains are shown, with VL CDRs (yellow) and VH CDRs (teal) highlighted at the interface. **c,** Backbone RMSD of the CD19-CAR complex over 200 ns for two independent replicates (Rep1, teal; Rep2, orange). Bar chart (right) shows trajectory-averaged RMSD (Rep1, 4.1 Å; Rep2, 6.0 Å); the higher Rep2 value reflects a conformational transition in the CD19 loop region after ~100 ns. **d,** Per-residue RMSF for both replicates, coloured by chain segment (CD19, light chain, linker, heavy chain); grey bands mark the six CDRs (LCDR1-3, HCDR1-3). **e, f,** Equivalent representations of the humanized (Hz) BCMA-scFv (VL, orange; VH, blue) bound to the BCMA ectodomain (BCMAR) (grey) at 0 ns (e) and 200 ns (f); CDRs coloured as in *a, b*. **g,** Backbone RMSD of the BCMAR-BCMA-scFv complex over 200 ns (mean, 2.4 Å). **h,** Per-residue RMSF for the BCMA complex, coloured by chain segment, with CDRs indicated by grey bands. **i, j,** MM-GBSA binding free energies (ΔG, kcal mol^-1^) for each complex and replicate; more negative values indicate stronger predicted binding.

**Supplementary Fig. 14**

**Nanobody-redirected ePIRYV-pseudotyped LVV generated CAR-T cells and their functional characterisation**

**a,** Flow cytometry analysis of CD3^+^ T cells transduced with ePIRYV^WT^ or ePIRYV^RBD+nbC3/7^ carrying a bi-CAR (BCMA/CD19) transgene. Histograms show transduction efficiency based on BCMA-CAR, CD19-CAR, or anti-G4S antibody detection, with corresponding quantitative analysis (n = 6 biologically independent samples). **b,** Percentage of CAR-expressing CD4^+^ and CD8^+^ T cells following activation with CD3/CD28 Dynabeads or vehicle control (PBS) (n = 6 biologically independent samples). **c,** Schematic of the pseudovirus transduction assay. Producer cells were co-transfected with packaging plasmids, a bi-CAR construct, and the ePIRYV envelope plasmid. Harvested pseudoviral particles were used to transduce target cells, and transduction efficiency was quantified by CAR expression in CD3^+^ T cells. **d,** Percentage CAR transduction in patient-derived CD3^+^ T cells, measured using an anti-G4S linker antibody. **e, f,** Dose-dependent transduction of ePIRYV^WT^ and ePIRYV^RBD+nbC3/7^ pseudotyped vectors across target cell lines at different effector-to-target (E:T) ratios (n = 5 biologically independent samples). **g-i,** Flow cytometric quantification of Granzyme B, IFN-γ, and IL-2 expression (n = 6 biologically independent samples). **j,** Pie charts and bar graph showing the distribution of CD8^+^ CAR-T cell subsets in the two experimental groups (n = 5 biologically independent samples). Data is presented as mean ± SEM. Statistical analysis was performed using a non-parametric t-test (****p < 0.0001; ns: not significant).

**Supplementary Fig. 15**

**Histological analysis of CAR-T cell infiltration and organ safety**

**a,** Bar graph showing Immunohistochemical detection of CAR^+^ cells with tissue infiltration across multiple organs. **b,** Bar graph of assessment of CAR vector biodistribution by quantification of vector copy number (VCN) in various tissues using PCR-based analysis. Data represent n = 5 mice per group, with multiple tissue sections analyzed per organ.

**Supplementary Fig. 16**

**miRNA-122 mediated silencing of bi-CAR in hepatocytes.**

**a,** Schematic illustration of lentiviral constructs encoding a bi-CAR (BCMA/CD19): a non-targeting control (NTC; top) and five tandem 5xmiR-122T target sites evaluated for expression in CAR-TRAP producer cells. **b,** Representative flow cytometry plots of CAR expression in primary human hepatocytes (PHH) transduced with bi-CAR expressing NTC or 5xmiR-122T (viroVbot2) (n=6 biologically independent samples). **c,** Representative flow plots of percentage CAR expression in CD3^+^ T cells transduced with NTC or mir-122 (viroVbot2) along with the quantative analysis (n=6 biologically independent samples). **d,** Anti-tumor activity against MM.1S cells at various E:T ratios (n=5 biologically independent samples). Data is presented as mean ± SEM. Statistical analysis was performed using a non-parametric t-test (****p < 0.0001).

**Supplementary Fig. 17**

**Functional characterization of CD47-engineered viroVbot2.1**

**a,** Representative immunofluorescence images showing surface localization of CD47 in CAR-TRAP producer cells, with DAPI nuclear counterstaining, compared to vector control (VC), along with quantitative analysis using Fiji/ImageJ (n = 6 images). **b,** Flow cytometry histograms showing CD47 expression in producer cells versus CAR-TRAP + VC control (CD47-PE), along with quantitative analysis of mean fluorescence intensity (MFI) (n = 6 biologically independent samples). **c,** Schematic representation of the CD47 lentiviral vector, its transduction into HEK293T CAR-TRAP producer cells, and the generation of viroVbot2.1. **d,** Comparative CD47-PE flow cytometry histograms and bar graphs of macrophage/monocyte cell lines (THP-1 and RAW264.7), comparing viroVbot2 and viroVbot2.1 vector (n = 9 and 6 biologically independent samples). **e,** Quantitative analysis of flow cytometry data showing CAR expression in CD14^+^ cells following transduction with the respective viroVbots in PBMCs obtained from different patient samples. Data is presented as mean ± SEM. Statistical analysis was performed using a non-parametric t-test (****p < 0.0001). Scale bar: 50 µm.

**Supplementary Fig. 18**

**PromoterForge synthetic promoter validation**

**a,** Schematic representation of synthetic promoter (Syn-T8) designs optimized for T cell-specific expression, incorporating integrated transcriptional response elements. **b,** Bar graph showing mean fluorescence intensity (MFI) from flow cytometry analysis of GFP expression in primary CD3^+^ T cells transduced with vectors containing either conventional (EF-1) or synthetic promoters. Cells were transduced with a GFP reporter under different promoter configurations, with EF-1 serving as a control (n = 5 biologically independent samples). **c,** GFP expression analysis across multiple cell lines comparing EF-1 and synthetic promoter (SynT-8)-driven constructs (n = 5 biologically independent samples). **d,** GFP expression in multiple myeloma patient–derived T cells, demonstrating consistent Syn-T8 promoter performance across patient samples (n = 5 biologically independent samples). Data are presented as mean ± SEM.

**Supplementary Fig. 19**

**Syn-T8 synthetic promoter shows robust CAR expression and anti-tumor activity in healthy and patient-derived T cells**

**a,** Representative flow cytometry plots showing CAR-APC expression in multiple myeloma patient-derived T cells transduced with vectors driven by the Syn-T8 promoter, along with quantitative analysis of the flow cytometry data. **b,** Bar graph showing the anti-tumor activity of healthy donor (HD)-derived T cells against MM.1S target cells. **c,** Bar graph showing the anti-tumor activity of patient-derived T cells against autologous multiple myeloma cells from the same patients. Data are presented as mean ± SEM.

**Supplementary Fig. 20**

**Biodistribution and tissue localization of CAR-expressing cells following In Vivo delivery**

**a,** Bar graph showing the percentage of CAR-positive cells, detected using an anti-G4S antibody, across different blood cell populations. Quantification was performed based on flow cytometry analysis (n = 5 biologically independent samples). **b,** Immunohistochemical analysis of CAR-positive cells in various tissues using a BCMA-CAR specific detection antibody. Representative tissue sections and corresponding quantitative analyses demonstrate the distribution and localization of CAR-expressing cells across the examined organs. (n = 5 biologically independent samples). Data are presented as mean ± SEM.

**Supplementary Fig. 21**

**Computational design and affinity maturation of peptide binders targeting sBCMA**

**a,** Peptide binders to soluble BCMA (sBCMA) were designed de novo using RFdiffusion, and the resulting peptide-sBCMA complexes were validated by co-folding predictions with AlphaFold2. Five initial hits were ranked by predicted binding energy, and the top three candidates were taken forward to detailed interface analysis and affinity maturation, yielding the lead designs sBCP1-3. **b,** For the sBCP series, interface residues were iteratively mutated. For example, the residues shown for sBP3 (L18M, L45F, I11L, E34M, S38T) and each variant was re-modeled in complex with sBCMA. Binding free energies (ΔG) were computed using PRODIGY, ranging from -9.4 to -9.8 kcal/mol. The mutant LA18M_LA45F showed the optimal predicted affinity (-9.4 kcal/mol) and better stability, and was selected as the optimized binder. Similar analysis was done for sBP1 and sBP2. This figure is supported by the detailed analysis in **Supplementary Information 2.**

**Supplementary Fig. 22**

**sBCMA-IL7R chimeric receptor optimization showing pSTAT5 activation**

**a,** Schematic of chimeric receptor designs combining sBCMA binder variants (sBC-P1, sBC-P2, sBC-P3) with IL-7R intracellular domains (full-length or minimal). **b**, STAT5 phosphorylation ELISA assay (in CAR-T cell total cell protein) showing dose-dependent activation to soluble BCMA for sBC-P3_min3-IL7R. **c,** Flow cytometry analysis of pSTAT5 shown as MFI in response to the supernatant obtained either as control DMEM media or supernatant (Sup) collected from MM.1S cells and incubated with CAR-T cells (n=6 biologically independent samples). **d,** Similarly, bone marrow aspirates (BMA) from five myeloma patients versus unrelated donor (URD). **e,** Flow cytometry analysis of % CAR of CD3^+^ T cells in PBMCs from ten multiple myeloma patients following transduction with viroVbot2.2 and viroVbot3. **f,** Bar graph depicting the cytotoxicity (% patient myeloma cell survival) mediated by viroVbot2.2-, viroVbot3- and vector control (VC)- treated CAR T cells across the same patient samples. sBC-P3-min1-IL7R selected as optimized variant for viroVbot3 integration. Data is presented as mean ± SEM.

**Supplementary Fig. 23**

**Organ biodistribution and immune cell infiltration by dual-targeted CAR-T cells**

**a, b,** Table showing biodistribution of BCMA (a) and GPRC5D (b) CAR-T cells across organs in mice treated with MM.1S+viroVbot3 (Group 2); MM.1S+viroVbot3+TR+viroVbot-CO (Group-4); or MM.1S+viroVbot3+TR+viroVbot-VS (Group-5) (n=5 mice each group). Values represent CAR-T cell counts per organ. **c,** Representative immunohistochemistry (IHC)-stained sections of major organs from BCMA and GPRC5D CAR-T cell-treated mice as a representation from Group-5. Red asterisks indicate CAR-T cell infiltration. **d, e,** Flow cytometry analysis quantifying BCMA and GPRC5D positive CAR-T cell infiltration into immune and myeloid subsets across treatment groups (n=5 mice each group). Data are presented as mean ± SEM.

**Supplementary Fig. 24**

**Biochemical safety profile of viroVbot3**

Serum biochemical parameters including hepatic markers (ALT, AST, bilirubin), renal function (creatinine, BUN), complete blood count, inflammatory cytokines (IL-2, IFN-γ, TNF-α, IL-6), and coagulation markers measured at baseline and days 7 to 28 post-viroVbot3 administration. Data is presented as mean ± SEM (n = 5 mice per group). No hepatotoxicity or nephrotoxicity detected; transient cytokine elevation consistent with CAR-T activation demonstrates favorable safety profile.
